# Convergent Hybrid Phase Ligation Strategy for Efficient Total Synthesis of Large Proteins Demonstrated for 212-residue Linker Histone H1.2

**DOI:** 10.1101/661744

**Authors:** Ziyong Z. Hong, Ruixuan R. Yu, Xiaoyu Zhang, Allison M. Webb, Nathaniel L. Burge, Michael G. Poirier, Jennifer J. Ottesen

**Author notes:** Correspondence: Jennifer J. Ottesen.

## Abstract

Simple and efficient total chemical synthesis of large proteins remains a significant challenge. Here, we report development of a convergent hybrid phase native chemical ligation (CHP-NCL) strategy that should be generally applicable for facile preparation of large proteins. Key to the strategy is the use of sequential ligation on the solid phase for the directed assembly of ~100-residue segments from short, synthetically accessible peptide components. These segments can then be assembled via convergent solution phase ligation, exploiting o-aminoaniline as a chemically flexible cryptic thioester with multiple activation modalitiies on resin and *in situ*. We demonstrate the feasibility of our approach through the total synthesis of 212-residue linker histone H1.2 in unmodified, phosphorylated, and citrullinated forms, each from eight component peptide segments. We further demonstrate that fully synthetic H1.2 replicates the binding interactions of linker histones to intact mononucleosomes, as a proxy for the essential function of linker histones in the formation and regulation of higher order chromatin structure.

---

The ability to generate homogenous samples of proteins with specific defined modifications remains a limiting step in understanding the structure, dynamic, and functional outcomes of protein post-translational modification (Fierz & Muir, 2012). Of the many strategies that have been developed in the past decades (Chuh, Batt, & Pratt, 2016; Dadová, Galan, & Davis, 2018; Howard, Yu, Gardner, Shimko, & Ottesen, 2015; Luger et al., 2018; Simon et al., 2007), total protein synthesis has the highest potential for combination of multiple modifications throughout a protein sequence, but also represents a significant effort. Typically, this has required division of a protein into accessible segments that can be stitched together using chemoligation strategies such as native chemical ligation (Agouridas et al., 2019; Dawson, Muir, & Kent, 1994; Kent, 2009; Müller & Muir, 2015) (NCL, resulting in a native peptide bond, most commonly at a cysteine site) followed by desulfurization (allowing expanded ligation sites, including alanine) (Jin, Li, Chow, Liu, & Li, 2017; Wan & Danishefsky, 2007; Yan & Dawson, 2001). Sequential (Raibaut, Ollivier, & Melnyk, 2012; Shimko, North, Bruns, Poirier, & Ottesen, 2011), one-pot (Bang & Kent, 2004; Kamo, Hayashi, & Okamoto, 2018; Li et al., 2014; Tang et al., 2015; Zuo, Zhang, Yan, & Zheng, 2018), and convergent (Bang, Pentelute, & Kent, 2006; Durek, Torbeev, & Kent, 2007; Fang, Wang, & Liu, 2012; Qi, He, Ai, Guo, & Li, 2017) ligation strategies have enabled the synthesis of small to medium-sized proteins (<150 residues) but have been limited in scope and practical application by the size and synthetic accessibility of the component peptide segments, the chemical or kinetic control of intermediate ligation specificity, and the number of purification steps required to achieve the final product. In theory, sequential ligation carried out on the solid phase should overcome these barriers, allowing facile recombination of peptide segments while eliminating the need for intermediate purification (Brik, Keinan, & Dawson, 2000; Canne et al., 1999; Jbara, Seenaiah, & Brik, 2014). However, in practice, this has proven challenging – solid phase ligation approaches have been successfully demonstrated only for small proteins.

The largest protein or segment prepared by entirely solid phase ligation as documented in the Protein Chemical Synthesis Database (Agouridas, El Mahdi, Cargoët, & Melnyk, 2017) is the 125-residue histone H2B (Jbara et al., 2014). Recently, our laboratory demonstrated that for two proteins (104-residue core histone H4 and 140-residue Cenp-A), solid phase ligation strategies could provide significant improvements in yield and purity in initial ligation steps, but yields were significantly reduced after 75-100 residues were assembled in three ligations (Yu et al., 2016). For the two proteins in question, we achieved our product by carrying out additional sequential solution phase ligation steps for a hybrid phase ligation approach. A similar approach was used by Melynk and coworkers for preparation of the 180-residue NK1-B, from an 83-residue segment prepared by solid phase ligation, and two ~50-residue synthetic segments ligated in solution (Ollivier et al., 2017). However, these hybrid approaches are inherently limited again by synthetic accessibility of the additional segments prepared by traditional approaches.

To explore the limitations of the solid phase and hybrid approaches, we selected the 212-residue linker histone H1.2. Variants of linker histone H1 facilitate chromatin condensation by interactions with folded nucleosome core particles (Fyodorov, Zhou, Skoultchi, & Bai, 2018). They are known to be heavily post-translationally modified (Izzo & Schneider, 2016), but the functions of the modifications are not well understood, in part because of the dearth of chemical tools for preparation of the modified proteins. As such, it is a high value target for synthetic protein chemistry but challenging to achieve due to its large size.

Initially, we attempted preparation of histone H1.2 through a simple solid phase ligation strategy from nine component peptides. However, as found in the literature studies described above, we observed significant yield loss on the solid phase after 4 ligations (~100 residues, see Supplemental S5.7).

We hypothesized that we could overcome this limitation by dividing the sequence into two “blocks”, each assembled by ligation of short, synthetically accessible peptide segments on the solid phase. Key to this strategy is use of a cryptic thioester in the N-terminal block that would be orthogonal to all required ligation chemistry, but could be selectively activated in situ for use in a convergent solution phase ligation. We identified peptide *o*-aminoanilides, incorporated through synthesis on the diaminobenzoic acid (Dbz) linker, as a reasonable fit for these requirements (Blanco-Canosa & Dawson, 2008; Blanco-Canosa, Nardone, Albericio, & Dawson, 2015; Mahto, Howard, Shimko, & Ottesen, 2011; Wang et al., 2015). This moiety survives peptide ligation, desulfurization, and cleavage conditions, but can be converted to thioester via a triazole intermediate in aqueous conditions [Scheme 1] (Wang et al., 2015). In addition, since residues C-terminal to the Dbz are eliminated in conversion to the active thioester, these can be exploited to anchor the peptide segment to resin. Here, we selected a reverse NCL anchoring strategy, in which a Cys incorporated through isopeptide linkage to a Lys side chain (**11**) was exploited for ligation to a thioester resin (**12**) to immobilize our peptide segment.

Using this strategy, we developed a ligation-desulfurization scheme such that native H1.2 could be prepared from ligation of 8 synthetically accessible peptide fragments [Scheme 2]. We identified 7 Ala residues at sites with reasonable predicted ligation kinetics (Haase, Rohde, & Seitz, 2008; Johnson & Kent, 2006) spaced such that each segment was < 36 residues in length. These were grouped into two segments of four fragments each. The anchor fragment for the N-terminal segment (**11**) would be synthesized with the internal aminoanilide and C-terminal isopeptide ligation handle for immobilization on the solid phase. The anchor segment for the C-terminal segment (**18**) was synthesized with an internal HMBA linker to allow generation of the native protein C-terminus after ligation (Harris & Brimble, 2010; Hossain et al., 2009; Yu et al., 2016).

Each individual peptide was synthesized with an N-terminal thiazolidine (Thz) protection strategy for the Cys, which is commonly used for chemical control of ligation (Bang & Kent, 2004). Interestingly, we found that use of Dbz as cryptic thioester required significant finesse, as no single approach was suitable for all peptides (full discussion in Supplemental S4). For all but the N-terminal anchoring peptide, N-methyl (**1**), N-alloc (**3**), or unprotected (**2**) diaminobenzoic acid strategies were used for the generation of the peptide thioester (**10**) (Blanco-Canosa & Dawson, 2008; Blanco-Canosa et al., 2015; Mahto et al., 2011; Wang et al., 2015). In brief, we found that while N-methyldiaminobenzoic acid is compatible with microwave peptide synthesis, it 1) adds complexity to the characterization of the peptide-MeDbz intermediate due to formation of the benzimidazole derivative under acidic cleavage or HPLC conditions (Bird, Silvestri, & Dawson, 2018), and 2) often shows reduced yield of the N-acylurea derivative, mostly likely due to reduced reactivity of the N-substituted amine. The N-(alloc)diaminobenzoic acid derivative is compatible with microwave synthesis only if piperidine treatment for Fmoc removal is carried out at room temperature; heated deprotection cycles resulted in premature N-acylurea formation and peptide release from the resin (Tsuda, Uemura, Mochizuki, Nishio, & Yoshiya, 2017). However, we observe minimal product loss (~1% loss per coupling cycle) when carrying out coupling steps using microwave heating.

With all peptides in hand, we proceeded to assess the solid phase ligation for the two major protein blocks. To generate the N-terminal block **26**, we first prepared thioester resin **12**. Rink amide PEGA resin was treated with diglycolic anhydride. This was further treated with thiophenol in presence of DIC and DMAP to obtain resin thiophenol thioester **12**. Anchor peptide **11** was added, and native chemical ligation afforded the linked product. A key step was a capping step in which the resin was treated with an excess of Cys in ligation buffer to deactivate any remaining thioester resin, to avoid potential crossreactivity in subsequent ligation steps.

Treatment with methoxylamine afforded the “ring-opened” Cys product **13** suitable for use in subsequent ligation cycles **(14-16)**. For documentation of these initial syntheses, ligation progress was also monitored by RP-HPLC (Fig. 3) and SDS-PAGE (see ESI); however, these analyses consume significantly more material than mass spectrometric analysis, and are less sensitive than MALDI-TOF for detection of unreacted material. For preparative purposes, we recommend eliminating these steps and restricting analysis to MALDI-TOF MS. In all, we carried out 4 replicated assembly of the N-terminal segment **16** to assess the robustness of the approach; one with unmodified segments, and three with citrullinated Arg53 to generate **16cit**. Typically, desulfurization is carried out as the last step in a convergent ligation scheme. However, desulfurization is complicated with large numbers of thiols in a protein (c.f. **27**). We noted that the thiols in the N-terminal segment **16** are not required for subsequent ligation steps. We therefore explored desulfurization on the solid phase, which we hypothesized could simplify the final desulfurization step (**27** to **28**). Across the four preparations, yields after four ligation/desulfurization cycles ranged from 50-72% by weight, in sufficient purity to use directly in solution phase ligations.

**Figure 1.**
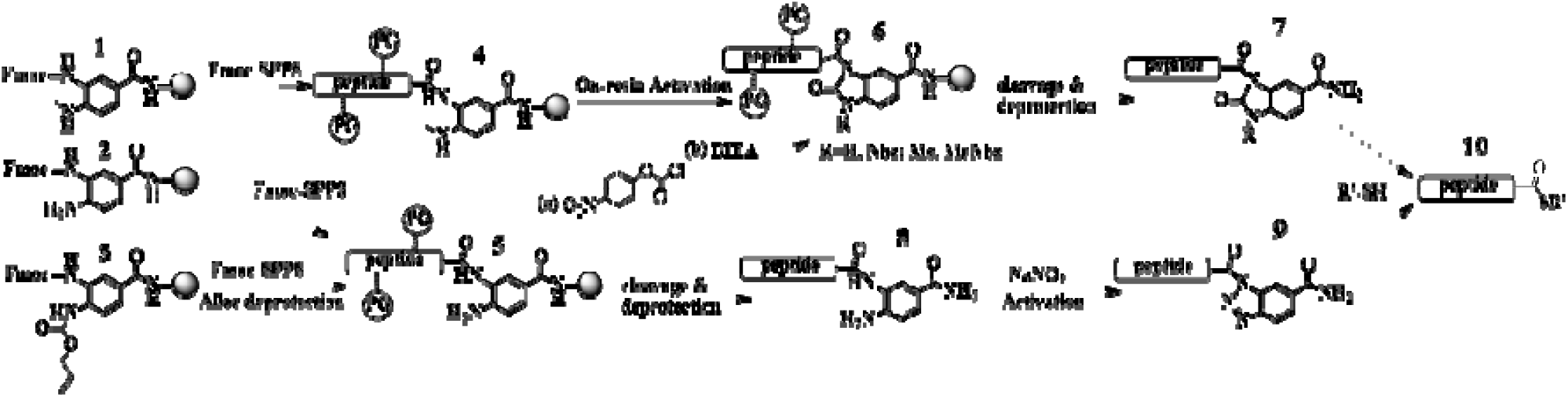
Different modalities for diaminobenzoic acid linkers as cryptic thioesters in peptide synthesis and ligation. Decision points for synthesis include the protection scheme for the linker (**1-3**) and the activation approach (**7,9**), all of which can result in generation of an active thioester **10**. In general, on-resin generation of the N-acylurea derivative **7** is most suitable for generation of standard peptide components. However, in situ generation of the triazole **9** in aqueous solution can be exploited in our convergent ligation scheme.

**Figure 2.**
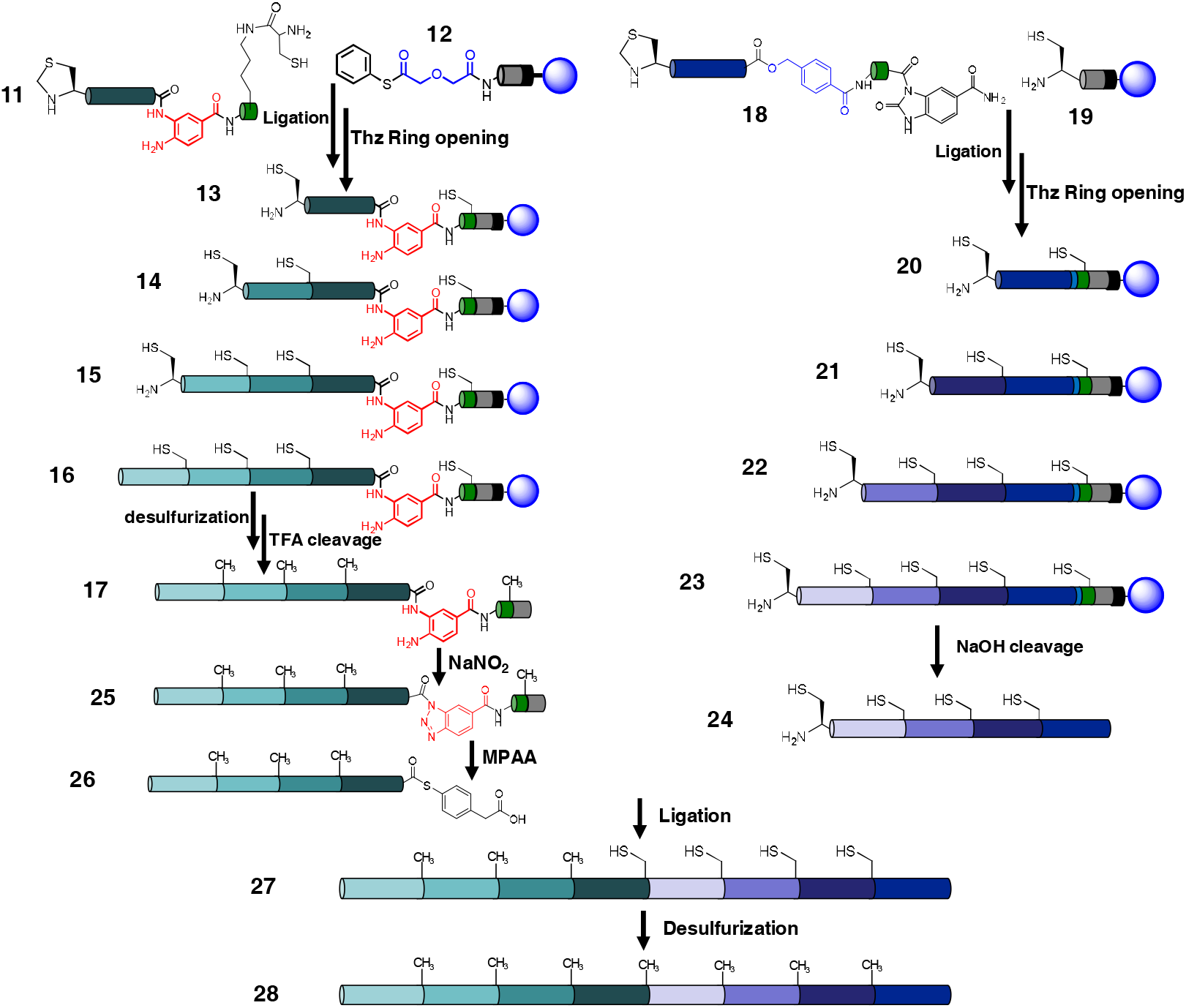
Convergent hybrid ligation scheme for preparation of linker histone H1.2.

**Figure 3.**
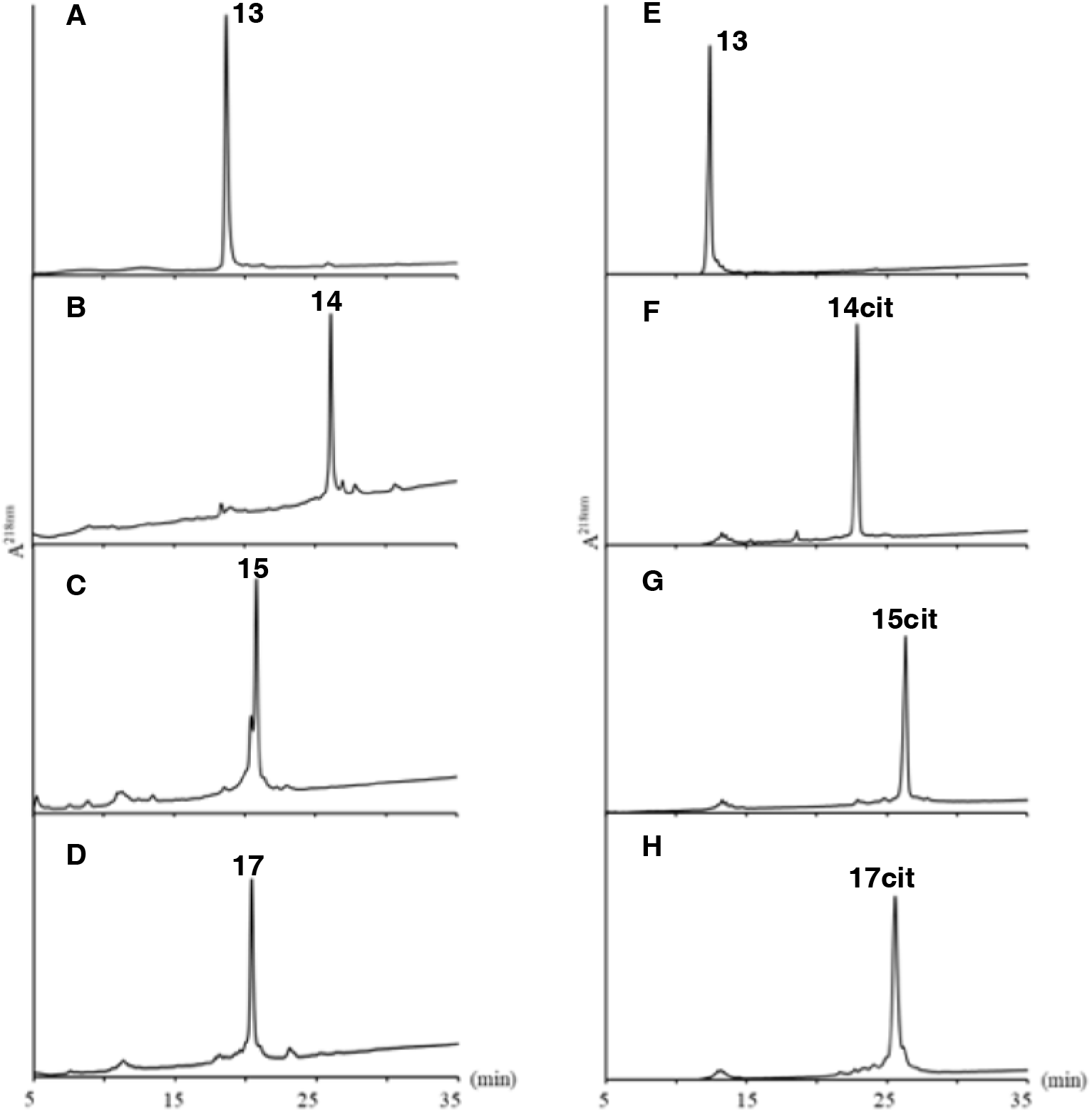
Analytical RP-HPLC analysis of crude products for representative ligations on the solid support for each intermediate product required to generate unmodified N-terminal segment **17** and citrullinated **17cit**. Each product is cleaved at the Rink linker and includes the Dbz linker required for activation to generate thioester **26** or **26cit**. All peptide identities were confirmed by MALDI-TOF MS (see supplemental information). Of note, the side peak in panel C was identified as a thiol adduct that was removed by treatment with reducing agent. Products were obtained in 72% (D) and 50% (H) yields by weight.

For preparation of the C-terminal segment, we followed the protocols developed for the synthesis of H4 and CENP-A (Yu et al., 2016). PEGA-PL resin was used for solid support. Fmoc-Thz-Ala-Ahx-Tyr-Lys-Gly handle sequence (sufficient for detection by RP-HPLC and MALDI-TOF MS) was introduced on the rink amide PEGA resin. Immediately prior to the first ligation, the Fmoc was removed by treatment with 20% piperidine in DMF. Treatment with methoxylamine then afforded the N-terminal cysteine, to enable the attachment of H1-(Thz_189_-Lys_212_)-HMBA-RG-MeNbz/Nbz through native chemical ligation. After ligation, resin was washed and the cycle was repeated for the remaining rounds of SP-NCL. To assess the robustness of our approach, we repeated this synthesis 3 times for unmodified peptide segments. Across the three syntheses, the yields ranged from 77-81% by weight, and the crude product was typically sufficiently pure to be used without purification for the final solution phase ligation (Fig. 4-D).

**Figure 4.**
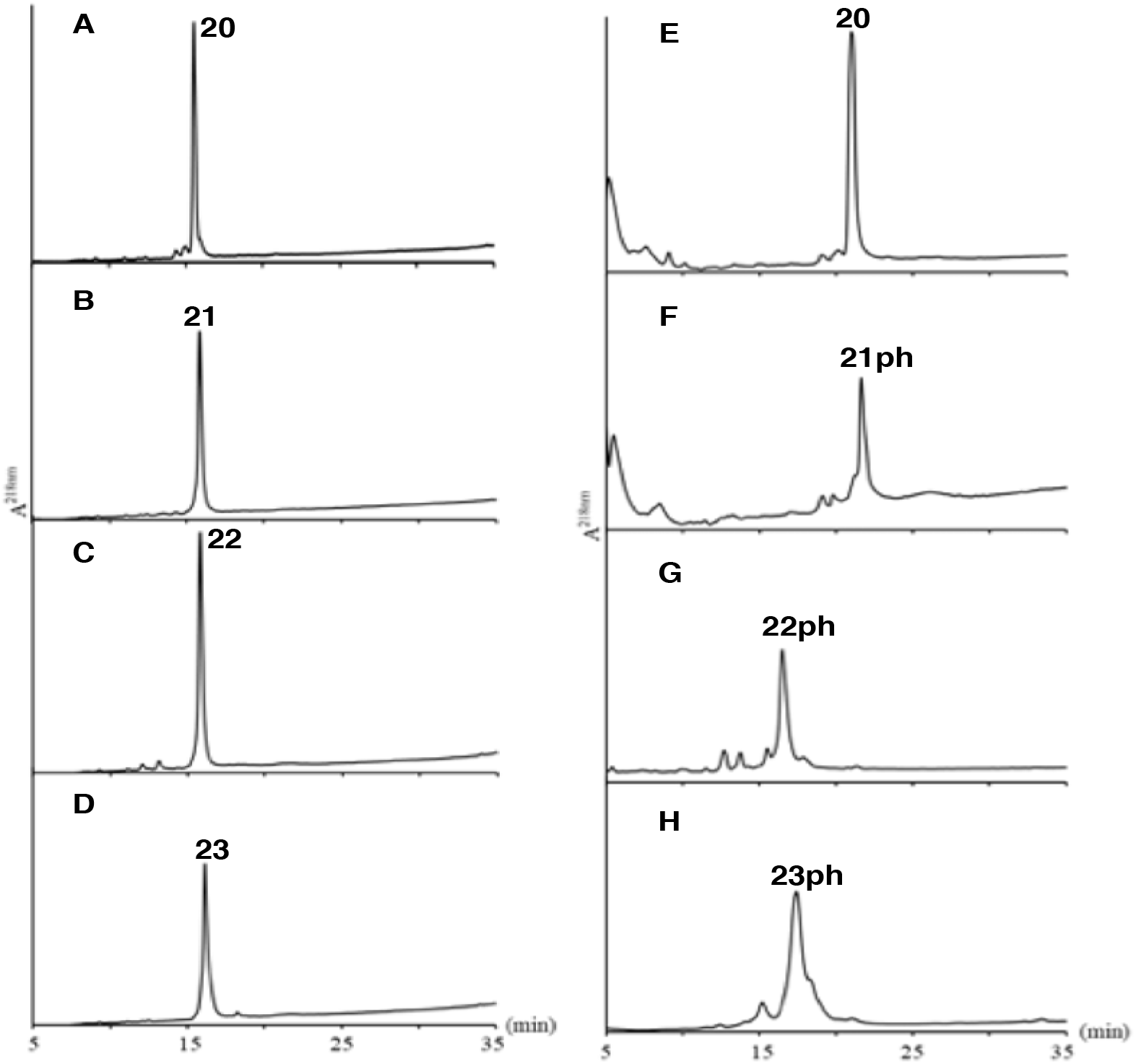
Analytical RP-HPLC analysis of crude products for representative ligations on the solid support for each intermediate product required to generate unmodified C-terminal segment **23** and phosphorylated **23ph**. Each product is cleaved at the HMBA linker (light blue in Fig. 2). All product identities were confirmed by MALDI-TOF MS (see supplemental information). Ligation cycles to generate E-H assessed an alternate deprotection scheme that was less successful. Products were obtained in 77% (D) and 73% (H) yields by weight.

We then prepared C-terminal segment with Ser172ph as described above. Of note, this modified variant is the one synthesis for which we assessed an alternate rapid palladium-based protocol for deprotection of the thiazolidine (Jbara, Maity, Seenaiah, & Brik, 2016). Given the lower purity observed for this product ([Fig. 4-H] compared to [Fig. 4-D]), we recommend the methoxyamine protocol on the solid phase. However, the crude product (73% yield) was still of sufficient purity to use without further purification.

With both large protein segments in hand, we proceeded to the final solution phase ligation. The N-terminal peptide **17** is activated by treatment of sodium nitrite to form the triazole **25**, for conversion to thioester **26** *in situ* for the following ligation. In initial ligation tests, we carried out thioester conversion of the N-terminal followed by addition of the C-terminal segment in ligation buffer (6M Gdn, 0.1M phosphate, 50mM MPAA). However, with ligation carried out at pH 7.3, we found significant hydrolysis of the triazole in competition with thioester formation to reduce our overall yield. We probed this process using the model peptide [H1.2(Lys_89_-Gly_99_)-Dbz-Gly-NH_2_], and found minimal hydrolysis over 6 hours at pH 6. We also assessed whether the process could be simplified by pre-mixing **17** with **24** prior to activation (Fang et al., 2011; Wang et al., 2015), rather than sequential addition of **24** to converted **25**. We found no significant differences in ligation kinetics or yield for this considerably simpler experimental protocol. We therefore converged to ligation via activation of the mixed components, followed by addition of ligation buffer to reach pH ~6.5 for ligation over 4 hours at room temperature.

The ligation product **27** was dialized extensively to remove MPAA, which can quench phosphine-mediated desulfurization (Rohde, Schmalisch, Harpaz, Diezmann, & Seitz, 2011). The final full-length protein was purified by RP-HPLC after desulfurization. Although the ligation reaction proceeds to near completion (Fig 5B, C, D), purification yields are typically poor for histone proteins – which is one driving factor for the development of minimal purification approaches such as solid phase NCL. A 12% isolated yield (assessed by in-gel quantitation against a commercial H1.0 standard) was obtained for the full-length H1.2 after ligation, desulfurization, and RP-HPLC purification. Ligations were characterized by SDS-PAGE (Fig. 5D lane 6) and mass spectrometry (Fig 5F, Supplemental S5.7).

**Figure 5.**
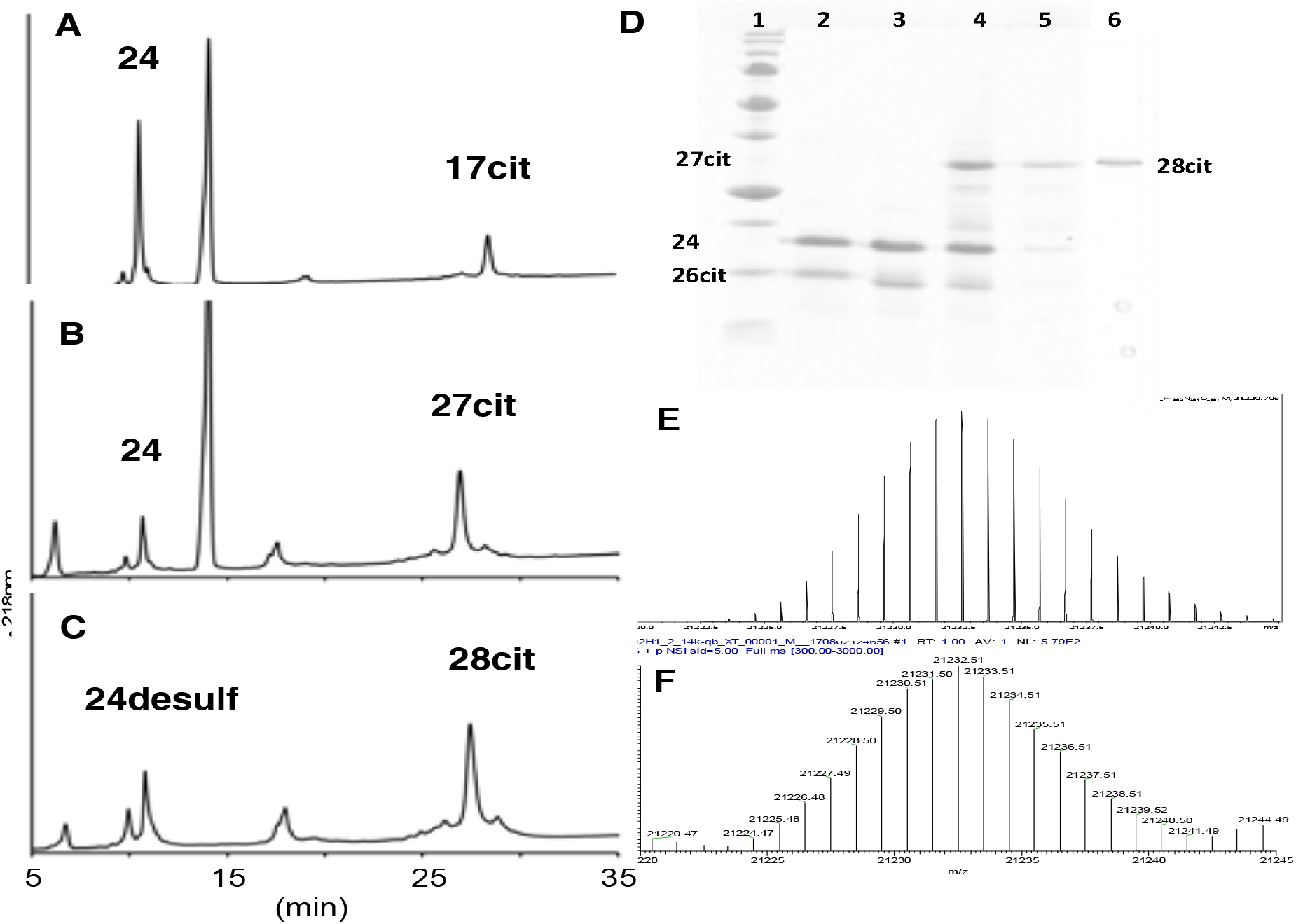
Analysis of solution phase ligation to generate full-length H1.2 **28**. RP-HPLC of A) pre-mixed 17cit and 24 prior to activation and ligation, B) 4 hours timepoint for ligation, C) ligation mixture after desulfurization. Peak identities were confirmed by MALDI-TOF MS (see supplemental information). D) SDS-PAGE analysis of ligation; Lanes 1: MWM; Lane 2, mixture prior to activation; Lane 3, 0 hour prior to pH shift to initiate ligation; Lane 4, 30 minutes timepoint; Lane 5, 2 hours timepoint for ligation; Lane 6, H1.2cit purified by RP-HPLC after desulfurization, taken from a separate gel. E) Calculated isotopic distribution for H1.2 **28**, F) High resolution deconvoluted ESI-MS characterization of purified, synthetic H1.2 **28**, unmodified, taken on a Thermo Q Exactive EMR Orbitrap instrument.

The primary native function of linker histones, including variants H1.0 and H1.2, is to interact with nucleosomes to form a chromatosome, which facilitates chromatin compaction and may alter nucleosome dynamics and local protein interactions (Bernier et al., 2015; Christophorou et al., 2014; Fyodorov et al., 2018; Liao & Mizzen, 2016; Lopez et al., 2015). Recent cryo-EM structural determination provides a consensus structure for the folded core of the H1.0 protein bound to the nucleosome (Bednar et al., 2017), depicted in Fig. 6E, although the majority of the H1 sequence remains relatively unstructured. In order to assess the functionality of our synthetic proteins, we carried out electrophoretic mobility shift assays (EMSA) to compare the binding of commercial unmodified H1.0 (NEB), our synthetic unmodified H1.2, H1.2-R53cit, and H1.2-S172ph to purified mono-nucleosomes (Figure 6). In each case, we observe that the mononucleosome band begins to shift at 5-10 nM with similar changes in electrophoretic mobility indicating that the unmodified and modified synthetic H1.2 form chromatosomes similarly to recombinant H1.0. At higher concentrations (≥30 nM), the nucleosomes do not migrate into the gel, which is consistent with the formation of larger aggregates mediated by H1. EMSA is not expected to be sufficiently sensitive to detect quantitative differences in H1.2 function due to individual modifications. H1 binds nucleosomes at picomolar affinities (Yue, Fang, Wei, Hayes, & Lee, 2016) and has a very slow dissociation rate that is great than 10 minutes (White, Hieb, & Luger, 2016). So, in this EMSA assay, H1 binds stoichiometrically; however, these results clearly demonstrate that the synthetic proteins carry out their expected functions, and that the modifications do not reduce the ability of H1 to form a chromatosome. These syntheses therefore enable future studies to determine how H1 modifications, individually or in combination, may impact nucleosome dynamics or macromolecular interactions, for example transcription factor binding to DNA sites within the linker histone-bound nucleosome (Brehove et al., 2015).

**Figure 6.**
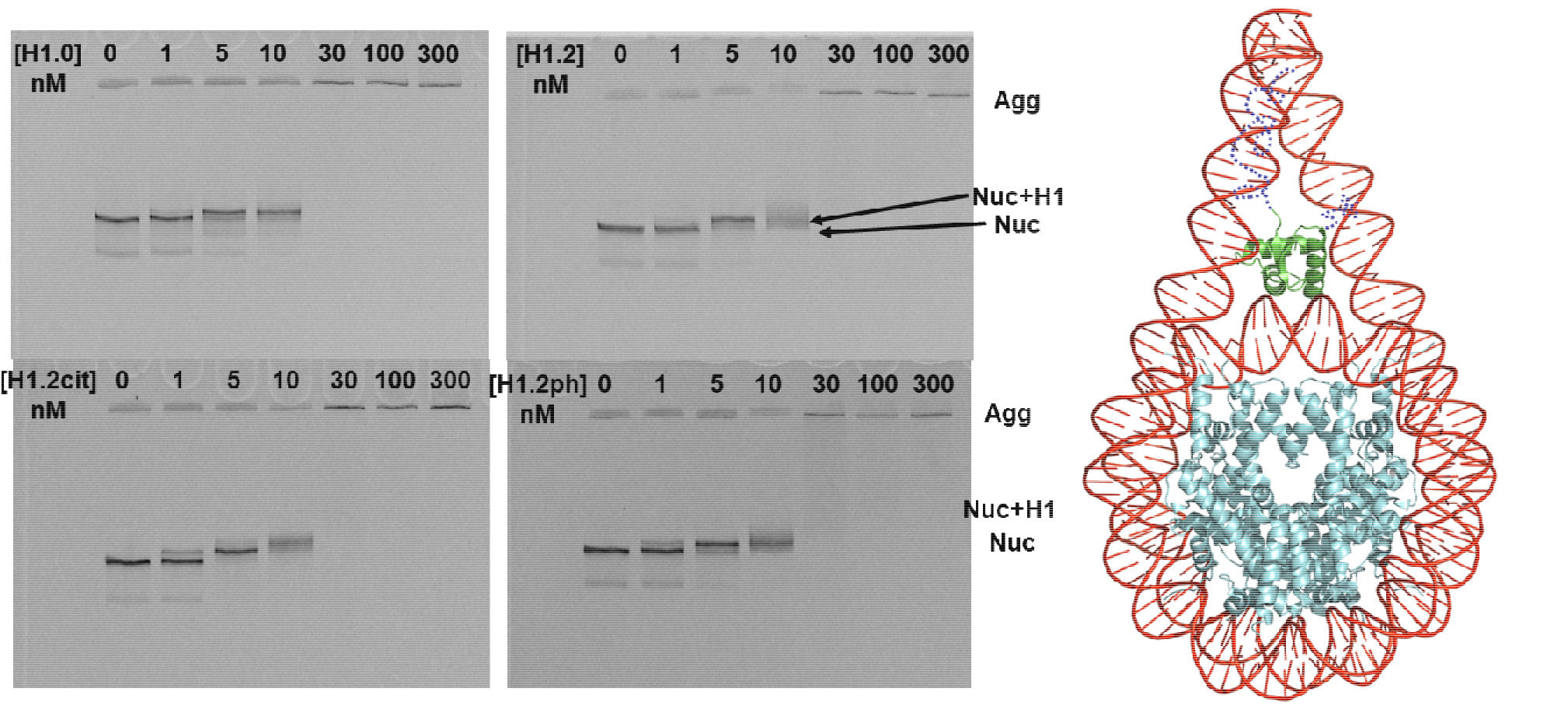
Characterization of H1 binding to nucleosomes. Left: EMSA showing A) H1.0, B) H1.2 **28**, C) H1.2-R53cit (**28cit**), D) H1.2-S172ph **28ph** binding to nucleosomes. Nucleosome band begins to shift at ~5nM H1 and is fully shifted at ~10nM H1. At 30 nM H1, nucleosomes do not migrate into the gel, demonstrating that they are forming H1-mediated aggregates. Right: Structured domain of H1 bound to nucleosome, from 5NL0.pdb^[50]^

In summary, we have developed a practical method to allow the benefits of solid phase protein ligation to be extended to the total synthesis of large proteins by exploiting diaminobenzoic acid (peptide O-aminoanilides) as a cryptic thioester. This allows the traceless assembly of ~100-residue protein segments from small, synthetically accessible peptides. We demonstrated its feasibility by the preparation of unmodified, R53cit, and S172ph modified linker histone H1.2 from 8 component peptides with only 1 purification step, enabling the first characterization of homogenously modified linker histone. We demonstrated its robustness by carrying out solid phase ligation cycles 7 times, with yields ranging from 50-81%, each time with sufficient purity to use in subsequent ligations. While we have demonstrated this for a 212-residue protein assembled from two “blocks” with only one purification step, we hypothesize that the method could logically be extended to one-pot, sequential, or convergent ligation of 3-5 “blocks” for the facile assembly of very large proteins.

## Acknowledgements

This work was supported by the National Institutes of Health (GM121966). We thank Prof. Vicki Wysocki (OSU) and Jing Yan (Wysocki Lab, OSU) for carrying out the high-resolution MS in Fig. 5E,F.

