## Supplemental Data for "Convergent Hybrid Phase Ligation Strategy for Efficient Total Synthesis of Large Proteins Demonstrated for 212-residue Linker Histone H1.2"

### 1 General

In this supplemental section we provide synthesis details, product characterization, but also extensive discussion of technical elements of the synthesis. Section 1 & 2 contains very general notes, reagent sources, etc. Section 3 contains peptide synthesis notes and characterization. For the subfield audience, we draw your attention particularly to section 4, which contains our notes on several issues (and our solutions) surrounding the use of Dbz and MeDbz linkers as cryptic thioesters by Fmoc SPPS. In section 5 we provide our synthesis notes for solid phase NCL of the two components; each is repeated several times, and all data is provided broken out into separate subsections with the detailed methods for each. We then provide our "best practices" summary, derived from extensive experience. In section 6 we detail the final solution phase convergent ligation step and product purification. In section 7, we provided methods for our EMSA characterization of synthetic H1.2 function.

#### 1.1 Materials

Rink Amide MBHA resin LL (100-200 mesh, 0.3-0.4 mmol/g loading) was purchased from Novabiochem. PL-PEGA resin (300-500 µm, 0.2 mmol/g) was purchased from Varian. Amino PEGA resin (150-300 µm, 0.34 mmol/g) was from Novabiochem. N,N-Dimethylformamide $C_3H_7NO$ (DMF), dichloromethane $CH_2Cl_2$ (DCM), acetonitrile $CH_3CN$ (ACN), and diethyl ether $(C_2H_5)_2O$ were purchased from Sigma Aldrich or Fisher Scientific. N-methyl pyrrolidone $C_5H_9NO$ (NMP) was purchased from AGTC Bioproducts or Fisher Scientific. Piperidine $C_5H_{11}N$, 4-nitrophenyl chloroformate $ClCO_2C_6H_4NO_2$ (4-NPCF), 4-mercaptophenyl acetic acid $HSC_6H_4CH_2CO_2H$ (MPAA), $C_9H_{15}O_6P$ Tris(2-carboxylmethyl)phosphine (TCEP), N,N-diisopropylethylamine $C_8H_{19}N$ (DIEA), Phenylsilane $C_6H_8Si$, and Tetrakis(triphenylphosphine)palladium(0) ($Pd(PPh_3)_4$) were purchased from Sigma Aldrich. Fmoc or Boc protected amino acids were purchased from AAPPTec and Novabiochem. Fmoc-6-Aminohexanoic acid (Fmoc-Ahx-OH), Fmoc-L-norleucine (Fmoc-Nle-OH), and 4-dimethylaminopyridine $C_7H_{10}N_2$ (DMAP) were purchased from Novabiochem. (2-(7-aza-1H-benzotriazole-1-yl)-1,1,3,3-tetramethyluronium. Oxyma-Pure was purchased from Novabiochem. (2-(7-aza-1*H*-benzotriazole-1-yl)-1,1,3,3-tetramethyluronium hexafluorophosphate $C_{10}H_{15}F_6N_6OP$ (HATU), 2-(6-chloro-1H-benzotriazole-1-yl)-1,1,3,3-tetramethylaminium hexafluorophosphate $C_{11}H_{15}ClF_6N_5OP$ (HCTU), and 1-hydroxy-6-chlorobenzotriazole $C_6H_4ClN_3O$ (6-Cl-HOBt) were purchased from AAPPTec. Acetic anhydride and sodium 2-sulfanylethane sulfonate $C_2H_5NaO_3S_2$ (MESNA) was purchased from Sigma-Aldrich. N,N'-diisopropylcarbodiimide $C_7H_{14}N_2$ (DIC) was purchased from Chem-Implex International. VA-044-US $C_{12}H_{22}N_6{\cdot}2HCl$ was purchased from Wako Chemicals. Ultra-pure guanidine-HCl $CH_6ClN_3$ (GuHCl) was purchased from MP Biomedicals. 3,4-diaminobenzoic acid (Dbz) was purchased from Sigma-Aldrich. Allyl chloroformate $C_4H_5ClO_2$ was purchased from Acros Organics. Triisopropylsilane $C_9H_{22}Si$ (TIS) was purchased from Sigma-Aldrich. α-Cyano-4-hydroxycinnamic acid $C_{10}H_7NO_3$ (HCCA) was purchased from Bruker Daltonics.

#### 1.2 RP-HPLC

Analytical reverse phase HPLC (RP-HPLC) was run on a Shimadzu or Waters instrument using an analytical column (Supelco C18 15 cm × 4.6 mm × 5 µm, flow rate 0.9 mL/min). Preparative RP-HPLC was run on a Waters instrument using a semi-preparative column (Supelco C18 25 cm × 10 mm × 10 µm, flow rate 5 mL/min), or a preparative column (Supelco C18 25 cm × 21.2 mm × 10 µm, flow rate 18 mL/min). Mini-preparative RP-HPLC for full-length linker histone H1.2 was run on a Shimadzu instrument using an analytical column (Discovery BIO Wide Pore C5 10 cm × 4.6 mm × 5 µm). Buffer A was 0.1 % TFA in water, and Buffer B was 1:9 water:acetonitrile, 0.1% TFA. Gradients in the ESI are reported as the percentage Buffer B in Buffer A if not specified.

##### 1.3 Mass spectrometry

Peptide masses were confirmed by MALDI-TOF-MS (Bruker Daltonics Microflex) and analyzed using and flexAnalysis 3.3 software. α-Cyano-4-hydroxycinnamic acid (HCCA) was used for the matrix and Peptide Calibration Standard II (Bruker) or Protein Calibration Standard I (Bruker) was used for calibration. To calculate masses, spectra were processed with Savitzky–Golay smoothing and peaks were picked using the Centroid algorithm in Bruker FlexAnalysis software. However, unprocessed/raw spectra are displayed throughout the supplemental. All expected masses are reported as the mass average.

### 2 Peptide Preparation

##### 2.1 Synthesis of 3-Fmoc-Dbz-OH

3-Fmoc-Dbz-OH was prepared as described previously[1]. In brief, 3,4-Diaminobenzoic acid (1 g, 6.5 mmol) was resuspended in 125 mL 1:1 $CH_3CN$:$NaHCO_3$. Reaction was initiated by the dropwise addition of Fmoc-OSu (2.4 g, 7.1 mmol) in 15 mL 1:1 $CH_3CN$:$NaHCO_3$ and proceeded for 2 h. HCl was added to a final pH of 1.0, and the mixture was filtered. Filtrate was dissolved in 4 mL DMSO, precipitated with acidified reaction buffer, washed extensively, and dried under vacuum to yield a light gray product. Product identity and purity was validated by NMR spectroscopy and following peptide synthesis RP-HPLC-MS analysis.

##### 2.2 Peptide Synthesis – general methods

Peptides were synthesized using the standard Fmoc-N-α protection strategies either manually (for short sequences) or on an automated AAPPTec APEX 396 or CEM Liberty Lite synthesizer. This is specified for each peptide in the relevant subsection in Section 3.

Peptides synthesized on an AAPPTec APEX 396 automated synthesizer proceeded as following for each coupling cycle unless otherwise specified: 5 min Fmoc deprotection with 20% piperidine for two times, 30min coupling with HCTU/DIEA, 5min capping with capping solution (300 mM 6-Cl-HOBt and 300 mM Acetic anhydride in 1:9 DCM: DMF).

The peptides synthesized on a CEM Liberty Lite synthesizer were carried out with microwave assistance. For standard syntheses when MeDbz was used as a linker, syntheses followed the following procedure for every coupling cycle: 90 seconds Fmoc deprotection with 10% piperazine or 20% piperidine at 88 $^{o}$C, 180 seconds amino acid/DIC/Oxyma (1:1:1, 5 equivalents) coupling at 88 $^{o}$C, 120 seconds capping with 10 equivalents of DIEA/acetic anhydride. When Dbz(Alloc) was used as a linker, the deprotection conditions were modified to be carried out at 25 °C; 2x 5 min. Other conditions were not changed.

Fmoc-MeDbz or Fmoc-Dbz was manually coupled to rink amide MBHA resin with HCTU activation (amino acid/HCTU/DIEA=2.2/2/4.4 equivalents to resin loading). Alloc protection (overnight treatment of the resin with 250 mM allyl chloroformate in DCM with 1 eq DIEA to resin loading) was followed to prepare Fmoc-Dbz(Alloc) resin.

Convergent peptide base resin Fmoc-Dbz(Alloc)-RGK(C) or Dbz(Alloc)-GK(C) were synthesized manually. Orthogonal Fmoc-Lys (Alloc)-OH was used for the first amino acid. Boc-Cys(trt)-OH was coupled to the side chain amine after removal of Alloc group.

Amino acid coupling directly onto 4-Alloc-Dbz resin was accomplished using HATU activation (amino acid/HATU/DIEA=16.5/15/33) with 1 hour coupling time[1], followed by acetyl capping using a fresh solution containing 15% acetic anhydride and 15% DIEA in DMF. Amino acid following the HMBA linker was double-coupled as the symmetric anhydride: 10 equivalents Fmoc-AA-OH was dissolved in DCM. 5 equivalents of DIC was added and incubated on ice for 30 minutes. The filtrate and 0.1 equivalent DMAP were added to DMF-swollen resin. The reaction was allowed to proceed for 1 hour.

Dbz (Alloc) deprotection was accomplished on resin by treatment with 0.35 eq Pd $(PPh_3)_4$ and 20 eq Phenyl silane in DCM for 45 minutes. If not specified, Dbz was converted into Nbz by treating the resin with 50mM NPCF in DCM for 30 minutes, followed by 0.5 M DIEA in DMF for 15 minutes as reported[2]. MeDbz is converted into MeNbz by treating the resin with 0.25M 4-NPCF in DCM for 2h, followed by 1M DIEA treatment in DMF for 1h as described previously[3]. Other conditions will be specified when needed.

All peptides were cleaved in 95:2.5:2.5 TFA:TIS:water for 2 hours where otherwise specified. TFA was reduced with a stream of nitrogen, and peptides were precipitated and washed with cold $Et_2O$, then resuspended in water and lyophilized before analysis and purification.

##### 2.3 Base Resin for SP-NCL

Two different PEGA resins were used in this paper: 0.2 mmol/g PL-PEGA resin (300-500 µm, Agilent Tech-Varian, no longer commercially available) or 0.35 mmol/g Amino PEGA resin (150-300 µm, Novabiochem).

In order to reduce steric crowding, the loading of the resin was reduced to 0.05 mmol/g by coupling of the resin with a mixture of 1:3 (PL-PEGA) or 1:6 (Amino-PEGA) Fmoc-Gly-OH:Boc-Gly-OH. Standard coupling using Fmoc-SPPS was performed after the loading cut. The sequences of the base resins used are listed below along with their theoretical loading after synthesis through the Fmoc-Thz. Gly where the loading cut is performed is indicated in bold. The extended peptide sequence was used to add sufficient length to the linker segment to allow for RP-HPLC and MALDI-TOF MS detection of incomplete ligation products. The base resins were stored in methanol, and the substitution was assessed based on the volume for methanol-swollen beads.

PL-PEGA resin: Fmoc-Thz-Ala-Ahx-Tyr-Lys-Gly-Rink-**Gly**–PEGA, 0.0025 mmol/mL
Amino PEGA resin: Fmoc-Thz-Ala-Ahx-Tyr-Lys-Gly-Rink-**Gly**–PEGA, 0.0033 mmol/mL

### 3 Preparation and Characterization of Linker Histone H1.2 peptides

This section provides brief details on the synthesis and characterization of all peptides required for the preparation of linker histone H1.2, modified or unmodified.

##### 3.1 Synthesis of H1.2-($Ser_1$-$Ala_{23}$)-Dbz-R-OH or H1.2-($Ser_1$-$Ala_{23}$)-MeNbz-$NH_2$

SETAPAAPAAAPPAEKAPVKKKA-X (X=Dbz-R or MeNbz)
X=Dbz-R-OH:
mFmoc-Dbz was manually coupled to the preloaded Fmoc-Arg-Wang resin. The remaining residues of peptide were coupled on an AAPPTec APEX 396 automated synthesizer. N-terminal Fmoc was removed with 20% piperidine manually. The peptide was purified using RP-HPLC with a gradient of 10-40% HPLC buffer B. The peptide will be activated by sodium nitrite treatment before native chemical ligation.

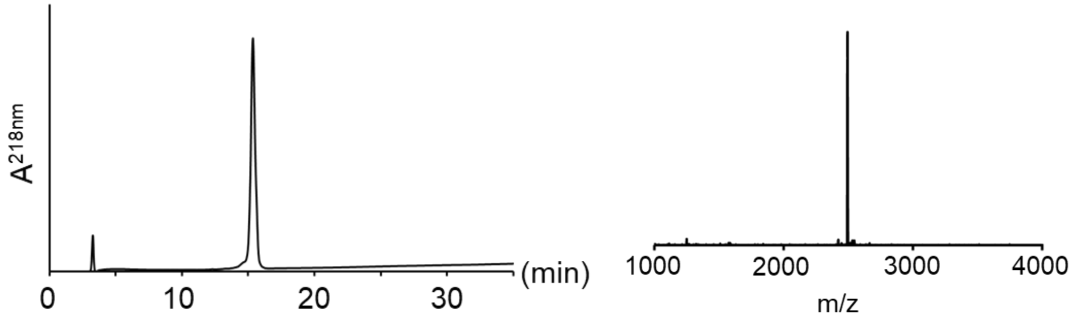

**Figure S1**. RP-HPLC (left 10-40 % B) of purified H1.2-($Ser_1$-$Ala_{23}$)-Dbz-R. MALDI-TOF (right) of purified H1.2-($Ser_1$-$Ala_{23}$)-Dbz-R-OH. H1.2-(Ser1-Ala23)-Dbz-R-OH: $[M + H]^+$ observed: *m/z* 2492, expected: *m/z* 2493.

X=MeNbz:
The peptide was coupled on CEM Liberty Lite synthesizer using preloaded Fmoc-MeDbz resin. The N-terminal residue was added as Boc-Thz-OH. Peptidyl MeDbz was converted to MeNbz by 4-nitrophenyl chloroformate and DIEA treatment. Final peptide was cleaved off the resin and deprotected with TFA. The peptide was purified using RP-HPLC in a gradient of 5-25% buffer B over 40min.

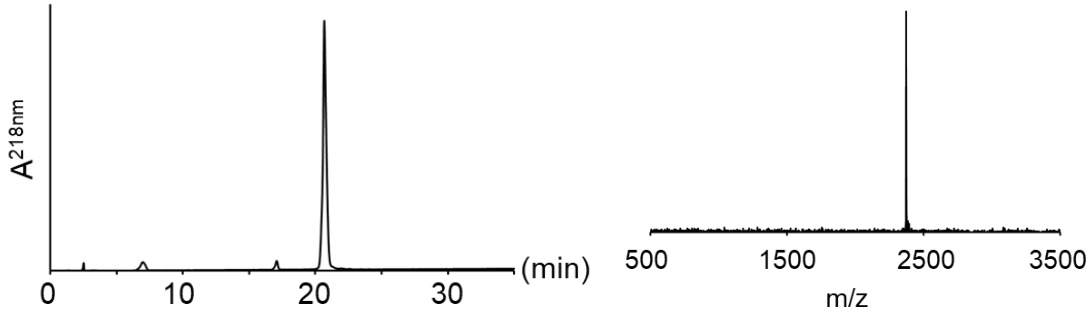

**Figure S2**. RP-HPLC (left 5-25 % B) and MALDI-TOF MS (right) of purified H1.2-($Ser_1$-$Ala_{23}$)-MeNbz. H1.2-($Ser_1$-$Ala_{23}$)-MeNbz: $[M + H]^+$ observed: *m/z* 2373, expected: *m/z* 2376.

##### 3.2 Synthesis of H1.2-(Thz$_{24}$-Ala$_{48}$)-Nbz or MeNbz

Thz-KKAGGTPRKASGPPVSELITKAVA-X (X=Nbz or MeNbz)

X=Nbz:

The first Ala was manually coupled to prepared mFmoc-Dbz(Alloc) rink amide MBHA resin. The remaining residues were coupled on an AAPPTEC APEX 396 automated synthesizer. The N-terminal residue was added as Boc-Thz-OH. Alloc was removed with catalyst $Pd(PPh_3)_4$ and phenylsilane as scavenger. Dbz was converted into Nbz on-resin by 4-NPCF and DIEA treatment. Peptide was cleaved from resin with a standard cleavage cocktail (TFA/H2O/TIPS 95:2.5:2.5). Peptide was purified by RP-HPLC using a gradient of 20-40% HPLC Buffer B over 40 minutes.

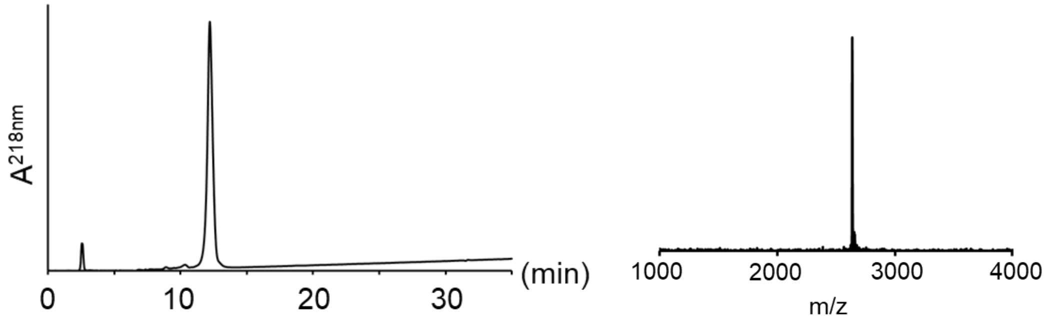

**Figure S3**. RP-HPLC (left 20-40 % HPLC buffer B) and MALDI-TOF MS (right) of purified H1.2-(Thz$_{24}$-Ala$_{48}$)-Nbz: $[M + H]^+$ observed: *m/z* 2637, expected: *m/z* 2639.

X=MeNbz:

The peptide was synthesized on CEM Liberty Lite synthesizer using preloaded Fmoc-MeDbz resin. The N-terminal residue was added as Boc-Thz-OH. Peptidyl MeDbz was converted ed to MeNbz by 4-nitrophenyl chloroformate and DIEA treatment. Final peptide was cleaved and deprotected with TFA. Peptide was purified by RP-HPLC using a gradient of 20-45% HPLC Buffer B over 40 minutes.

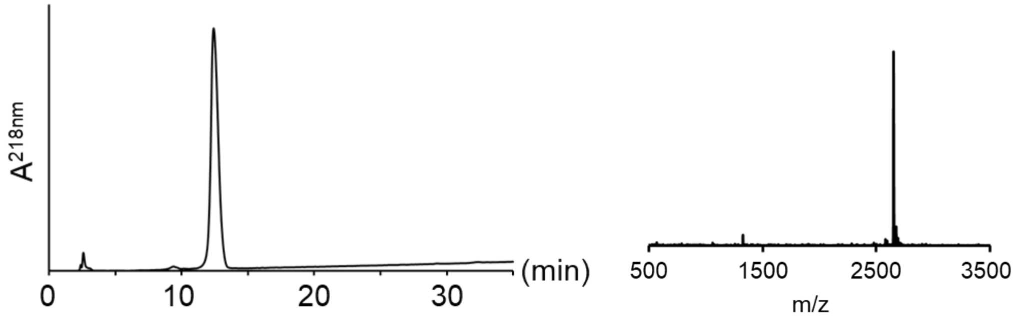

**Figure S4**. RP-HPLC (left 20-45 % HPLC buffer B) and MALDI-TOF MS (right) of purified H1.2-(Thz$_{24}$-Ala$_{48}$)-MeNbz: $[M + H]^+$ observed: *m/z* 2652, expected: *m/z* 2653.

##### 3.3 Synthesis of H1.2-($Thz_{49}$-$Ala_{66}$)-Nbz or MeNbz

Thz-SKERSGVSLAALKKALA-X (X=Nbz or MeNbz)

X=Nbz:

The first Ala was manually coupled to prepared mFmoc-Dbz(Alloc) rink amide MBHA resin. The remaining residues were coupled on an AAPPTEC APEX 396 automated synthesizer. The N-terminal residue was added as Boc-Thz-OH. Alloc was removed with catalyst $Pd(PPh_3)_4$ and phenylsilane as scavenger. Dbz was converted into Nbz on-resin by 4-NPCF and DIEA treatment. Peptide was cleaved from resin with a standard cleavage cocktail (TFA/H2O/TIPS 95:2.5:2.5). Peptide was purified by RP-HPLC using a gradient of 15-45% HPLC Buffer B over 40 minutes.

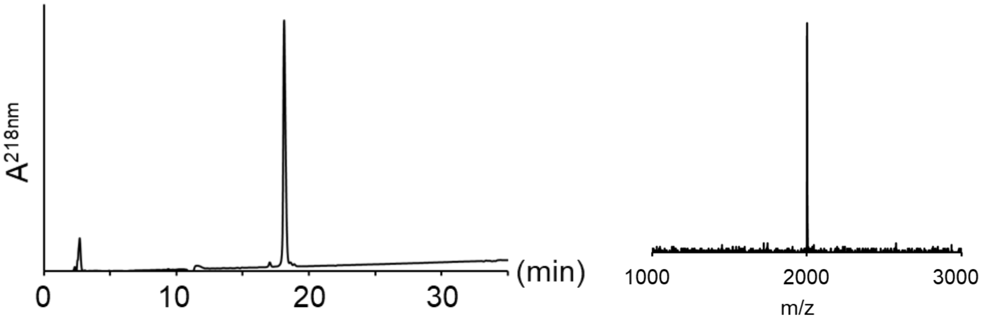

**Figure S5**. RP-HPLC (middle 15-45 % B) and MALDI-TOF MS (right) of purified H1.2-($Thz_{49}$-$Ala_{66}$)-Nbz: $[M + H]^+$ observed: *m/z* 2003, expected: *m/z* 2004.

X=MeNbz:

The peptide was synthesized on CEM Liberty Lite automated synthesizer using preloaded Fmoc-MeDbz resin. The N-terminal residue was added as Boc-Thz-OH. MeDbz was converted into MeNbz on-resin by 4-NPCF and DIEA treatment at 50 $^{o}$C and more details about the conversion will be discussed later. Peptide was cleaved with TFA and purified by RP-HPLC using a gradient of 15-45% Buffer B over 40 minutes.

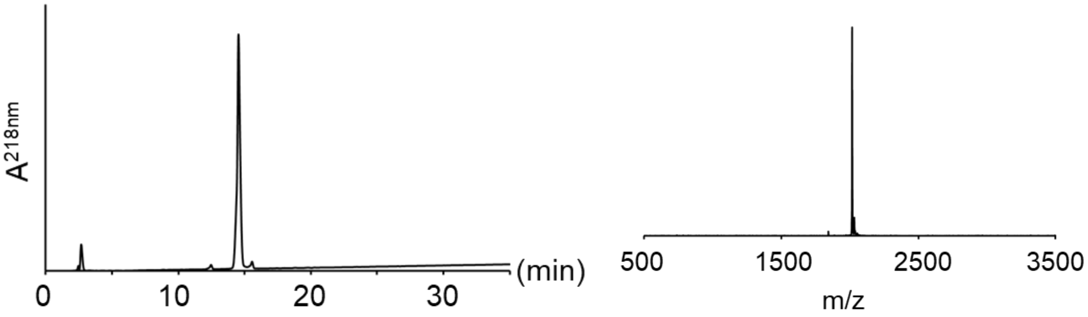

**Figure S6**. RP-HPLC (left 15-45 % Buffer B) and MALDI-TOF MS (right) of purified H1.2-($Thz_{49}$-$Ala_{66}$)-MeNbz. H1.2-($Thz_{49}$-$Ala_{66}$)-MeNbz: $[M + H]^+$ observed: *m/z* 2017, expected: *m/z* 2018.

##### 3.4 Synthesis of H1.2-($Thz_{67}$-$Gly_{99}$)-Dbz-GK(C) or Dbz-RGK(C)

Thz-AGYDVEKNNSRIKLGLKSLVSKGTLVQTKGTG-DBz-X (X=RGK(C) or GK(C))

The peptides were synthesized on CEM Liberty Lite automated synthesizer using preloaded Fmoc-Gly-Dbz(Alloc)-GK(C) or Fmoc-Gly-Dbz(Alloc)-RGK(C) resin. The N-terminal residue was added as Boc-Thz-OH. Room temperature Fmoc deprotection was applied to avoid side reaction on Dbz(Alloc), which will be discussed in detail later. Alloc was removed after synthesis using $Pd(PPh_3)_4$ and peptide was cleaved with TFA. The peptides were purified by RP-HPLC either using a gradient of 20-35% or 15-35% Buffer B over 40 minutes.

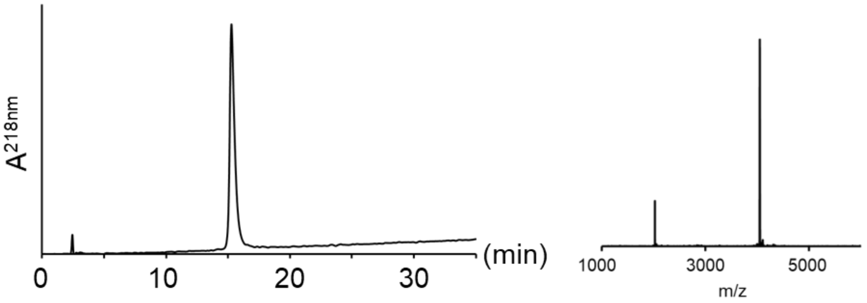

**Figure S7**. RP-HPLC (left 20-35% buffer B) and MALDI-TOF MS (right) of purified H1.2-($Thz_{67}$-$Gly_{99}$)-Dbz-RGK(C): $[M + H]^+$ observed: *m/z* 4055, expected: *m/z* 4057.

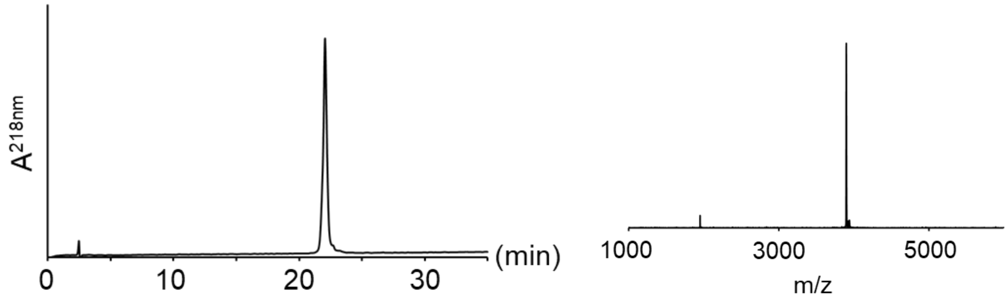

**Figure S8**. RP-HPLC (left 15-35% buffer B) and MALDI-TOF MS (right) of purified H1.2-($Thz_{67}$-$Gly_{99}$)-Dbz-GK(C): $[M + H]^+$ observed: *m/z* 3901, expected: *m/z* 3901.

##### 3.5 Synthesis of H1.2-($Thz_{100}$-$Ala_{133}$)-MeNbz

Thz-SGSFKLNKKAASGEAKPKVKKAGGTKPKKPVGA-MeNbz

The peptide was synthesized on CEM Liberty Lite automated synthesizer using preloaded Fmoc-MeDbz resin. The N-terminal residue was added as Boc-Thz-OH. MeDbz was converted into MeNbz on-resin by 4-NPCF and DIEA treatment. Peptide was cleaved from resin with and purified by RP-HPLC using a gradient of 10-30% HPLC Buffer B over 40 minutes.

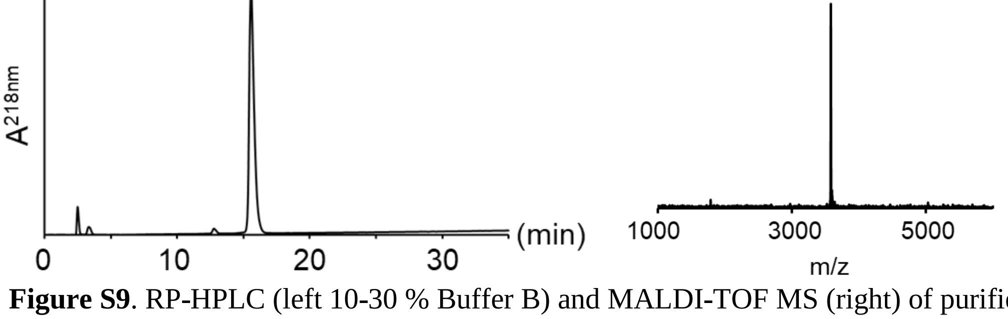

**Figure S9**. RP-HPLC (left 10-30 % Buffer B) and MALDI-TOF MS (right) of purified H1.2-($Thz_{100}$-$Ala_{133}$)-MeNbz: $[M + H]^+$ observed: *m/z* 3585, expected: *m/z* 3585.

**3.6 Synthesis of H1.2-(Thz$_{134}$-Ala$_{162}$)-Nbz or MeNbz**

Thz-KKPKKAAGGATPKKSAKKTPKKAKKPAA-X (X=Nbz or MeNbz)

X=Nbz:

The first Ala was manually coupled to prepared mFmoc-Dbz(Alloc) rink amide MBHA resin. The remaining residues were coupled on an AAPPTEC APEX 396 automated synthesizer. The N-terminal residue was added as Boc-Thz-OH. Alloc was removed with catalyst $Pd(PPh_3)_4$ and phenylsilane as scavenger. Dbz was converted into Nbz on-resin by 4-NPCF and DIEA treatment. Peptide was cleaved from resin with a standard cleavage cocktail (TFA/H2O/TIPS 95:2.5:2.5). Peptide was purified by RP-HPLC using a gradient of 0-25% HPLC Buffer B over 40 minutes. This peptide hydrolyzes fast, which is indicated by the small peak in RP-HPLC trace of purified sample.

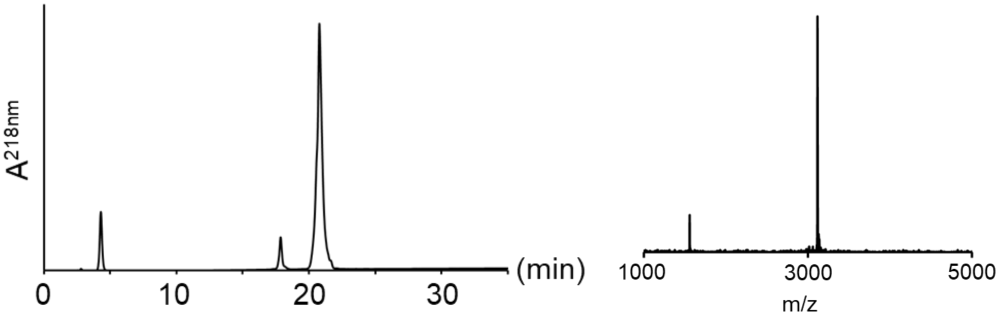

**Figure S10**. RP-HPLC (left 0-25 % B) and MALDI-TOF MS (right) of purified H1.2-(Thz$_{134}$-Ala$_{162}$)-Nbz: $[M + H]^+$ observed: *m/z* 3120, expected: *m/z* 3121.

X=MeNbz:

The peptide was synthesized on CEM Liberty Lite automated synthesizer using preloaded Fmoc-MeDbz resin. The N-terminal residue was added as Boc-Thz-OH. MeDbz was converted into MeNbz on-resin by 4-NPCF and DIEA treatment. Peptide was cleaved from resin with TFA and purified by RP-HPLC using a gradient of 5-25% HPLC Buffer B over 40 minutes.

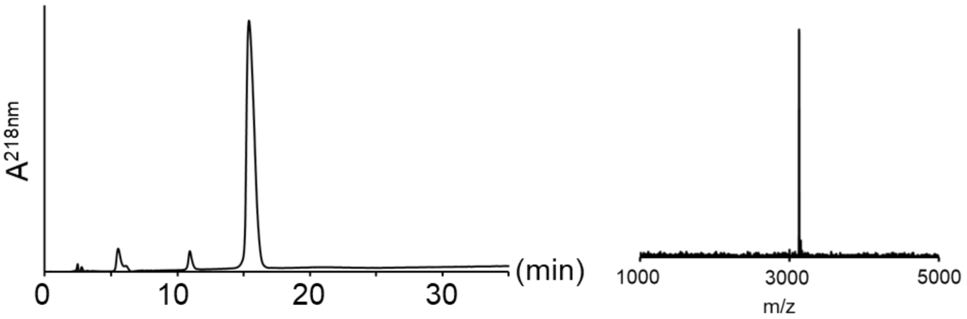

**Figure S11**. RP-HPLC (left 5-25 % B) and MALDI-TOF MS (right) of purified H1.2-(Thz$_{134}$-Ala$_{162}$)-MeNbz: $[M + H]^+$ observed: *m/z* 3133, expected: *m/z* 3135.

##### 3.7 Synthesis of H1.2-($Thz_{163}$-$Ala_{188}$)-Nbz or MeNbz

Thz-TVTKKVAKSPKKAKVAKPKKAAKSA-X (X=Nbz or MeNbz)

X=Nbz:

The first Ala was manually coupled to prepared mFmoc-Dbz(Alloc) rink amide MBHA resin. The remaining residues were coupled on an AAPPTEC APEX 396 automated synthesizer. The N-terminal residue was added as Boc-Thz-OH. Alloc was removed with catalyst $Pd(PPh_3)_4$ and phenylsilane as scavenger. Dbz was converted into Nbz on-resin by 4-NPCF and DIEA treatment. Peptide was cleaved from resin with a standard cleavage cocktail (TFA/H2O/TIPS 95:2.5:2.5). Peptide was purified by RP-HPLC using a gradient of 5-25% HPLC Buffer B over 40 minutes.

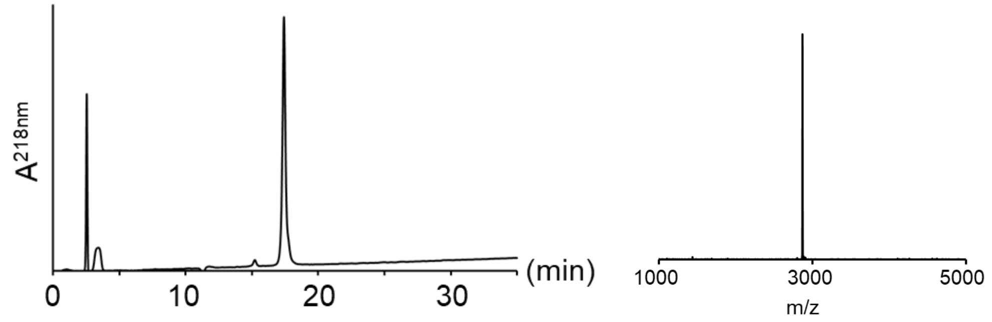

**Figure S12**. RP-HPLC (left 5-25 %Buffer B) and MALDI-TOF MS (right) of purified H1.2-($Thz_{163}$-$Ala_{188}$)-Nbz: $[M + H]^+$ observed: *m/z* 2868, expected: *m/z* 2869.

X=MeNbz:

The peptide was synthesized on CEM Liberty Lite automated synthesizer using preloaded Fmoc-MeDbz resin. The N-terminal residue was added as Boc-Thz-OH. MeDbz was converted into MeNbz on-resin by 4-NPCF and DIEA treatment. Peptide was purified by RP-HPLC using a gradient of 15-25% HPLC Buffer B over 40 minutes.

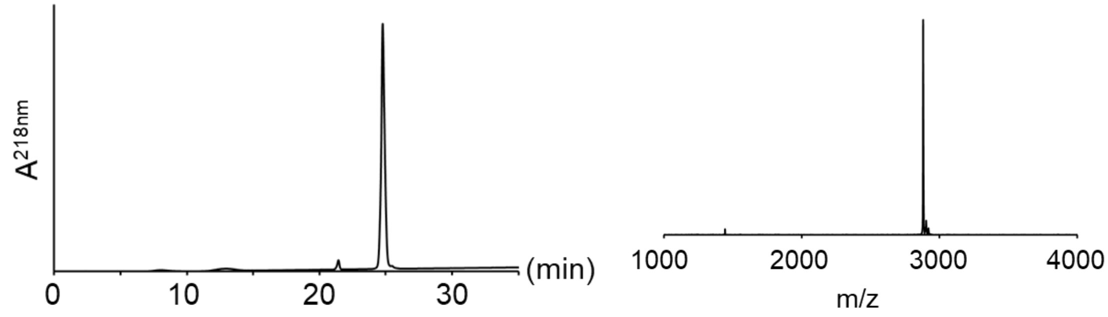

**Figure S13**. RP-HPLC (left 5-25 % Buffer B) and MALDI-TOF MS (right) of purified H1.2-($Thz_{163}$-$Ala_{188}$)-MeNbz: $[M + H]^+$ observed: *m/z* 2882, expected: *m/z* 2884.

**3.8 Synthesis of H1.2-(Thz$_{189}$-Lys$_{212}$)-HMBA-RG-Nbz or MeNbz**

Thz-KAVKPKAAKPKVVKPKKAAPKKK-HMBA-RG-X (X=Nbz or MeNbz)

X=Nbz:

The peptide was synthesized in on an AAPPTEC APEX 396 automated synthesizer using preloaded Fmoc-Lys-HMBA-RG-Dbz(Alloc) resin. The N-terminal residue was added as Boc-Thz-OH. Alloc was removed with catalyst $Pd(PPh_3)_4$ and phenylsilane as scavenger. Dbz was converted into Nbz on-resin by 4-NPCF and DIEA treatment. Peptide was cleaved from resin with a standard cleavage cocktail (TFA/H2O/TIPS 95:2.5:2.5). Peptide was purified by RP-HPLC using a gradient of 0-30% HPLC Buffer B over 40 minutes.

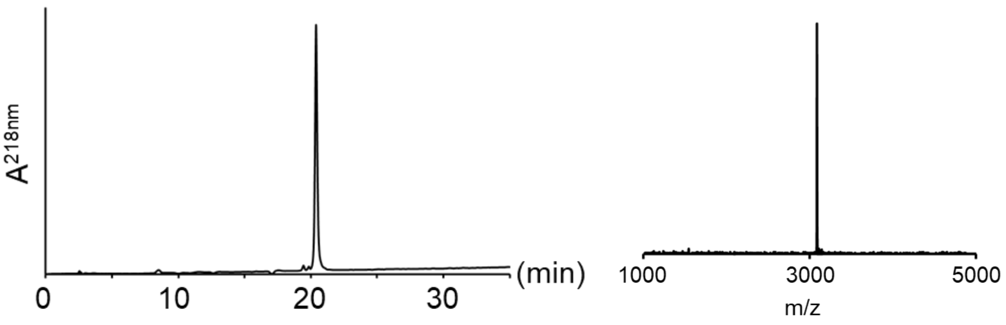

**Figure S14**. RP-HPLC (left 0-30 % B) and MALDI-TOF MS (right) of purified H1.2-(Thz$_{189}$-Lys$_{212}$)-HMBA-RG-Nbz: $[M + H]^+$ observed: *m/z* 3088, expected: *m/z* 3092.

X=MeNbz:

The peptide was synthesized on CEM Liberty Lite automated synthesizer using preloaded Fmoc-Lys-HMBA-RG-MeDbz resin. The N-terminal residue was added as Boc-Thz-OH. Room temperature Fmoc deprotection was applied due to the presence of ester bond. MeDbz was converted into MeNbz on-resin by 4-NPCF and DIEA treatment. Peptide was cleaved from resin with TFA and purified by RP-HPLC using a gradient of 5-25% HPLC Buffer B over 40 minutes.

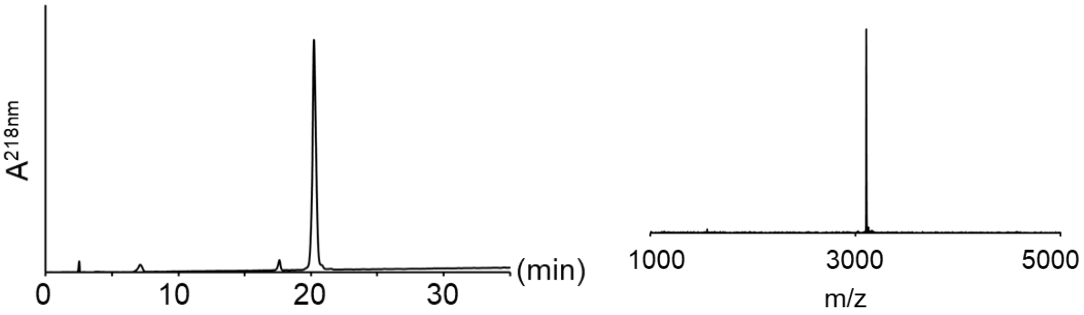

**Figure S15**. RP-HPLC (left 5-25 % buffer B) and MALDI-TOF MS (right) of purified H1.2-(Thz$_{189}$-Lys$_{212}$)-HMBA-RG-MeNbz: $[M + H]^+$ observed: *m/z* 3105, expected: *m/z* 3106.

**3.9 Synthesis of H1.2-($Thz_{49}$-$Ala_{66}$)-R53cit-MeNbz**

Thz-SKE-R53cit-SGVSLAALKKALA-MeNbz

The peptide was synthesized on CEM Liberty Lite automated synthesizer using preloaded Fmoc-MeDbz resin. The N-terminal residue was added as Boc-Thz-OH. Fmoc-Cit-OH was used at position 53 instead of Fmoc-Arg(Pbf)-OH to achieve the modification. MeDbz was converted into MeNbz on-resin by 4-NPCF and DIEA treatment at 50 $^{o}$C. Peptide was purified by RP-HPLC using a gradient of 20-40% HPLC Buffer B over 40 minutes.

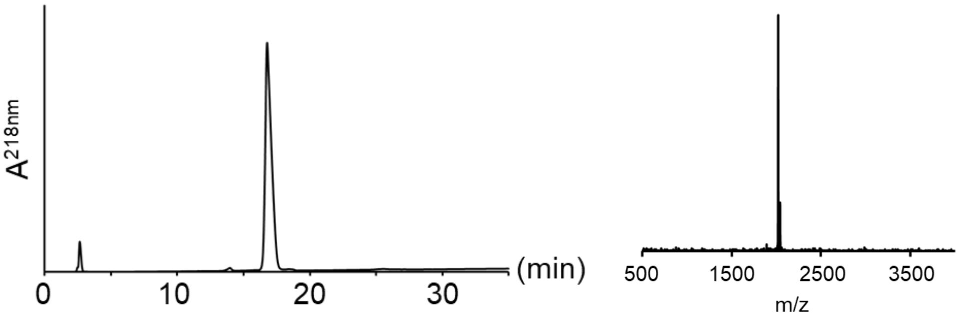

**Figure S16**. RP-HPLC (left 20-40 % B) and MALDI-TOF MS (right) of purified H1.2-($Thz_{49}$-$Ala_{66}$)-R53cit-MeNbz: $[M + H]^+$ observed: *m/z* 2019, expected: *m/z* 2019.

**3.10 Synthesis of H1.2-($Thz_{163}$-$Ala_{188}$)-S172ph-MeNbz**

Thz-TVTKKVAK-S172ph-PKKAKVAKPKKAAKSA-MeNbz

The peptide was synthesized on CEM Liberty Lite automated synthesizer using preloaded Fmoc-MeDbz resin. The N-terminal residue was added as Boc-Thz-OH. Fmoc-Ser(PO(OBzl)OH)-OH was used at position 172 instead of Fmoc-Ser(OtBu)-OH to achieve the phosphorylation. Room temperature Fmoc deprotection was applied before the coupling of next Lysine. MeDbz was converted into MeNbz on-resin by 4-NPCF and DIEA treatment at 50 $^{o}$C. Peptide was cleaved from resin with reagent K (TFA/thioanisole/phenol/$H_2O$/EDT 82.5/5/5/5/2.5). Peptide was purified by RP-HPLC using a gradient of 5-25% HPLC Buffer B over 40 minutes.

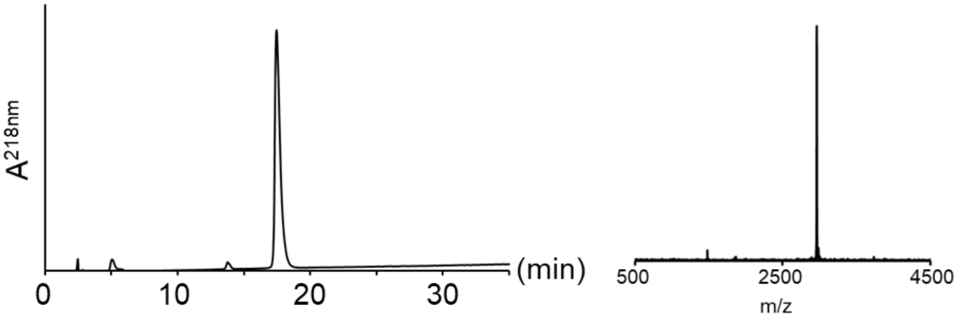

**Figure S17**. RP-HPLC (left 5-25 % B) and MALDI-TOF MS (right) of purified H1.2-($Thz_{163}$-$Ala_{188}$)-S172ph-MeNbz: $[M + H]^+$ observed: *m/z* 2962, expected: *m/z* 2964.

#### 4 Synthesis notes for using and selecting between Dbz(Alloc) and MeDbz

The diaminobenzoic acid moiety is versatile as a cryptic thioester, but also requires careful optimization for different use conditions and outcomes. For example, under field-standard room temperature and microwave-assisted peptide synthesis, the linker is susceptible to acylation at the second amine, particularly with small C-terminal residues such as Gly or Ala, or under standard capping conditions[1]– although careful optimization of coupling conditions has been proposed to alleviate some of these issues[4]. In addition to the simple Dbz moiety, we use two protected variants as required for our syntheses, protected Dbz(Alloc)[1] and MeDbz[3].

Here, we provide the flowchart (Scheme S1) that our researchers currently use to select the derivative most likely to give a successful outcome with minimal effort/customization of the synthesis. In sections 4.1-4.4, we describe some of the issues that we have encountered with each derivative that inform these choices.

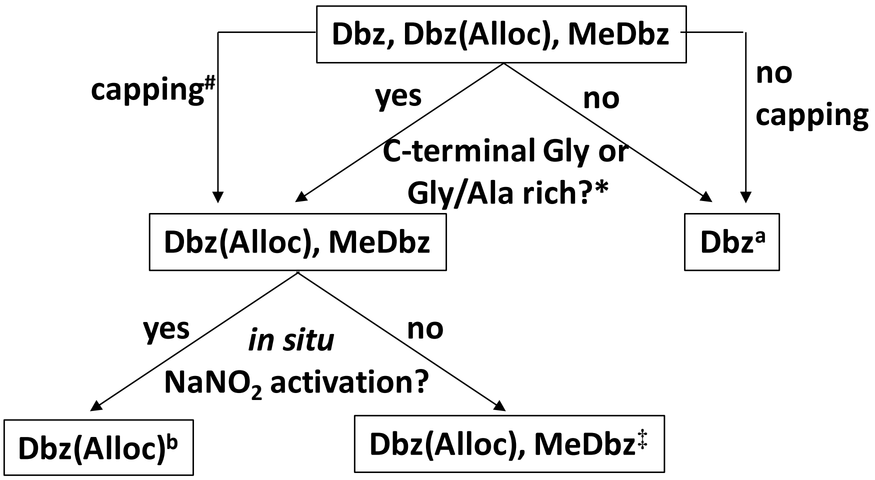

**Scheme S1**. Selection guide of Dbz linker variants for practical peptide synthesis. Quick notes follow:
* Dbz gives satisfactory results under multiple experimental conditions if the residue directly attached has significant steric bulk or is branched at the β-carbon, such as Val or Ile. In these cases, Dbz is the most affordable and simplest choice. However, with highly activating conditions or microwave assistance, we see acylation if that residue is Gly or Ala. In these cases, we typically select an alternate variant.

### Mild capping conditions are compatible with MeDbz; partial acylation is observed with stringent capping conditions. See also [7].
‡ MeDbz is commercially available, and easily compatible with microwave-assisted synthesis. However, for several peptides we have found that significant optimization is required for clean conversion to MeNbz (see for example S4.4), and analysis is complicated by acid-induced rearrangement of MeDbz (S4.3). We have not yet identified the sequence factors underlying this. Currently, we invest the up-front effort to optimize this process for peptides we intend to synthesize multiple times. However, for peptides we plan to synthesize only once, we typically use Dbz(Alloc) as it gives predictable results across a broad range of sequences.

Comments for microwave synthesis:
**a:** Under very specific sequence conditions described above, Dbz can be used with microwave synthesis, with amino acid/Oxyma/DIC ratio of 5:5:4 for heated coupling.
**b:** Dbz(Alloc) is not stable under standard Fmoc-deprotection conditions in heated, microwave-assisted SPPS. However, it is stable under heated DIC/Oxyma coupling conditions, with room-temperature Fmoc-deprotection cycles, as described in S4.1-4.2.

##### 4.1 Observation of Dbz(Alloc) side reactions during microwave-assisted synthesis

Dbz(Alloc) has been extensively assessed for use in room temperature peptide synthesis. During this project, we purchased a CEM Liberty Lite synthesizer, which achieves high temperatures through microwave assistance to accelerate reaction times. However, we observed a pattern of obtaining surprisingly low synthesis yield (<10-40% for a typical 20-mer peptide) when using Dbz(Alloc) linker during the synthesis. This was unacceptable for our purposes.

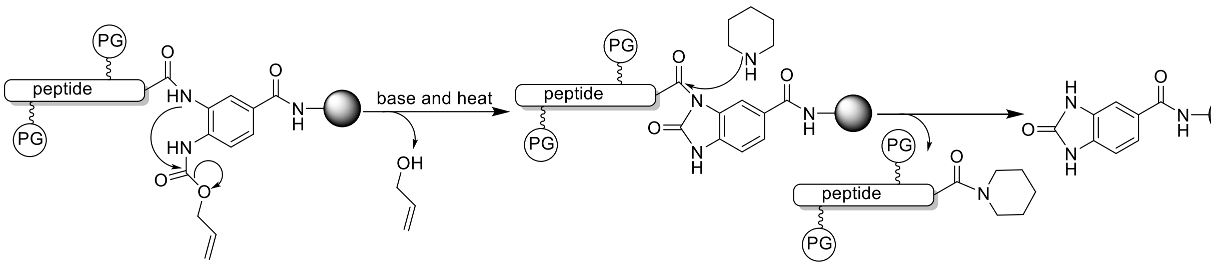

**Scheme S2**: Proposed mechanism for peptide release from resin with Fmoc deprotection condition under elevated temperature. (adapted from reference [5])

A major hint as to cause of yield loss came during synthesis of the "anchor peptide" prepared on Fmoc-Gly-Dbz(Alloc)-RGK(C) resin, which had an internal Dbz(Alloc) followed by a peptide sequence of sufficient size we could observe a major side product with a mass corresponding to Nbz-RGK(C) (Fig. S18A). This suggested internal cleavage at the Dbz(Alloc) under standard microwave synthesis conditions. With this in mind we revisited several syntheses carried out with Dbz(Alloc) linker with various C terminal sequence extensions, and after careful sample preparation and characterization, could identify similar products (Figure S18). Of note, the extent of side reaction was roughly correlated with the number of coupling cycles that had been carried out (compare for example Fig. S18A and Fig. S18E).

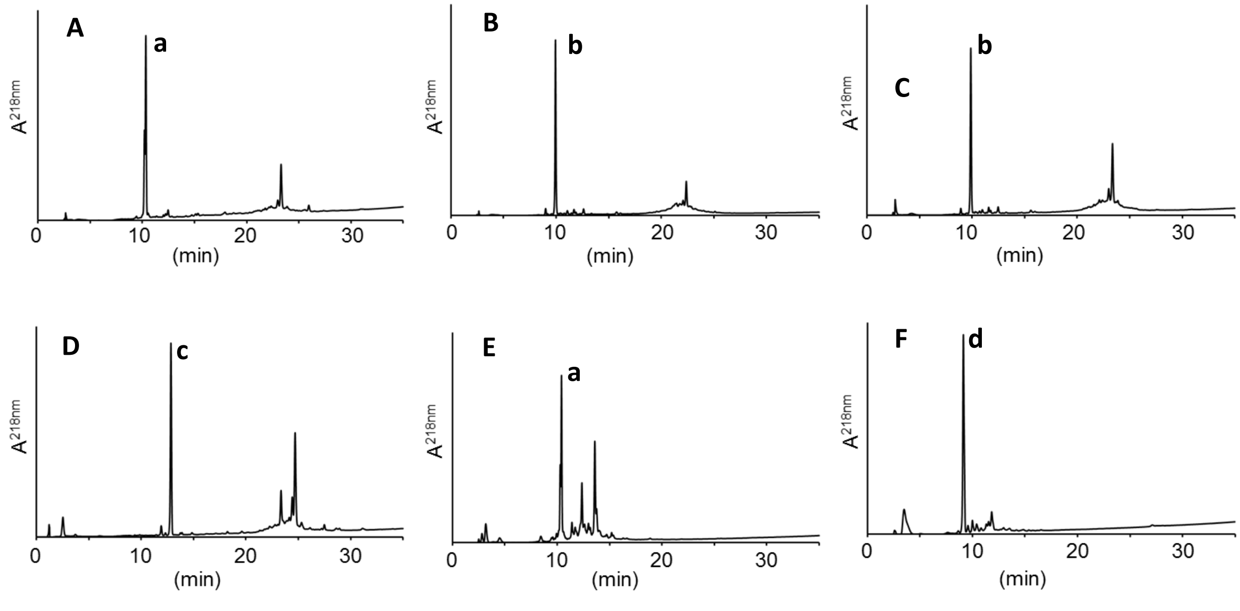

**Figure S18.** Example RP-HPLC traces of Dbz(Alloc)-containing peptides prepared under standard microwave conditions. For each, the relevant cleavage product is labeled. A: 27mer peptide synthesis on Dbz(Alloc)-RGK(C). B: 26mer peptide synthesis on Dbz(Alloc)-GK(C)G. C: 26mer peptide synthesis on Dbz(Alloc)-GK(C)G. D: 27mer peptide synthesis on Dbz(Alloc)-GK(Alloc). E: 9mer peptide synthesis on Dbz(Alloc)-RGK(C). F: 12mer peptide synthesis on Dbz(Alloc)-R. **a**=Nbz-RGK(C). **b**=Nbz-GK(C)G. **c**=Nbz-GK(Alloc). **d**=Nbz-R.

**4.2 Optimizing microwave SPPS on Dbz(Alloc) resin: temperature dependence of deprotection and coupling cycles**

To fully understand the parameters for Dbz(Alloc) cleavage and provide a feasible route for synthesis, we assessed the stability of Fmoc-Gly-Dbz(Alloc)-GK(Alloc) resin towards possible deprotection and activation cycles (summarized in Table S1).

In a typical experiment, ~10 mg of protected resin was incubated in 100 µL of the relevant reaction conditions (Table S1) and heated as described in a Bio-Rad T100™ Thermal Cycler. Resin was cleaved following standard protocols, and the products were characterized by RP-HPLC (Fig. S19) and MALDI-TOF MS. Please note that yields can only be roughly estimated from the RP-HPLC traces – the table reports integration of the 218nm absorbance, but Nbz- species typically have a larger extinction coefficient than Dbz(Alloc), such that this likely underestimates the Dbz(Alloc) species. However, since this is also dependent on the peptide sequence, we cannot quantify more accurately.

In standard microwave conditions, deprotection and coupling cycles are carried out at ~90 °C. We found that even at mildly elevated reaction temperatures (50 °C), all piperidine and piperazine treatment conditions (characteristic of Fmoc deprotection cycles) resulted in significant cleavage and loss of product (>50%) (Figure S19). However, we observed only minimal loss of product (~ 14%) after 24-hour treatment with 20% piperidine in DMF at room temperature. Since under manual or room temperature automated synthesis conditions each deprotection cycle holds these conditions for ~10 minutes, 24 hours would represent >100 coupling cycles – this explains why no significant yield loss was observed under standard SPPS conditions.

In contrast, standard activation/coupling conditions did not lead to significant yield loss. After 24 hours at elevated temperature (50 °C), we observed only ~10% cleavage and product loss in the presence of DIPEA as a typical activation/coupling solution base.

We therefore proposed that for Dbz(Alloc)-containing peptides, deprotection steps should be carried out at room temperature, while activation/coupling could be accelerated by microwave heating.

Table S1: 24h treatment of Dbz(Alloc) at various conditions

| Experiment # | Temperature (°C) | condition | Nbz (%) | Dbz(Alloc) (%) |
|---|---|---|---|---|
| A | 50 | 2% DBU, 24 hr | 90 | 10 |
| B | 50 | 20% piperidine, 24 hr | 72 | 28 |
| C | 50 | 10% piperazine, 24 hr | 48 | 52 |
| D | 50 | 20% piperidine, 24 hr | 14 | 86 |
| E | 25 | 10% DIEA, 24 hr | 10 | 90 |

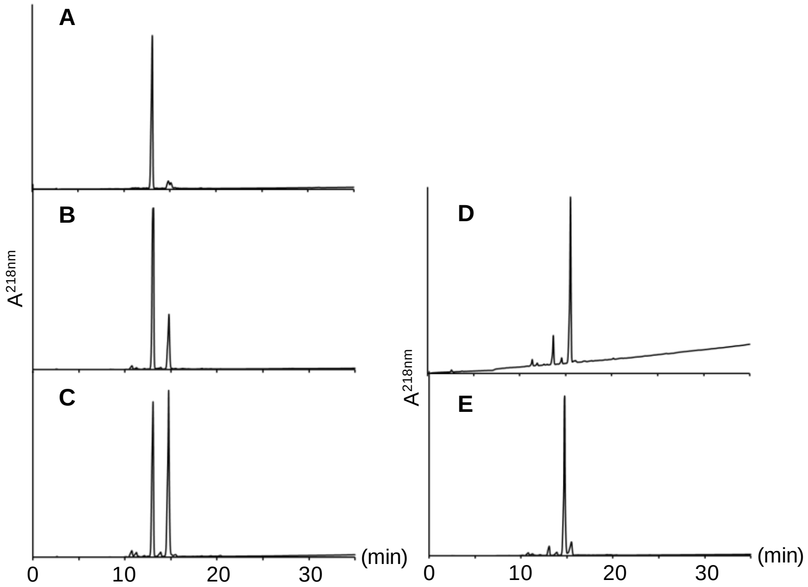

**Figure S19**: RP-HPLC traces of Nbz side product formation by treating Fmoc-Gly-Dbz(Alloc)-GK(Alloc) resin with various conditions. A: 2% DBU for 24h at 50 °C. B: 20% piperidine for 24h at 50 °C. C: 10% piperazine for 24h at 50 °C. D: 20% piperidine for 24h at room temperature. E: 20% DIEA for 24h at 50 °C.

To test these protocols in the context of a standard peptide, we assessed the synthesis of H1.2-($Thz_{67}$-$Gly_{99}$)-Dbz-GK(C). After manual addition of the $Gly_{99}$ to Dbz(Alloc) resin, 32 additional deprotection/coupling cycles were carried out as follows :
Deprotection: 20% piperidine in DMF at 25 °C, 2 x 5 min treatments
Coupling: 5:5:5 AA:DIC:Oxyma in DMF, 88 °C, 3 min.

We obtained 73% on-resin yield of the protected peptide (assessed by weight). We cleaved the resin and characterized products by RP-HPLC and confirmed peak identity with MALDI-TOF MS; the two major products were the hydrolyzed product and the desired peptide. Again, please note that the hydrolyzed Nbz-containing product would be expected to show significantly higher absorbance than the peptide-Dbz desired product, making quantitation challenging. However, assuming that all coupling steps through Gly were performed in 100% yield and all subsequent steps achieved 99% completion, we would expect a yield of $(0.99)^{32}$ = 72.5%. We therefore conclude that under these conditions, we have an outer bound of $< 1\%$ loss per coupling cycle, making these synthesis conditions appropriate for longer peptides.

Prior to publication, we found in the literature a report from Yoshiya and co-workers[5], who also observed cleavage using protected Dbz(Alloc) under microwave conditions. They developed an alternative Dbz($NO_2$) protected linker; however, this derivative can only be used for peptides with Gly as the C-terminal residue. Blanco-Canosa, Dawson, and coworkers also reported the methyl-protected MeDbz linker as a microwave-safe alternative; we also use this group extensively. However, there are several conditions under which MeDbz is nonideal (see also section 4.3), such that Dbz(Alloc) remains a valuable synthesis tool.

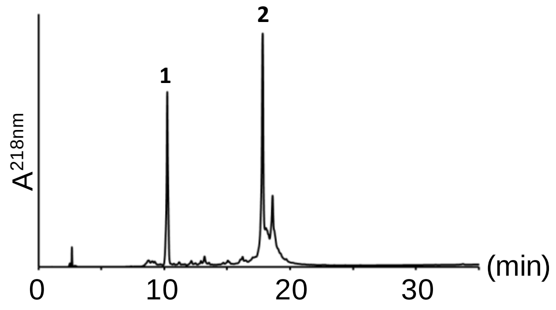

**Figure S20.** RP-HPLC trace of crude peptide H1.2-($Thz_{67}$-$Gly_{99}$)-Dbz-GK(C) synthesized with Dbz(Alloc) linker under modified coupling cycle (5min double Fmoc deprotection with 20% piperidine and 0.1M Oxyma at room temperature and 3min DIC/Oxyma coupling at 88 °C). **1**=Nbz-GK(C) calculated m/z: $[MH]^+$ 466.5 observed m/z: $[MH]^+$ **2=** H1.2-($Thz_{67}$-$Gly_{99}$)-Dbz-GK(C) calculated m/z: $[MH]^+$ 3900.6, observed m/z: $[MH]^+$ 3900.3.

##### 4.3 Benzimidazole formation from peptidyl MeDbz under acidic condition

When synthesizing peptides on Dbz linkers, we typically carry out a test cleavage and analysis prior to conversion to the active Nbz form, to enable us to separate side reactions in the synthesis component from side reactions of linker activation. When working with the MeDbz linker, we consistently observed an extraneous peak on RP-HPLC and MALDI-TOF analysis (Figure S21-A, S22-A). The mass of side product was 18 Da less than the expected peptidyl MeDbz, indicating a dehydration process.

We found that the dehydration process continued in HPLC buffer (0.1%TFA in water) (Figure S21-B) and could be inhibited by adjusting pH to 7 (Figure S21-C), suggesting an acid-catalyzed dehydration process. After a search of the literature, we hypothesized that this was due to acid-catalyzed cyclization of MeDbz to form peptidyl benzimidazole (Scheme S3). In fact, this reaction is sufficiently robust that recently the Dawson group reported the use of this reaction to prepare benzimidazole product, which might be promising in drug development[6] by treatment in neat TFA for 24 hours.

However, we observe that this side reaction is robust not just under cleavage conditions, but when peptides are stored under standard RP-HPLC conditions (0.1% aqueous TFA, with or without acetonitrile cosolvent), such that this peak is significant in all RP-HPLC analyses of these peptides. In fact, we find that the reaction appears more significant under aqueous conditions than neat TFA ((details in Table S2 and Figure S22B & S23B, S22C & S23C).

We find that the kinetics are dependent not only the C-terminal residue but also the peptide sequence (Figure S21, S22). We also found that the dehydration process is even faster under HPLC buffer condition than that under TFA cleavage condition (Figure S22 & S23). We carried out 2h, 4h, and 8 hours TFA cleavage and run the product immediately on RP-HPLC and observe that prolonged cleavage time will increase the dehydration product (Figure S22). However, if we adjust pH of the 2h cleavage product to pH 7, we observe no change after storage for 20h at room temperature (Figure S22A & S23A). Finally, after adjusting the pH of this product back to ~1 (RP-HPLC conditions), we observed increased dehydration product after 2h and 6h.

Table S2. Dehydration product ratio change under various conditions (based on HPLC traces from Figure S22 & S23).

| **Entries** | **Treatment conditions** | **MeDbz form ratio** | **Methylbenzimidazole form ratio** |
|---|---|---|---|
| **1** | 2h TFA cleavage | 80.9% | 19.1% |
| **2** | 4h TFA cleavage | 72.5% | 27.5% |
| **3** | 8h TFA cleavage | 59.3% | 40.7% |
| **4** | 20h at pH 7 from entry 1 | 80.9% | 19.1% |
| **5** | 2h at pH 1 from entry 4 | 64.4% | 35.6% |
| **6** | 6h at pH 1 from entry 4 | 47.8% | 52.2% |

Since we did not observe this -18 Da product in the case of Dbz, we attribute this to the extra methyl group on the 4-amine on MeDbz. We hypothesize that the presence of the methyl group causes steric hinderance for acylation during amino acid coupling step, thus avoiding the branched peptide side product. The recent paper reported that the methyl amine would be capped under traditional acetyl capping condition but would not by a larger benzoic acid, suggesting some impaction of the steric hinderance for acylation[7]. However, it increases the reactivity of the secondary amine with amide carbonyl group compared to that on Dbz under acidic conditions, most likely due to electron-donating effect of the methyl group.

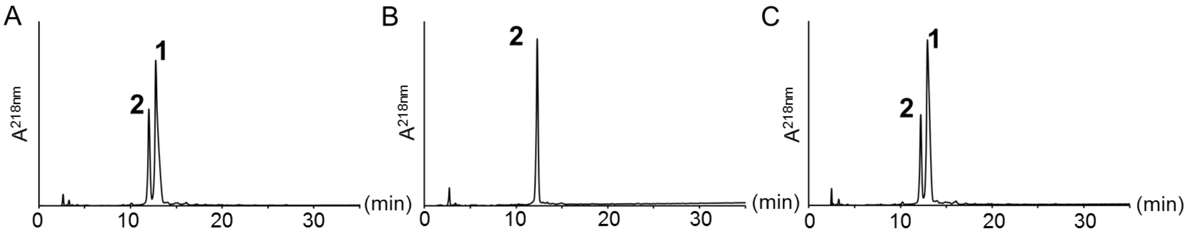

**Figure S21**: pH-controlled benzimidazole formation. A) 0h after 2 hours cleavage in TFA cleavage buffer (95%.TFA+ 2.5% $H_2O$+2.5%TIPS). B) 24h in HPLC buffer (0.1%TFA in $H_2O$, pH ~1). C) 24h in neutralized HPLC buffer (pH ~7). Peak **1**=H1.2-($Pro_{173}$-$Ala_{188}$)-MeDbz-$NH_2$, calculated m/z: $[MH]^+$ 1799.3, observed m/z: $[MH]^+$ 1799.4, peak **2**= H1.2-($Pro_{173}$-$Ala_{188}$)-methylbenzimidazole-$NH_2$ calculated m/z: $[MH]^+$ 1781.3, observed m/z: $[MH]^+$ 1781.3

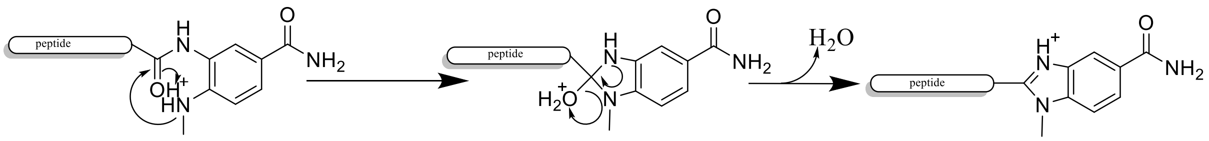

**Scheme S3**: Proposed acid-catalyzed mechanism for the generation of peptidyl benzimidazole from peptidyl MeDbz.

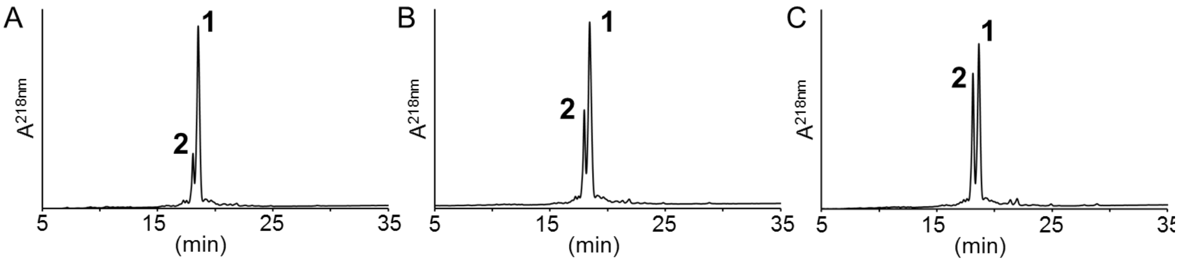

**Figure S22**: Benzimidazole formation at various timepoints of cleavage. A) 2 hours cleavage in TFA cleavage buffer (95%.TFA+ 2.5% $H_2O$+2.5%TIPS). B) 4 hours cleavage in TFA cleavage buffer. C) 8 hours cleavage in TFA cleavage buffer. Peak **1**=H1.2-($Ser_1$-$Ala_{23}$)-MeDbz-$NH_2$, calculated m/z: $[MH]^+$ 2349.8, observed m/z: $[MH]^+$ 2349.1, peak **2**= H1.2-($Ser_1$-$Ala_{23}$)-methylbenzimidazole-$NH_2$. calculated m/z: $[MH]^+$ 2331.8, observed m/z: $[MH]^+$ 2331.3

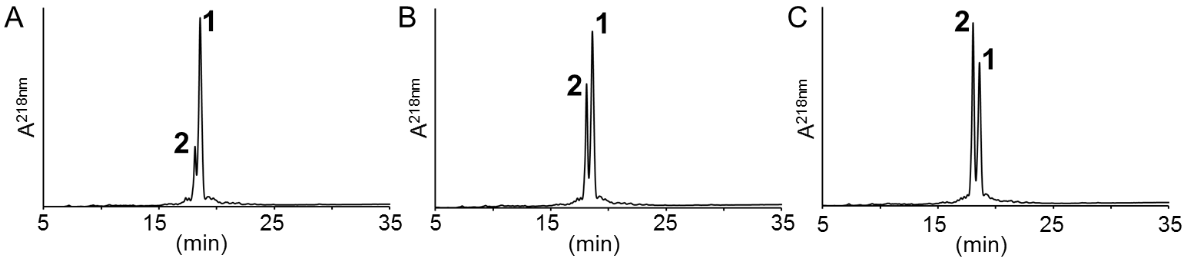

**Figure S23**: Benzimidazole formation in acidic aqueous solution. A) 20 hours sitting in neutralized HPLC buffer (pH ~7) after 2 hours of cleavage in TFA cleavage buffer (95%.TFA+ 2.5% $H_2O$+2.5%TIPS). B) 2 hours after pH adjusted to ~1 from A). C) 6 hours after pH adjusted to ~1 from A). Peak **1**=H1.2-($Ser_1$-$Ala_{23}$)-MeDbz-$NH_2$, calculated m/z: $[MH]^+$ 2349.8, observed m/z: $[MH]^+$ 2349.1, peak **2**= H1.2-($Ser_1$-$Ala_{23}$)-methylbenzimidazole-$NH_2$. calculated m/z: $[MH]^+$ 2331.8, observed m/z: $[MH]^+$ 2331.3

##### 4.4 Incomplete on-resin MeDbz cyclization to MeNbz

During the peptide synthesis, we observed incomplete on-resin MeDbz cyclization to MeNbz for several H1.2 peptides. Here we take the H1.2-($Thz_{49}$-$Ala_{66}$)-R53cit-MeDbz peptide as an example.

We initially used the conditions recommended for difficult peptide by Blanco-Canosa et al [3]. We treated the resin with 0.25M 4-NPCF in DCM for 1h, treatment with 15% DIEA for 1h. However, we observed significant amount of un-converted MeDbz peak and 4-nitrophenyl formyl adduct intermediate (Figure S24A). We next tested two cycles of 4-NPCF treatment but observed no significant differences. (Figure S24B).

We hypothesized that the incomplete addition and cyclization might result from reduced reactivity of the substituted amine. Therefore, we carried out the two steps of MeNbz conversion at elevated temperature (50 °C) and found improved conversion to peptidyl MeNbz (Figure 24C).

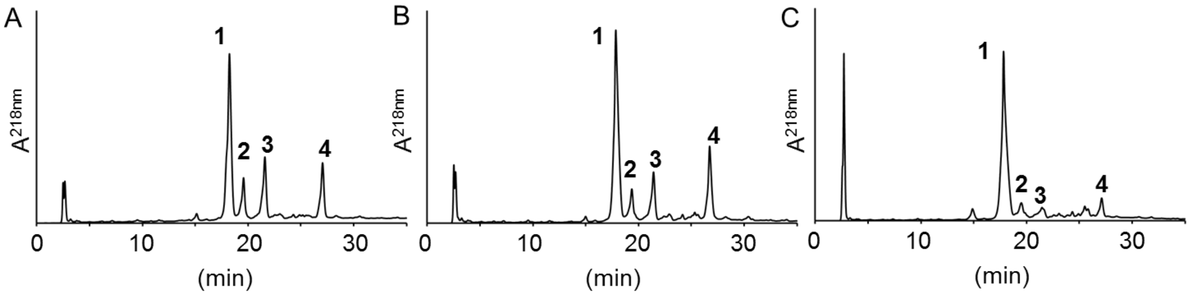

**Figure S24**: On-resin MeDbz cyclization to MeNbz with H1.2-($Thz_{49}$-$Ala_{66}$)-R53cit-MeDbz peptide at various conditions. A) 1h 0.25M 4-NPCF treatment and 1h 15% DIEA treatment. B) double 1h 0.25M 4-NPCF treatment and 1h 15% DIEA treatment. C) 1h 0.25M 4-NPCF treatment and 1h 15% DIEA treatment at 50 ºC. peak **1=** H1.2-($Thz_{49}$-$Ala_{66}$)-R53cit-MeNbz, calculated m/z: $[MH]^+$ 2019.4, observed m/z: $[MH]^+$ 2018.7, peak **2**= H1.2-($Thz_{49}$-$Ala_{66}$)-R53cit-methylbenzimidazole, calculated m/z: $[MH]^+$ 1975.4, observed m/z: $[MH]^+$ 1975.1, peak **3**= H1.2-($Thz_{49}$-$Ala_{66}$)-R53cit-MeDbz, calculated m/z: $[MH]^+$ 1993.4, observed m/z: $[MH]^+$ 1993.2, peak **4**= H1.2-($Thz_{49}$-$Ala_{66}$)-R53cit-MeDbz (4-nitrophenyl formyl). calculated m/z: $[MH]^+$ 2158.5, observed m/z: $[MH]^+$ 2157.4

### 5 H1.2 N and C terminal segment Synthesis by Solid Phase NCL

In this section, we provide the experimental protocols and details for the solid phase ligation.
**Sections 5.1-6** summarize general protocols for the key steps in solid phase ligation.
**Section 5.7** describes our initial attempt to prepare H1.2 solely by sequential solid phase ligation.
**Sections 5.8-5.11** provide the protocols and data for 4 repetitions of solid phase ligation to assemble the C-terminal protein segment (H1.2-(Cys100-Lys212)-OH, with and without S172ph.  Also note, section 5.9 & 5.10 describe our efforts to demonstrate that currently available commercial PEGA resins are suitable for solid phase ligation.
**Sections 5.12-5.14** provide protocols and data for 4 repetitions of solid phase ligation to assemble the N-terminal protein segment (H1.2-(Ser1-Gly99)-Dbz-x) with and without R53cit.

Our most significant recommendations, based on our results, are 1) to include an extra wash step with cysteine-contained wash buffer after each ligation, 2) use the methoxyamine approach for Thz ring opening, and 3) the resin cannot be stored in methanol or ethanol once the anchor peptide that contains the HMBA linker is attached, to avoid product loss due to cleavage of the ester linkage.

**Buffers:**
Wash Buffer: 0.1 M Phosphate, 6 M GuHCl, pH 6.5
Thz Ring Opening Buffer (RO Buffer): 0.1 M Phosphate, 0.4 M Methoxyamine, 6 M GuHCl, pH 4
Ligation Buffer: 0.1 M Phosphate, 0.05 M MPAA, 6 M GuHCl, 10mM TCEP, pH 7

##### 5.1 Fmoc Deprotection of Base Resin

200 µL of the base resin Fmoc-Thz-Ala-Ahx-Tyr-Lys-Gly-Rink-PL PEGA (section 2.3), swelled in methanol was measured in a 1.2 mL bed volume Bio-Spin Chromatography column (Bio-Rad). Based on the theoretical loading of 0.0025 mmol/mL, the scale was 0.5 µmol. The resin was treated with three repetitions of 20% Piperidine in NMP for 5 minutes. The resin was then washed with 15 column volumes of DMF, followed with 5 columns volumes of methanol, and then with 15 column volumes of water. The resin was nutated in water for 10 minutes. The resin was then flow-washed with 9 column volumes of Wash Buffer, and nutated in Wash Buffer for 5 minutes. The flow-wash/nutation step was repeated 3 more times.

##### 5.2 Thz Deprotection on Solid-Phase

Conversion of thiazolidine to cysteine was typically carried out in Thz Ring Opening Buffer (RO buffer). Resin was treated with 2-3 column volumes of buffer and nutated for 2 hours.  The column was drained, and the cycle repeated for a total of 3-4 cycles. The resin was flow-washed with 15 column volumes of wash buffer, then nutated in wash buffer for 5 minutes and drained.  This wash cycle was repeated x2. Finally wash buffer with 10mM TCEP was added, nutated for 10 minutes, drained, and flow-washed with 5 column volumes wash buffer, then nutated with ligation buffer for 5 min and drained prior to ligation.

Analysis of ring opening with microcleavage is recommended in order to avoid incomplete deprotection. Analysis of deprotection with minimal product loss is carried out as follows:  5-10 beads are washed with water, then cleaved in a minimal volume of TFA for 15-30 minutes with residual water acting as scavenger.  MALDI-TOF MS analysis is used to assess reaction completion.

Please note section 5.9, which details our exploration of alternate Thz deprotection systems, which we found to be less effective than the simple methoxylamine treatment.

**5.3 Thioester resin activation**

Diglycolic acid PEGA resin was activated with Thiophenol/DIC/DMAP in DMF (for 1mL reaction mixture 48µL DIC 30µL thiophenol, 10µL 0.2M DMAP were added). The activation was carried out twice, each for 30min. The activated resin was washed with DMF thoroughly and then with ethanol followed by water in a flow-wash/nutation manner. Flow-wash with 3 column volumes Wash Buffer, followed by 2-3 column volumes ligation buffer, and drain before addition of peptide in Ligation Buffer.

**5.4 Solid-Phase Native Chemical Ligation**

For a typical ligation cycle, after thorough washing (see 5.2), ~2-5 equivalents peptide was dissolved in Ligation Buffer, and reaction allowed to proceed with nutation for 4 hours. Ligation was assessed via microcleavage and MALDI-TOF MS, supplemented by RP-HPLC analysis of the peptide content of the supernatant liquid. If complete, resin was drained, and flow washed with 15 column volumes of Wash Buffer.

Occasional thioester or disulfide adducts were observed after ligation steps, presumably through side chain Cys thioester (see section 5.15). To revert these, an additional Cys wash step was added; 50 mM cysteine in ligation buffer was added to resin and nutated for 30 min at room temperature, followed by thorough washing (see 5.2 for typical cycle).

**5.5 Solid-Phase Desulfurization**

2-3 column volumes of desulfurization buffer (3 M Guanidine, 0.1 M phosphate, 400mM TCEP, 150 mM MESNA, pH 7.4) were added to the resin. The solution was thoroughly sparged with argon. Free radical initiator VA-044 was added (to 20 mM final concentration), and the mixture was heated to 42 °C in a water bath. Reaction proceeded until complete as assessed by MALDI-TOF MS; typically 4 hours. If reaction was incomplete after 4 hours, resin was drained, washed, and the cycle repeated.

**5.6 C-terminal segment product release: NaOH Cleavage at the HMBA linker**

H1.2 C-terminal segment was released from the resin by treatment with 0.01 M NaOH for 30 minutes. In a typical reaction, 500µL of 0.01M NaOH was added and nutated for 30 minutes at room temperature; pH was checked to ensure the reaction was carried out at pH 10.0. The column was drained, and 500µL of 0.01 M HCl was added and drained to neutralize the NaOH. 3x additional column volumes were added and drained into the cleavage. The resin was rinsed with water (stored separately), and finally extracted with TFA (stored separately) to ensure maximal product recovery. In the ligation/cleavages described here, product was solely located in the initial NaOH/HCl rinses. The final product was accessed by RP-HPLC and SDS-PAGE and lyophilized for further use.

**5.7 Test Synthesis of H1.2 through solid phase native chemical ligation**

To access the need for an alternative ligation strategy, we attempted a sequential solid phase ligation for preparation of full-length linker histone H1.2. We divided the protein into 9 segments. 150µL Fmoc-Thz-Ala-Ahx-Lys-Gly-Rink-PL PEGA resin was used (0.25µmol) for the test ligation. The following ligation steps were carried out as describe in section 5.2 and 5.4. Ring opening of dmThz was tripled (18h with 9 buffer exchange) as described in section 5.2. RP-HPLC, MALDI-TOF MS were used to monitor the ligation process. The ligation details were summarized in Table S3. We obtained relatively pure product H1.2-(dmThz$_{87}$-Lys$_{212}$) (~13kDa) after first five ligations (Figure S25A-E, lane e on SDS-PAGE S25G). However, in the next ligation, there was almost no ligated product observed after cleavage from the resin either accessed by RP-HPLC or SDS-PAGE (Figure S25F and lane f on SDS-PAGE G ). Qualitatively, we also observed a decrease of the product peak in the HPLC trace after 5 ligations, which migh suggest a elution problem as we observed before in the synthesis of CENP-A. We reasoned that the size limitation (100-150 residues) issue for solid phase ligations is common for histone proteins. Therefore, we developed the new convergent hybdrid-solid phase ligation strategy for preparation of large and challenging proteins, including the linker histone H1.2 here.

Table S3. Test ligation of H1.2 peptides on solid phase.

| Ligation Round | Peptide | MW (g/mol) | Peptide (mg) | Volume (mL) | Conc. (mM) | Molar Equivalent | Time (h) |
|---|---|---|---|---|---|---|---|
| 1 | H1.2-($Thz_{189}$-$Lys_{212}$)-HMBA-RG-Nbz | 3091 | 4.0 | 0.5 | 2.6 | 5.2 | 15 |
| 2 | H1.2-($Thz_{163}$-$Ala_{188}$)- Nbz | 2869 | 4.0 | 0.5 | 2.8 | 5.6 | 14 |
| 3 | H1.2-($Thz_{134}$-$Ala_{162}$)-Nbz | 3120 | 5.9 | 0.5 | 3.8 | 7.6 | 16 |
| 4 | H1.2-( $Thz_{111}$-$Ala_{133}$)-Nbz | 2438 | 4.0 | 0.5 | 3.3 | 6.6 | 18 |
| 5 | H1.2-( $dmThz_{87}$-$Ala_{110}$)-Nbz fmyl | 2639 | 5.8 | 0.5 | 4.4 | 8.8 | 15 |
| 6 | H1.2-($Thz_{67}$-$Leu_{86}$)-Nbz | 2380 | 4.0 | 0.5 | 3.3 | 6.7 | 22 |

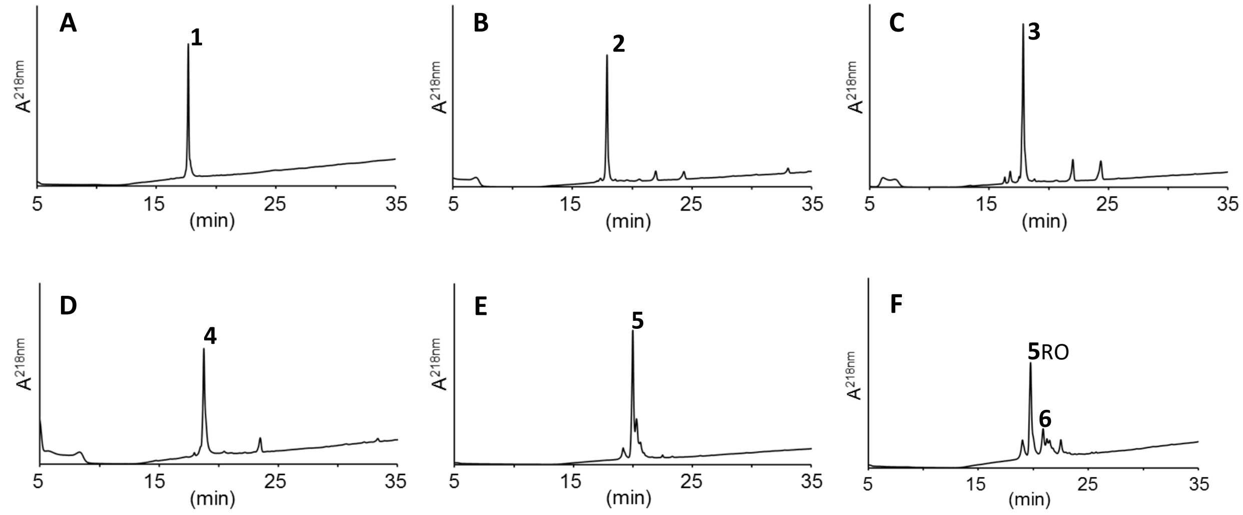

G

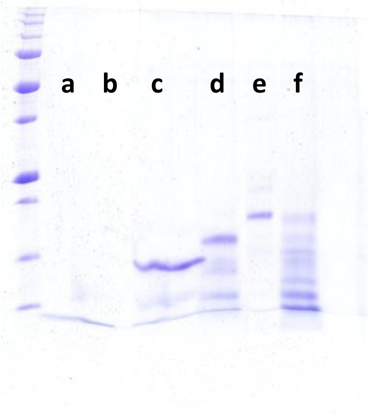

**Figure S25**. A-F: Crude RP-HPLC traces in a gradient 0-73% buffer B. A: crude after ligation of H1.2-($Thz_{189}$-$Lys_{212}$) peptide. B: crude after ligation of peptide H1.2-($Thz_{163}$-$Ala_{188}$). C: crude after ligation of peptide H1.2-($Thz_{134}$-$Ala_{162}$)-o. D: crude after ligation of peptide H1.2-($Thz_{111}$-$Ala_{133}$). E: crude after ligation of peptide H1.2-($dmThz_{87}$-$Ala_{110}$)-o. F: crude after ligation of peptide H1.2-($Thz_{67}$-$Leu_{86}$). G: monitoring of ligations by SDS-PAGE. **1**= H1.2-($Thz_{189}$-$Lys_{212}$)-o, **2**= H1.2-($Thz_{163}$-$Lys_{212}$)-o, **3**= H1.2-($Thz_{67}$-$Leu_{86}$)-o, **4**= H1.2-($Thz_{111}$-$Lys_{212}$)-o, **5**= H1.2-($dmThz_{87}$-$Lys_{212}$)-o, **6**= H1.2-($Thz_{67}$-$Lys_{212}$)-o, o= HMBA-RG-Cys-Ala-Ahx-Lys-Gly-$NH_2$. G: SDS-PAGE of ligations.

##### 5.8 Solid Phase Ligation of H1.2-($Cys_{100}$-$Lys_{212}$)-OH – First trial

To access the feasibility of the new convergent hybrid-solid phase ligation strategy, we first carried out the solid phase ligations for assembly of 4 C-terminal peptides. The ligation protocol used here was modified from previous hybrid phase ligation developed by our lab[2].

In our initial trial, 200µL of the Thz-Ala-Ahx-Tyr-Lys-Gly-Rink-PL-PEGA resin (0.5µmol, 300-500µm, no longer available) was used for the ligation. The following ligation steps were carried out as described in section 5.2 and 5.4. RP-HPLC, MALDI-TOF MS were used to monitor the ligation process (Figure S26, S27). Ligation went smoothly under the conditions we used. Cleavage of the peptide from resin was carried out as described in section 5.6. 4.0mg final peptide (77% overall yield or 93% yield for each ligation cycle after excluding the analysis loss) product was obtained in high purity, which would be directly used in further ligation. The ligation details were summarized in Table S4.

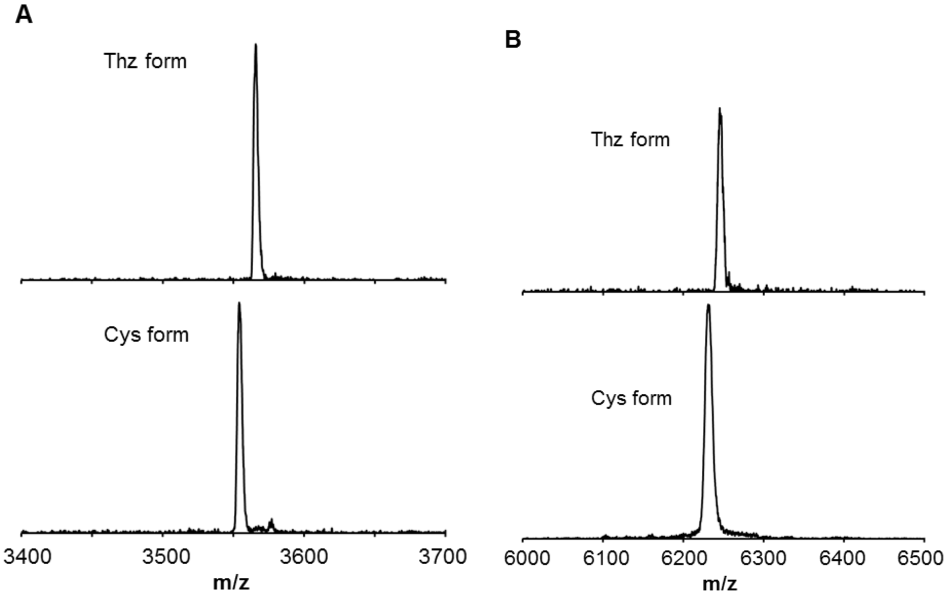

**Figure S26.** A typical MALDI-TOF MS analysis of the Thz ring opening reaction. Top: Thz form before ring opening. Bottom: Cys form after ring opening.

Table S4. Summary of H1.2-($Cys_{100}$-$Lys_{212}$)-OH ligations on solid phase.

| Ligation Round | Peptide | MW (g/mol) | Peptide (mg) | Volume (mL) | Conc. (mM) | Molar Equivalent | Time (h) |
|---|---|---|---|---|---|---|---|
| 1 | H1.2-($Thz_{189}$-$Lys_{212}$)-HMBA-RG-Nbz | 3091 | 6.0 | 0.7 | 2.7 | 3.8 | 16 |
| 2 | H1.2-($Thz_{163}$-$Ala_{188}$)- Nbz | 2869 | 6.0 | 0.7 | 3.0 | 4.2 | 12 |
| 3 | H1.2-($Thz_{134}$-$Ala_{162}$)-Nbz | 3120 | 6.0 | 0.7 | 2.5 | 3.8 | 13 |
| 4 | H1.2-( $Thz_{100}$-$Ala_{133}$)-MeNbz | 3585 | 7.2 | 0.7 | 2.7 | 4.0 | 16 |

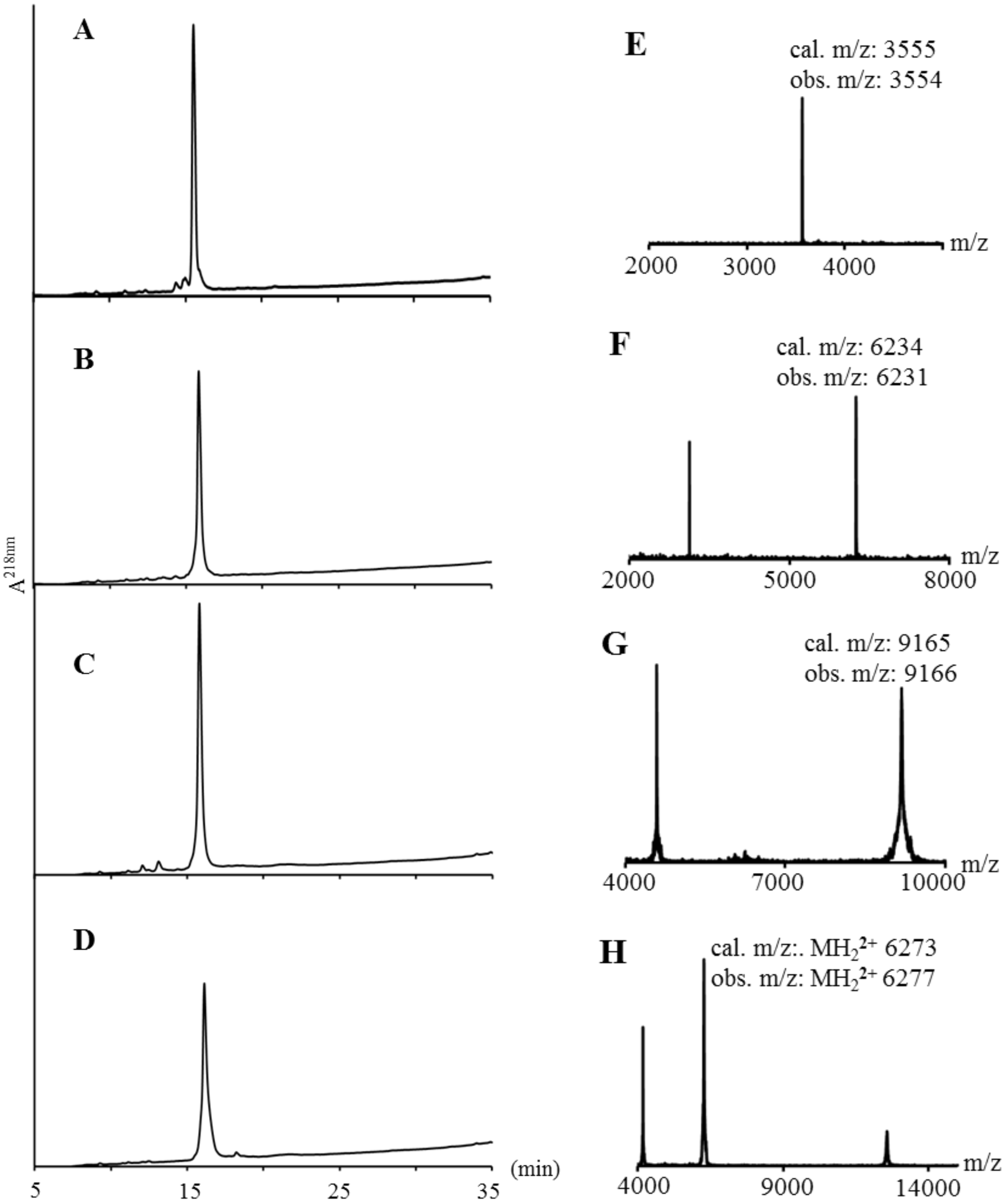

**Figure S27**: Crude RP-HPLC traces. A: H1.2-($Thz_{189}$-$Lys_{212}$)-o. B: H1.2-($Thz_{163}$-$Lys_{212}$)-O. C: H1.2-($Thz_{134}$-$Lys_{212}$)-o. D: H1.2-($Thz_{100}Lys_{212}$)-o. MALDI-TOF MS of E: H1.2-($Cys_{189}$-$Lys_{212}$)-o. F: H1.2-($Cys_{163}$-$Lys_{212}$)-o. G: H1.2-($Cys_{134}$-$Lys_{212}$)-o. H: H1.2-($Cys_{100}$-$Lys_{212}$)-o. o = HMBA-RG-Cys-Ala-Ahx-Tyr-Lys-Gly-$NH_2$.

##### 5.9 Solid Phase Ligation of H1.2-(Cys$_{100}$-Lys$_{212}$)-OH – Second trial

Since the 300-500µm PEGA resin (with pores of sufficient size for use with proteins < 70kDa in aqueous conditions) is no longer commercially available due to the acquisition of the manufacturer Varian by Agilent Technologies, we assessed solid phase ligation with a smaller bead resin in our second trial as a comparison. 200µL of the Fmoc-Thz-Ala-Ahx-Tyr-Lys-Gly-Rink PEGA resin (0.7µmol, 150-300µm, pores sufficiently for use with proteins <40kDa) was used for the ligation after Fmoc removal (section 5.1). We were initially concerned that the tighter mesh size would result in limited protein yield, considering the size limit of ~100 residues observed with the larger mesh PEGA beads.

The following ligation steps were carried out as describe in section 5.2 and 5.4. RP-HPLC, MALDI-TOF MS and SDS-PAGE were used to monitor or access the ligation process (Figure S28). Ligation went smoothly under the conditions we used. Cleavage of the peptide from resin was carried out as described in section 5.6. 5.6mg final peptide (81% overall yield or 95% yield for each ligation cycle) product was obtained in high purity, which will be directly used in further ligation. The ligation details were summarized in Table S5. This demonstrates that we observe no significant difference for solid phase ligation between the two different resins, such that the more common commercially available resin is suitable for use in our protocols.

Table S5. Summary of H1.2-(Cys$_{100}$-Lys$_{212}$)-OH ligations on solid phase.

| Ligation Round | Peptide | MW (g/mol) | Peptide (mg) | Volume (mL) | Conc. (mM) | Molar Equivalent | Time (h) |
|---|---|---|---|---|---|---|---|
| 1 | H1.2-(Thz$_{189}$-Lys$_{212}$)-HMBA-RG-Nbz | 3091 | 6.0 | 0.7 | 2.7 | 3.8 | 16 |
| 2 | H1.2-(Thz$_{163}$-Ala$_{188}$)- Nbz | 2869 | 6.0 | 0.7 | 3.0 | 4.2 | 12 |
| 3 | H1.2-(Thz$_{134}$-Ala$_{162}$)-Nbz | 3120 | 6.0 | 0.7 | 2.5 | 3.8 | 13 |
| 4 | H1.2-( Thz$_{100}$-Ala$_{133}$)-MeNbz | 3585 | 7.2 | 0.7 | 2.7 | 4.0 | 16 |

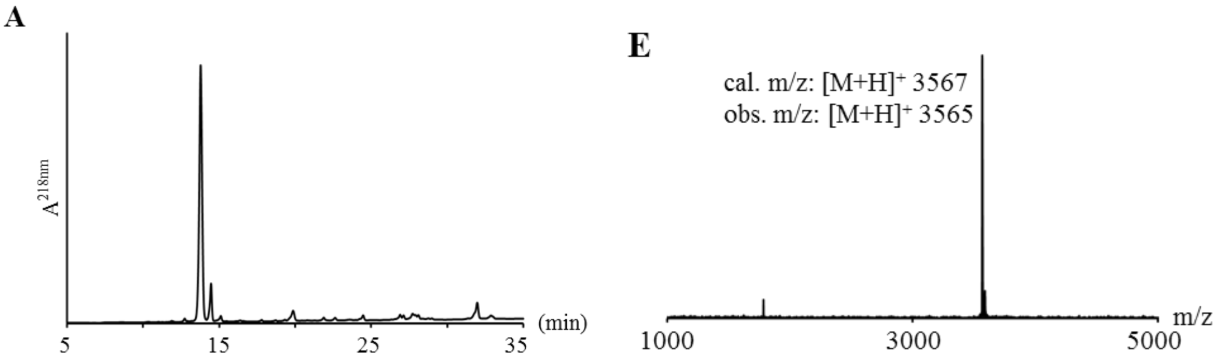

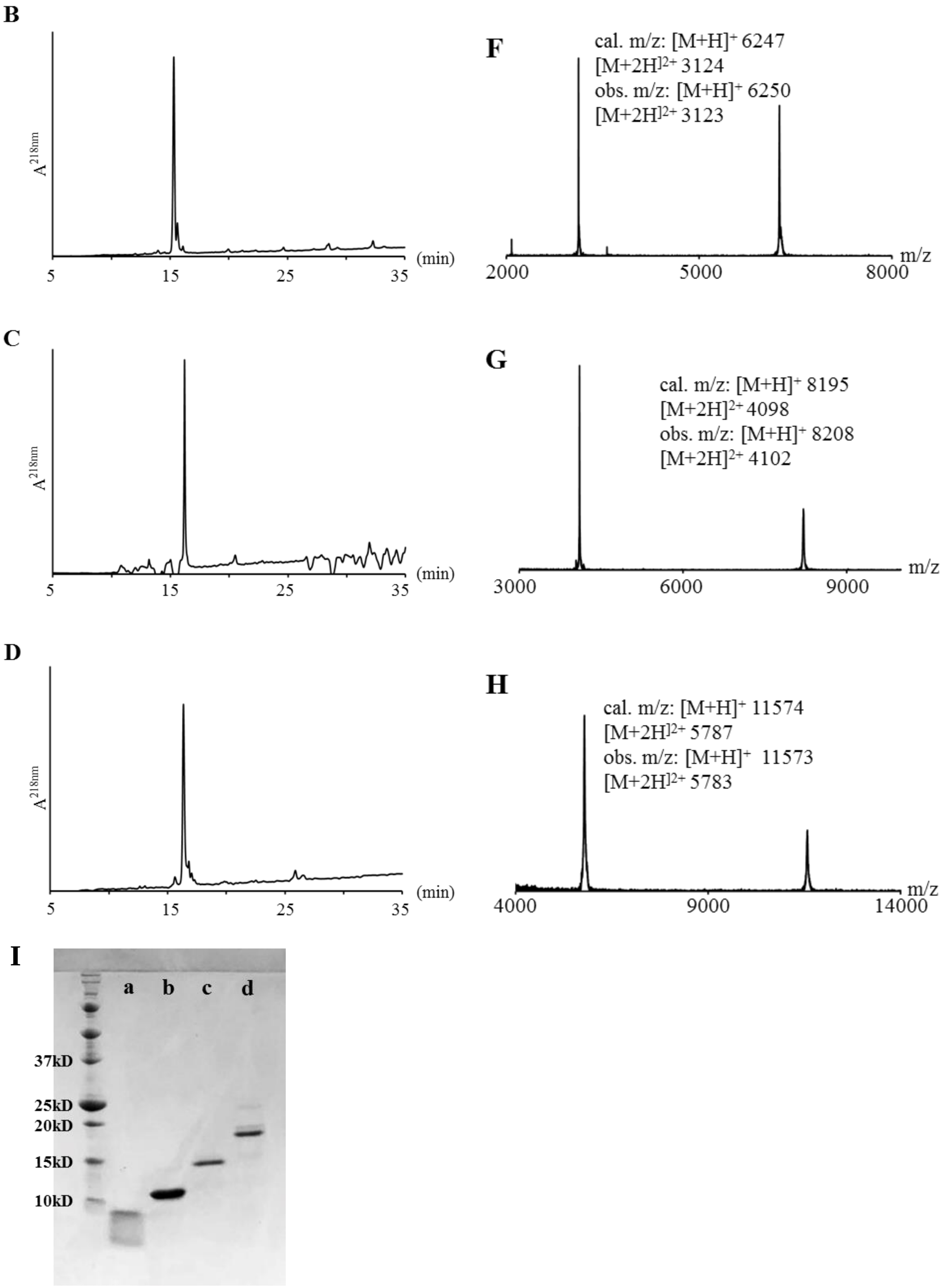

B
A218nm
(min)
5
15
25
35
C
A218nm
(min)
5
15
25
35
D
A218nm
(min)
5
15
25
35
F
cal. m/z: [M+H]+ 6247
[M+2H]2+ 3124
obs. m/z: [M+H]+ 6250
[M+2H]2+ 3123
m/z
2000
5000
8000
G
cal. m/z: [M+H]+ 8195
[M+2H]2+ 4098
obs. m/z: [M+H]+ 8208
[M+2H]2+ 4102
m/z
3000
6000
9000
H
cal. m/z: [M+H]+ 11574
[M+2H]2+ 5787
obs. m/z: [M+H]+ 11573
[M+2H]2+ 5783
m/z
4000
9000
14000
I
a
b
c
d
37kD
25kD
20kD
15kD
10kD

**Figure S28**. Crude RP-HPLC traces, MALDI-TOF MS and SDS-PAGE of H1.2-($Cys_{100}$-$Lys_{212}$)-OH solid phase ligation products. RP-HPLC gradient used: 0-50% buffer B over 30min. A: H1.2-($Thz_{189}$-$Lys_{212}$)-o. B: H1.2-($Thz_{163}$-$Lys_{212}$)-o. C: H1.2-($Thz_{134}$-$Lys_{212}$)-OH. D: H1.2-($Cys_{100}$-$Lys_{212}$)-OH. E: H1.2-($Thz_{189}$-$Lys_{212}$)-o. F: H1.2-($Thz_{163}$-$Lys_{212}$)-o. G: H1.2-($Thz_{134}$-$Lys_{212}$)-OH. H: H1.2-($Cys_{100}$-$Lys_{212}$)-o. I: SDS-PAGE of ligations. o = HMBA-RG-Cys-Ala-Ahx-Tyr-Lys-Gly-$NH_2$.

##### 5.10 Solid Phase Ligation of H1.2-($Cys_{100}$-$Lys_{212}$)-OH – Third trial

To further access the reproducibility and robustness of the solid phase ligation for preparation of C-terminal peptide segment, we carried out another trial. In this trial, 200µL of the Thz-Ala-Ahx-Tyr-Lys-Gly-Rink PEGA resin (0.5µmol, 150-300µm) was used for the ligation. The following ligation steps were carried out as described in section 5.2 and 5.4. RP-HPLC, MALDI-TOF MS were used to monitor the ligation process (Figure S29). Ligations went smoothly under the conditions as summarized in Table S6. Cleavage of the peptide from resin was carried out as described in section 5.6. 4.0mg final peptide (77% overall yield or 93% yield for each ligation cycle) product was obtained in high purity, which will be directly used in further ligation. So far, we demonstrated the robustness and reproducibility of the assembly of 4 H1.2 C-terminal peptides by solid phase ligation, two of which were prepared on the newer small-pore PEGA resin.

Table S6. Summary of H1.2-($Cys_{100}$-$Lys_{212}$)-OH ligations on solid phase.

| Ligation Round | Peptide | MW (g/mol) | Peptide (mg) | Volume (mL) | Conc. (mM) | Molar Equivalent | Time (h) |
|---|---|---|---|---|---|---|---|
| 1 | H1.2-($Thz_{189}$-$Lys_{212}$)-HMBA-RG-MeNbz | 4105 | 6.0 | 0.7 | 2.7 | 3.8 | 16 |
| 2 | H1.2-($Thz_{163}$-$Ala_{188}$)- MeNbz | 2883 | 6.0 | 0.7 | 3.0 | 4.2 | 12 |
| 3 | H1.2-($Thz_{134}$-$Ala_{162}$)-MeNbz | 3134 | 4.0 | 0.7 | 1.8 | 2.6 | 24 |
| 4 | H1.2-( $Thz_{100}$-$Ala_{133}$)-MeNbz | 3585 | 5.3 | 0.7 | 2.1 | 3.0 | 16 |

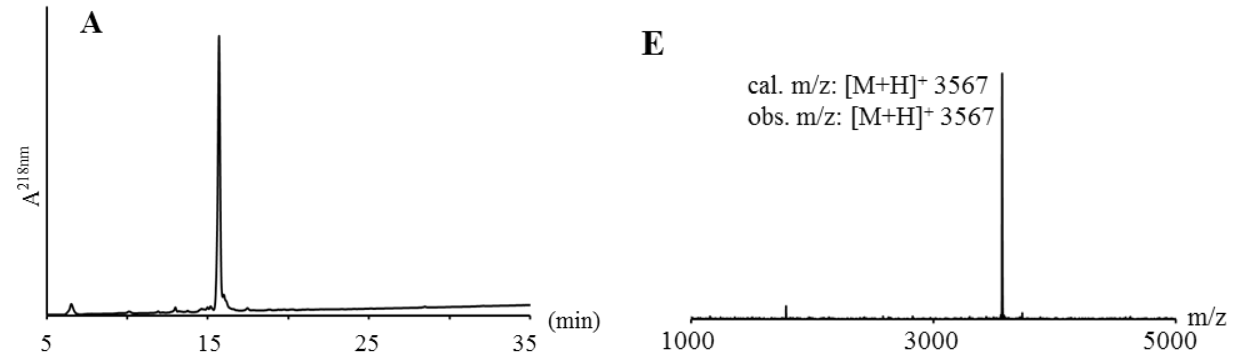

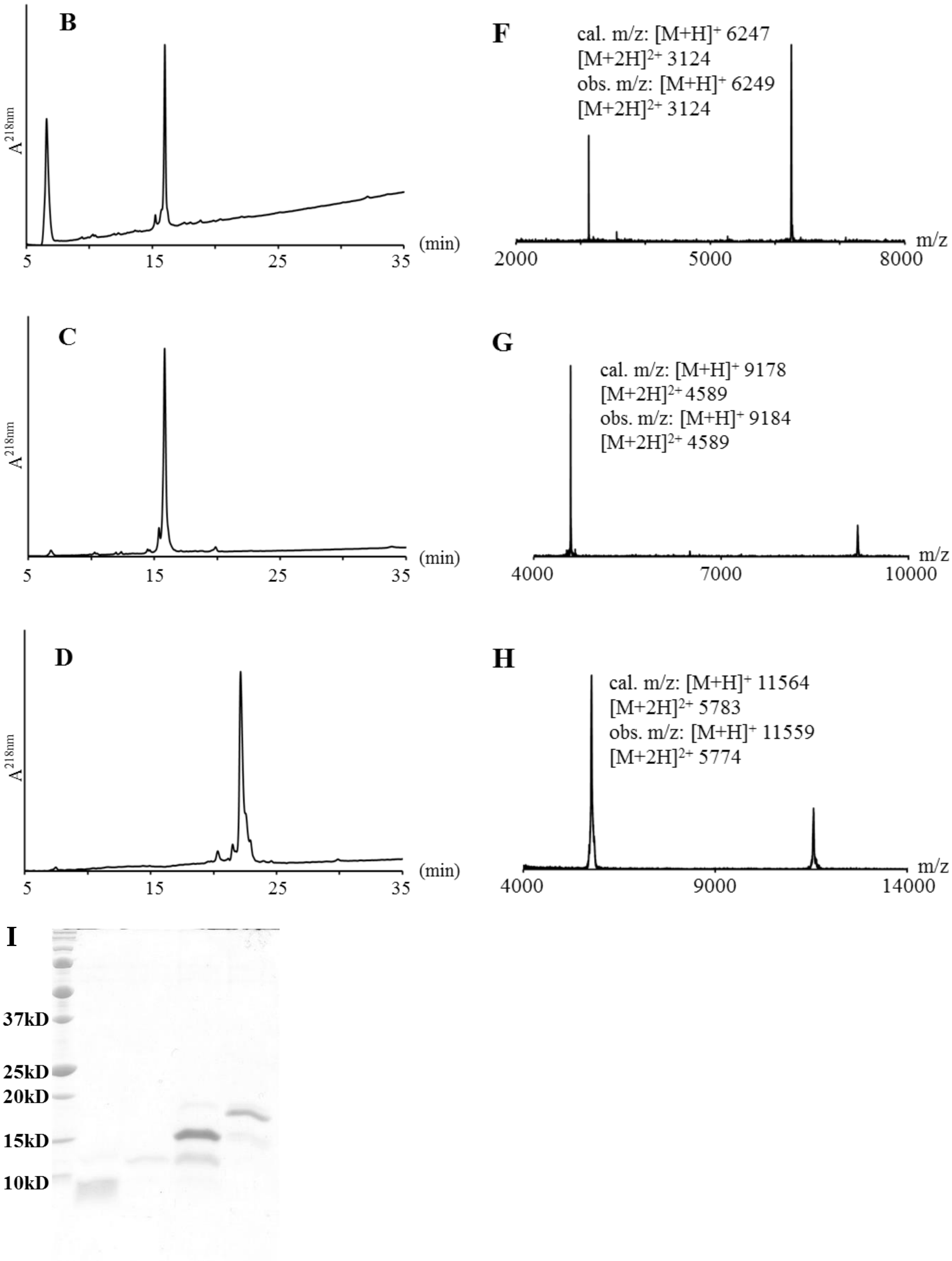

**Figure S29**: Crude RP-HPLC traces. A: H1.2-($Thz_{189}$-$Lys_{212}$)-o, 0-50% buffer B over 30min. B: H1.2-($Thz_{163}$-$Lys_{212}$)-o, 0-50% buffer B over 30min. C: H1.2-($Thz_{134}$-$Lys_{212}$)-o, 0-50% buffer B over 30min. D: H1.2-($Thz_{100}Lys_{212}$)-OH, 5-25% buffer B over 30min. MALDI-TOF MS of E: H1.2-($Cys_{189}$-$Lys_{212}$)-o. F: H1.2-($Cys_{163}$-$Lys_{212}$)-o. G: H1.2-($Cys_{134}$-$Lys_{212}$)-o. H: H1.2-($Cys_{100}$-$Lys_{212}$)-o. I: SDS-PAGE monitoring of the ligations. o = HMBA-RG-Cys-Ala-Ahx-Tyr-Lys-Gly-$NH_2$.

#### 5.11 Solid Phase Ligation of H1.2-($Cys_{100}$-$Lys_{212}$)-S172ph-OH

After demonstrating the feasibility of the solid phase ligation for preparation of unmodified H1.2 C-terminal peptide fragment, we moved forward to synthesize the phosphorylated H1.2 C-terminal peptide fragment. In this trial, 200uLof the Thz-Ala-Ahx-Tyr-Lys-Gly-Rink PL-PEGA resin was used (~0.5µmol, 300-500µm) for ligations.

During this trial, we tested the Pd-catalyzed Thz ring opening procedure recently developed by Brik group[8]. A modified procedure was followed on solid phase: 500uL 100mM $Pd[(Allyl)Cl]_2$ in ligation buffer was added to drained resin and incubated for 30min at 37 $^{o}$C, cycle repeated. Resin was thoroughly washed with ligation buffer until all the color was gone. The resin was next treated with 50mM DTT in wash buffer for 10min and cycle repeated. Repeated the above steps until the reaction is complete by MALDI-TOF MS check. We found that it was difficult to confirm/effect the complete removal of Pd catalyst during resin wash steps, although we did observe the desired color change.

All ligations were carried out as described in section 5.4. Ligation details were summarized in Table S7. Double ligation was carried out when necessary. 3.2mg crude peptide (73% overall yield) was obtained in relatively high purity, which will be directly used in following ligations. RP-HPLC, MALDI-TOF MS were used to monitor the ligation process and ligation product was accessed by SDS-PAGE (Figure S30, S31).

Overall, the yields and purity of this set of solid phase ligations were the lowest of any of our repeated syntheses. While it is hypothetically possible that the phosphorylated Ser caused some unknown challenges to the ligation, we attribute these lower yields to the Thz deprotection approach. A recent literature report additionally found that in at least one other system, $Pd[(Allyl)Cl]_2$ was less effective as a Thz deprotection agent, although little detail was provided[9].

Therefore, we recommend methoxyamine as Thz ring opening reagent in solid phase ligation – while $Pd[(Allyl)Cl]_2$ did work rapidly in solution phase tests, we found it less optimal in repeated solid phase use.

Table S7. Summary of H1.2-($Cys_{100}$-$Lys_{212}$)-S172ph-OH ligations on solid phase.

| Ligation Round | Peptide | MW (g/mol) | Peptide (mg) | Volume (mL) | Conc. (mM) | Molar Equivalent | Time (h) |
|---|---|---|---|---|---|---|---|
| 1 | H1.2-($Thz_{189}$-$Lys_{212}$) | 3091 | 3.0 | 0.5 | 2.0 | 2.5 | 16 |
| 2a | H1.2-($Thz_{163}$-$Ala_{188}$)-S172ph | 2961 | 2.9 | 0.5 | 2.0 | 2.5 | 8 |
| 2b | H1.2-($Thz_{163}$-$Ala_{188}$)-S172ph | 2961 | 3.7 | 0.5 | 2.4 | 3.0 | 16 |
| 3a | H1.2-($Thz_{134}$-$Ala_{162}$) | 3118 | 3.0 | 0.5 | 2.0 | 2.5 | 16 |
| 3b | H1.2-($Thz_{134}$-$Ala_{162}$) | 3118 | 3.0 | 0.5 | 2.0 | 2.5 | 16 |
| 4 | H1.2-($Thz_{100}$-$Ala_{133}$) | 3584 | 4.1 | 0.5 | 2.2 | 2.8 | 16 |

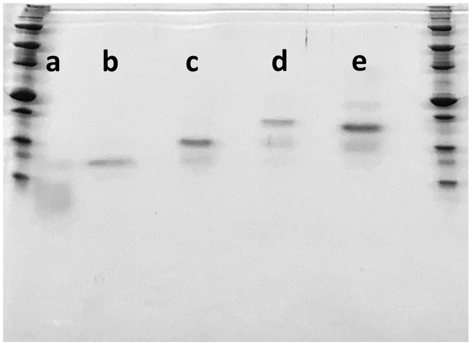

**Figure S30**: SDS-PAGE monitoring of H1.2-($Cys_{100}$-$Lys_{212}$)-S172ph ligations. lane a-d, products after each round of ligation cleaved by TFA. lane e, final product cleaved by NaOH.

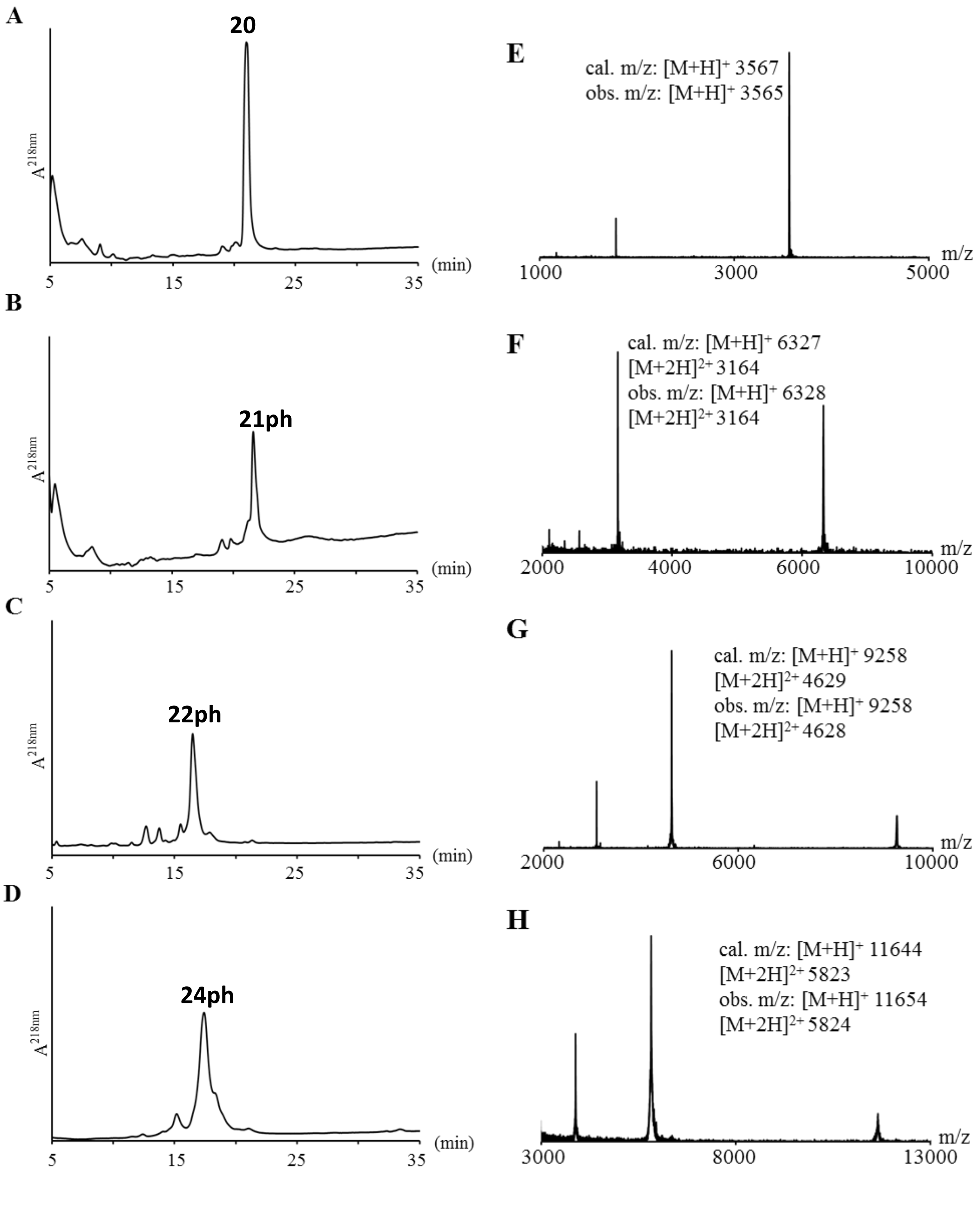

**Figure S31**: Crude RP-HPLC traces and corresponding MALDI-TOF MS. A: H1.2-(Thz$_{189}$-Lys$_{212}$)-o, 0-30% buffer B over 30min. B: H1.2-(Thz$_{163}$-Lys$_{212}$)-S172ph-o, 0-30% buffer B over 30min. C: H1.2-(Thz$_{134}$-Lys$_{212}$)-S172ph-o, 10-25% buffer B over 30min. D: H1.2-(Cys$_{100}$Lys$_{212}$)-S172ph-OH, 10-25% buffer B over 30min. o = HMBA-RG-Cys-Ala-Ahx-Tyr-Lys-Gly-$NH_2$. E: **20**. F: **21ph**. G: **22ph** H: **24ph**.

##### 5.12 Solid Phase Ligation of H1.2-($Ser_1$-$Gly_{99}$)-Dbz-x

To access the feasibility of the assembly of H1.2 N-terminal peptides by solid phase ligation, 200µL of the diglycolic acid-Rink PL-PEGA resin (~2.4µmol) was activated by thiophenol treatment in the presence of DIC and DMAP (section 5.3). 2.6mg H1.2-($Thz_{67}$-$Gly_{99}$)-Dbz-RGK(C) (0.6µmol) peptide was dissolved in 500µL ligation buffer and added to the drained resin. The capture reaction process was monitored by RP-HPLC. After 20h ligation, about 50% peptide was captured by the thioester resin. The resin was washed with Wash Buffer at pH 10 twice, each for 15min to hydrolyze the remaining thioester. The following ligation steps were carried out as described in section 5.2 and 5.4. We didn't observe any cyclized anchor peptide product in following analysis, suggesting complete hydrolysis of the remaining thioester by base treatment. Desulfurization was carried out on solid phase (section 5.5) after the last ligation. Peptide was cleaved from resin by TFA and precipitated by cold diethyl ether after concentrated by nitrogen flow. 1.5mg final peptide product was obtained in high purity, which will be directly used in further ligation. Overall yield of 72% was achieved after counting the loss during analysis steps. RP-HPLC and MALDI-TOF MS were used to monitor the ligation process (Figure S32). The details were summarized in Table S8.

Table S8. Summary of H1.2-($Ser_1$-$Gly_{99}$)-Dbz-x ligations on solid phase.

| Ligation Round | Peptide | MW (g/mol) | Peptide (mg) | Volume (mL) | Conc. (mM) | Molar Equivalent | Time (h) |
|---|---|---|---|---|---|---|---|
| 1 | H1.2-($Thz_{67}$-$Gly_{99}$)-Dbz-RGK(C) | 4055 | 2.6 | 0.7 | 0.9 | | 20 |
| 2 | H1.2-($Thz_{49}$-$Ala_{66}$)-Nbz | 2003 | 2.0 | 0.7 | 1.4 | 3.3 | 16 |
| 3 | H1.2-($Thz_{24}$-$Ala_{48}$)-Nbz | 2638 | 2.9 | 0.7 | 1.6 | 3.7 | 16 |
| 4 | H1.2-($Ser_1$-$Ala_{23}$)-Dbz-R | 2492 | 2.9 | 0.5 | 2.3 | 4.0 | 16 |

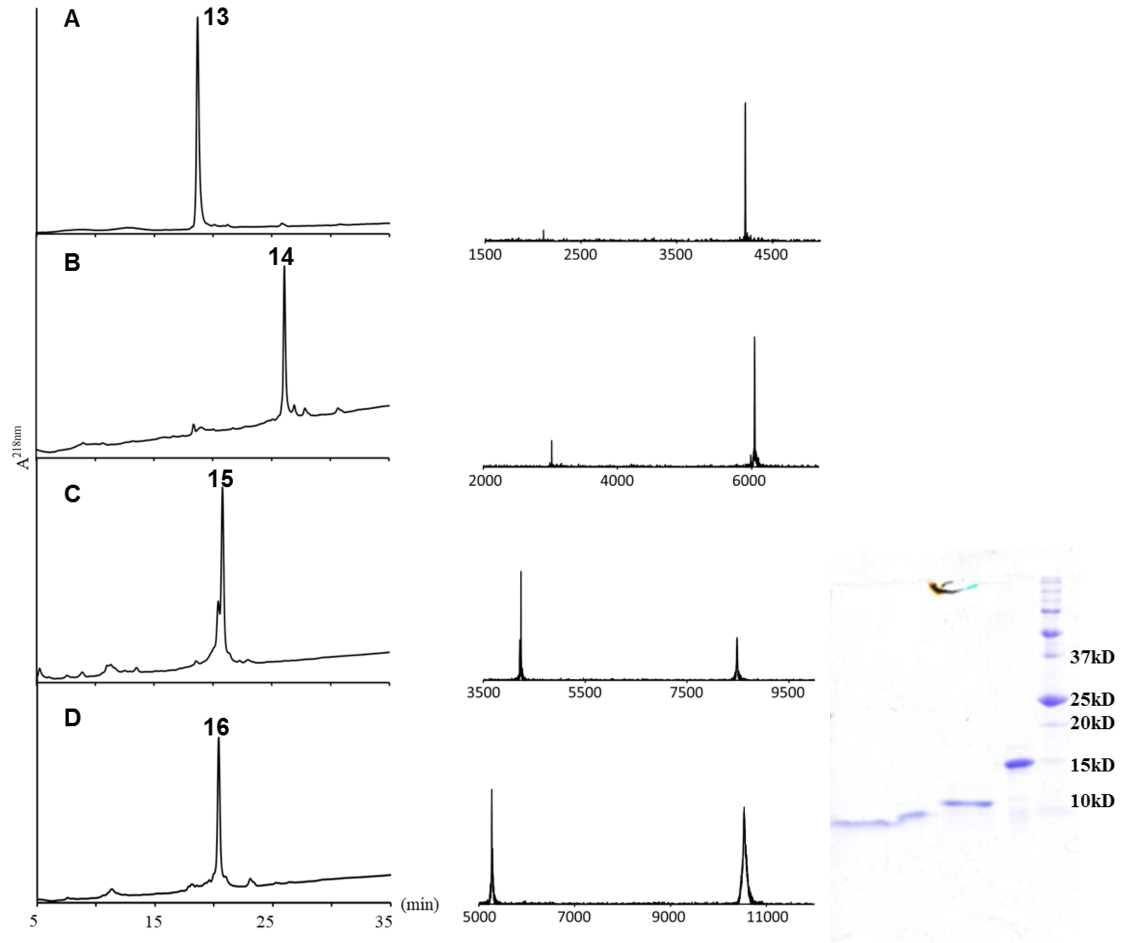

**Figure S32**: Crude RP-HPLC (left panel), MALDI-TOF MS (middle panel), and SDS-PAGE (right panel) monitoring of H1.2-($Ser_1$-$Gly_{99}$) assembly on solid phase. A: H1.2-($Cys_{67}$-$Gly_{99}$)-Dbz-x. calculated m/z: $[M+H]^+$ 4229, $[M+2H]^{2+}$ 2115. observed m/z: $[M+H]^+$ 4227, $[M+2H]^{2+}$ 2114. B: H1.2-($Thz_{49}$-$Gly_{99}$)-Dbz-x. calculated m/z: $[M+H]^+$ 6044, $[M+2H]^{2+}$ 3022. observed m/z: $[M+H]^+$ 6046, $[M+2H]^{2+}$ 3022. C: H1.2-( $Thz_{24}$-$Gly_{99}$)-Dbz-x. calculated m/z: $[M+H]^+$ 8493, $[M+2H]^{2+}$ 4247. observed m/z: $[M+H]^+$ 8494, $[M+2H]^{2+}$ 4247. D: after desulfurization H1.2-($Ser_1$-$Gly_{99}$)-Dbz-x'. calculated m/z: $[M+H]^+$ 10536, $[M+2H]^{2+}$ 5269. observed m/z: $[M+H]^+$ 10543, $[M+2H]^{2+}$5272. x=GK (C-diclycolic acid-Gly-$NH_2$), x'=GK (A-diclycolic acid-Gly-$NH_2$)

**5.13 Solid Phase Ligation of H1.2-($Ser_1$-$Gly_{99}$)-R53cit-Dbz-x - First Trial**

After demonstrating the feasibility of the preparation of unmodified H1.2 N-terminal peptide fragment, we moved onto the synthesis of citrullinated H1.2 N-terminal peptide. In the first trial, 200μL diglycolic acid-Ala-Ahx-Lys-Gly-Rink PEGA resin (~0.9μmol) was activated by treatment of thiophenol in the presence of DIC and DMAP (section5.3). 4.0mg H1.2-($Thz_{67}$-$Gly_{99}$)-Dbz-GK(C) (1.3μmol) peptide was dissolved in 400μL ligation buffer and added to the drained resin. Benzamidine was added to a final concentration of 25mM as an internal reference. The capture reaction process was monitored by RP-HPLC. After 16h ligation, about 70% peptide was captured by the thioester resin. Although the base treatment gives us satisfied results for thioester capping as described in section 5.12, a milder condition (for instance, neutral condition) would better be expected. The resin was washed with MPAA ligation buffer containing 100-200mM cysteine twice, each for 30min, to remove any remaining thioester. The following ligation steps were carried out as described in section 5.2 and 5.4. No cyclized product was observed in later analysis, indicating the success of Cys wash for thioester capping.

Desulfurization was carried out on solid phase (section 5.5) after the last ligation, since the N-terminal assembled segments contain no necessary Cys residues. The peptide was cleaved from resin by TFA and precipitated by cold diethyl ether after concentrated by nitrogen flow. 3.6mg of final peptide product was obtained in high purity, which will be directly used in further ligation. Overall yield of 59% was achieved after counting the loss during analysis steps. RP-HPLC, MALDI-TOF MS, and SDS-PAGE were used to monitor the ligation process (Figure S33). The details were summarized in Table S9.

Table S9. Summary of H1.2-($Ser_1$-$Gly_{99}$)-R53cit-Dbz-x ligations on solid phase.

| Ligation Round | Peptide | MW (g/mol) | Peptide (mg) | Volume (mL) | Conc. (mM) | Molar Equivalent | Time (h) |
|---|---|---|---|---|---|---|---|
| 1 | H1.2-($Thz_{67}$-$Gly_{99}$)-Dbz-GK(C) | 3091 | 4.0 | 0.7 | 1.8 | | 16 |
| 2 | H1.2-($Thz_{49}$-$Ala_{66}$)-R53cit-MeNbz | 2018 | 5.2 | 0.5 | 5.1 | 5.1 | 14 |
| 3 | H1.2-($Thz_{24}$-$Ala_{48}$)-Nbz | 2638 | 4.0 | 0.6 | 2.5 | 3.0 | 16 |
| 4 | H1.2-($Ser_1$-$Ala_{23}$)-Dbz-R | 2492 | 6.0 | 0.5 | 4.8 | 4.8 | 24 |

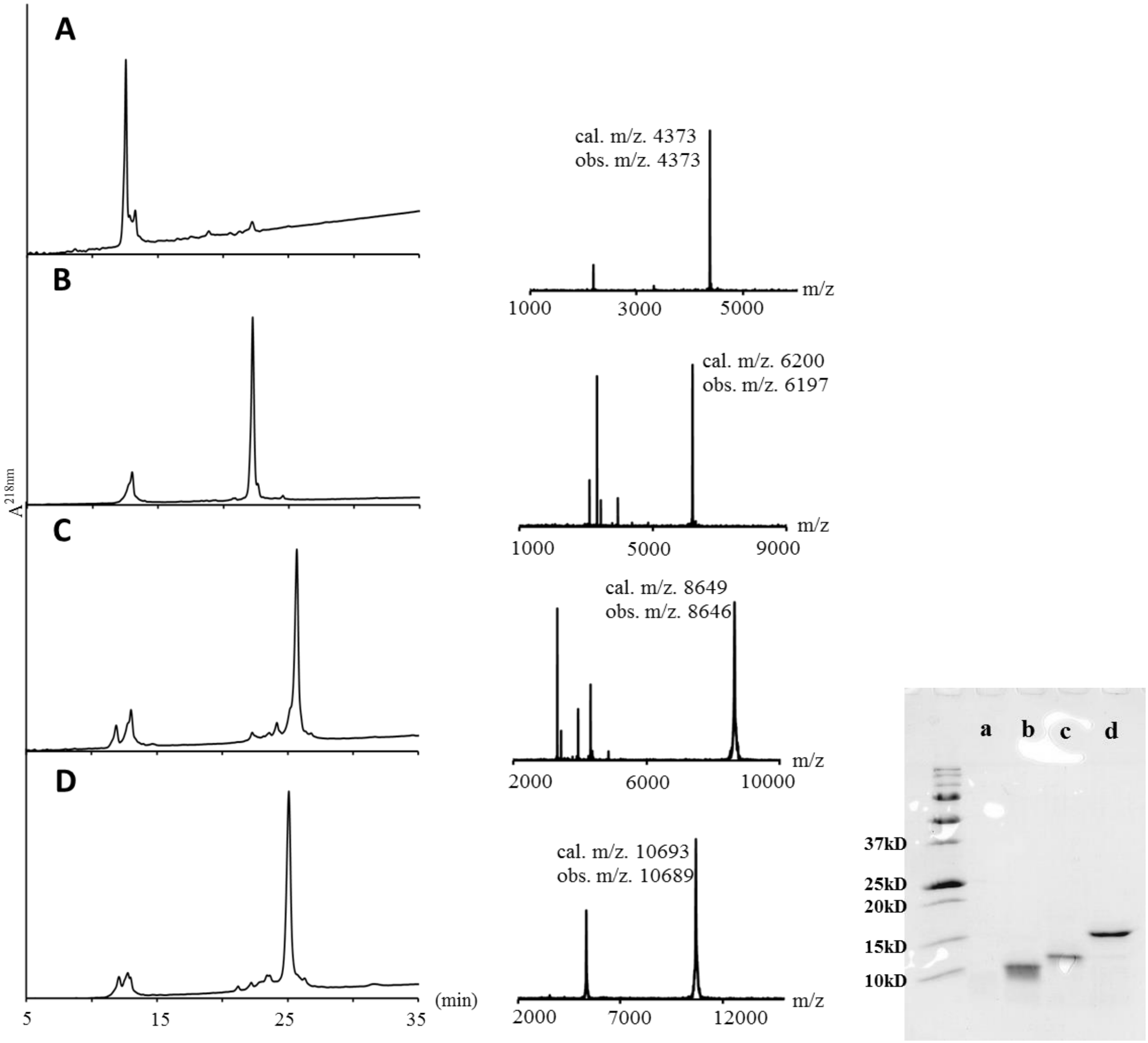

**Figure S33**: Crude RP-HPLC (left panel), MALDI-TOF MS (middle panel) and SDS-PAGE monitoring of H1.2-($Ser_1$-$Gly_{99}$)-R53cit assembly on solid phase. A: H1.2-($Cys_{67}$-$Gly_{99}$)-Dbz-x. B: H1.2-($Thz_{49}$-$Gly_{99}$)-R53cit-Dbz-x. C: H1.2-( $Thz_{24}$-$Gly_{99}$)-R53cit-Dbz-x. D: after desulfurization H1.2-($Ser_1$-$Gly_{99}$)-R53cit-Dbz-x'. x=GK (C-diclycolic acid-Ala-Ahx-Lys-Gly-$NH_2$), x'=GK (A-diclycolic acid-Ala-Ahx-Lys-Gly-$NH_2$). a-A, b-B, c-C, d-D.

**5.14 Solid Phase Ligation of H1.2-(Ser$_1$-Gly$_{99}$)-R53cit-Dbz-x - Second Trial**

In the second trial,400µLof diglycolic acid-Gly-Rink PEGA resin (4umol) was activated by treatment of thiophenol in the presence of DIC and DMAP (section5.3). 5.0mg H1.2-(Thz$_{67}$-Gly$_{99}$)-Dbz-GK(C) (1.6µmol) peptide was dissolved in 400µL ligation buffer and added to the drained resin. Benzamidine was added to a final concentration of 25mM as an internal reference. The capture reaction process was monitored by RP-HPLC. After 16h ligation, about 50% peptide was captured by the thioester resin. Cys wash was carried out as described in section 5.13 to cap remaining thioester. The following ligation steps were carried out as described in section 5.2 and 5.4. Desulfurization was carried out on solid phase (section 5.5) after the last ligation. Peptide was cleaved from resin by TFA and precipitated by cold diethyl ether after concentrated by nitrogen flow. 2.5mg of final peptide product was obtained in high purity, which will be directly used in further ligation. Overall yield of 50% was achieved after taking the loss during analysis steps into consideration. RP-HPLC, MALDI-TOF MS, and SDS-PAGE were used to monitor the ligation process (Figure S34). The details were summarized in Table S10.

In this trial, we tested shorter ligation time for two rounds of ligations (Table S10) and found that the ligation is compatible with these reduced time scales when high concentration of peptides is used.

Table S10. Summary of H1.2-(Ser$_1$-Gly$_{99}$)-R53cit-Dbz-x ligations on solid phase.

| Ligation Round | Peptide | MW (g/mol) | Peptide (mg) | Volume (mL) | Conc. (mM) | Molar Equivalent | Time (h) |
|---|---|---|---|---|---|---|---|
| 1 | H1.2-(Thz$_{67}$-Gly$_{99}$)-Dbz-GK(C) | 4071 | 5.0 | 1.0 | 1.2 | | 16 |
| 2 | H1.2-(Thz$_{49}$-Ala$_{66}$)-R53cit-MeNbz | 2018 | 8.9 | 1.0 | 4.4 | 7.3 | 5 |
| 3 | H1.2-(Thz$_{24}$-Ala$_{48}$)-MeNbz | 2652 | 9.0 | 0.8 | 4.2 | 5.7 | 5 |
| 4 | H1.2-(Ser$_1$-Ala$_{23}$)-Dbz-R | 2492 | 9.0 | 0.8 | 4.2 | 5.7 | 14 |

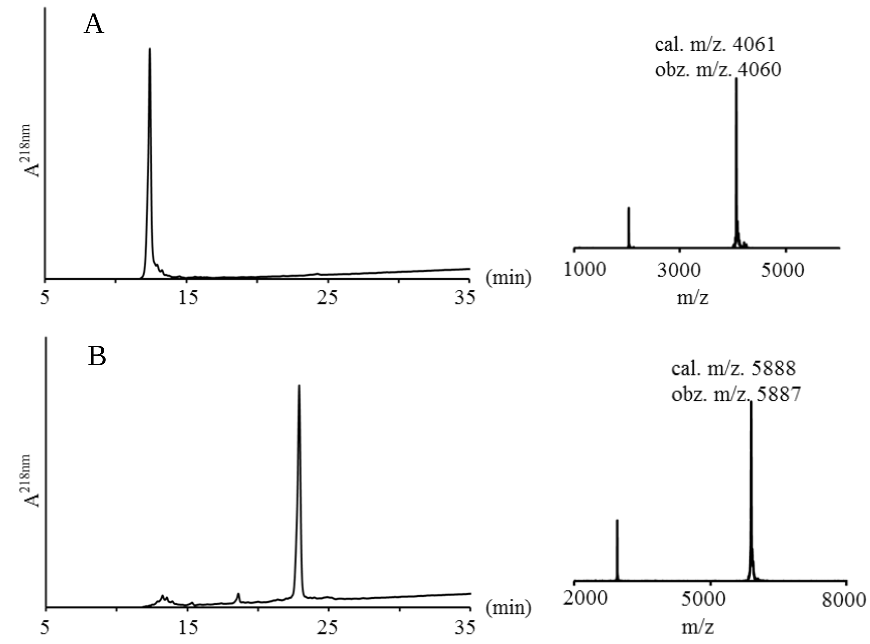

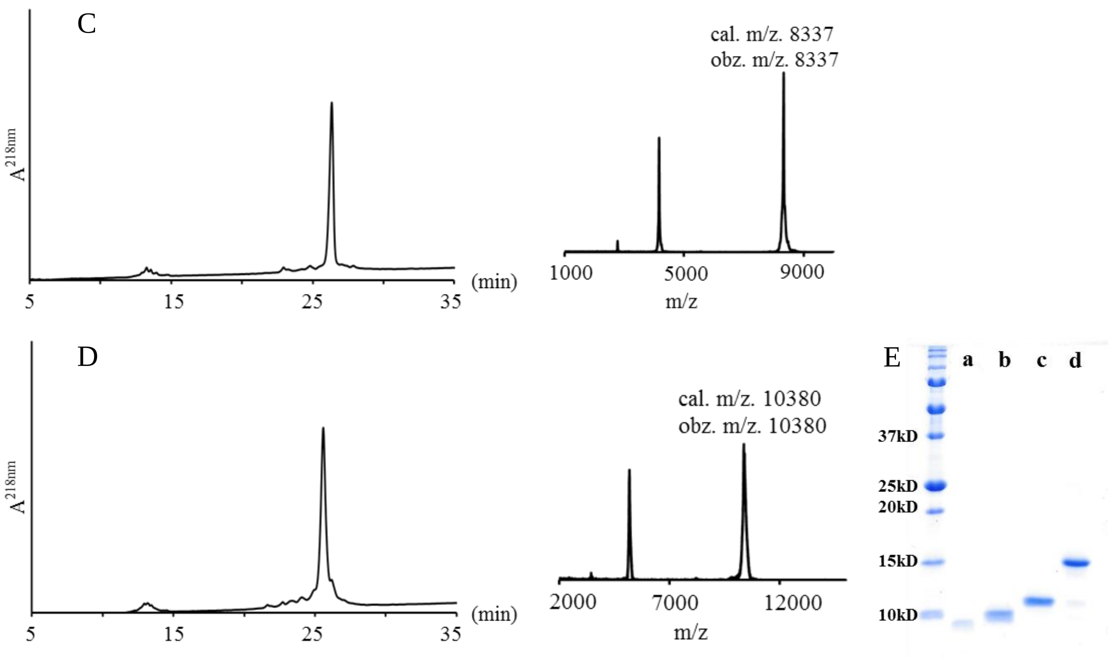

**Figure S34**: Crude RP-HPLC (left panel), MALDI-TOF MS (middle panel), and SDS-PAGE (right panel) of solid phase ligation products. A: H1.2-($Cys_{67}$-$Gly_{99}$)-Dbz-x. B: H1.2-($Thz_{49}$-$Gly_{99}$)-Dbz-x C: H1.2-( $Thz_{24}$-$Gly_{99}$)-Dbz-x. D: after desulfurization H1.2-($Ser_{1}$-$Gly_{99}$)-Dbz-x'. E: a-A, b-B, c-C, d-D. x=GK (C-diclycolic acid-Gly-$NH_2$), x'=GK (A-diclycolic acid-Gly-$NH_2$)

#### 5.15 Peptidyl Thioester Adduct Formation and Elimination during Ligation

We sometimes observed an extra peak during solid phase native chemical ligation. As described previously, this might be caused by reaction between the internal Cys side chain and the excess of peptidyl thioester (Scheme S3). This side product might cause some problems in following ligation process although we didn't observe any in our case. Therefore, developed a protocol to eliminate this side product before moving to next step to avoid any possible problem. We hypothesized that treatment with free Cysteine could be used to remove the side chain adduct due to the reversibility of the transthioesterification, but irreversibility of the reaction of Cys with the adduct. As expected, all side product peaks disappeared after about 30min Cys ligation buffer (6M guanidine, 0.1M phosphate, 0.05M MPAA, 0.05M Cysteine, 10mM TCEP) treatment (Figure S35).

Based on these results, we recommend an extra Cys wash step after each solid phase ligation.

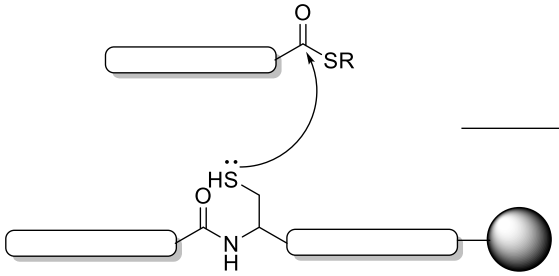

**Scheme S3**: Formation of peptidyl thioester on the side chain of internal Cysteine during ligation.

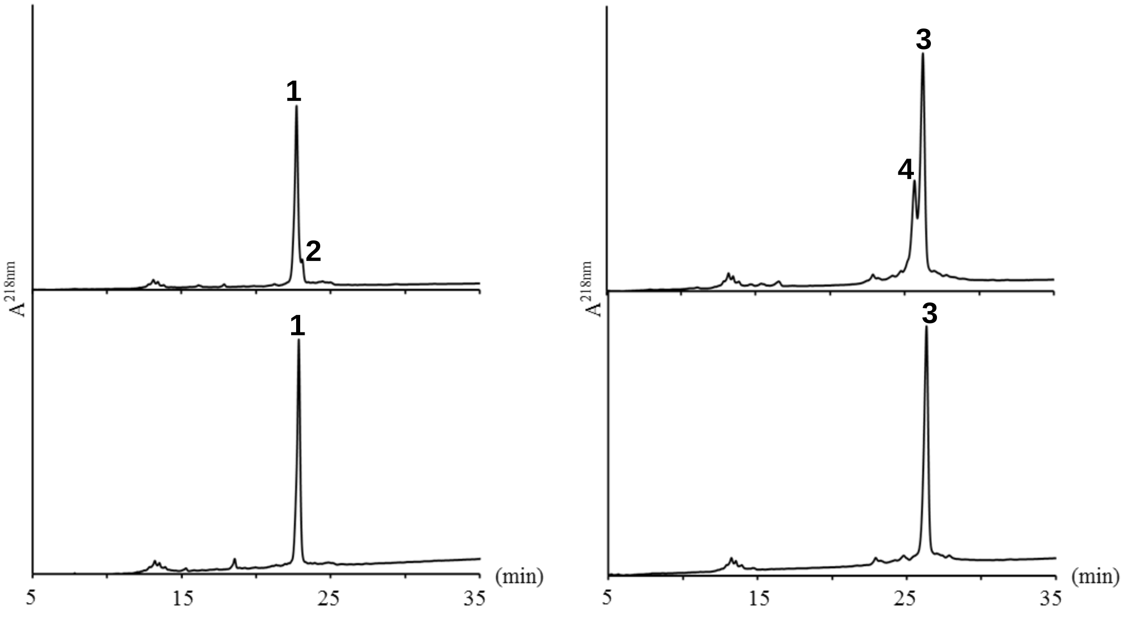

**Figure S35**: Removal of internal Cys side chain peptidyl thioester by short (30min) Cys ligation buffer (6M guanidine, 0.1M phosphate, 0.05M MPAA, 0.05M Cysteine, 10mM TCEP) wash. Top: Before Cys wash. Bottom: After Cys wash. **1**= H1.2-(Thz$_{49}$-Gly$_{99}$)-R53cit-Dbz-x, **2**= H1.2-(Thz$_{49}$-Gly$_{99}$)-R53cit-Dbz-x (H1.2-(Thz$_{49}$-Ala$_{66}$)-R53cit), **3**= H1.2-(Thz$_{24}$-Gly$_{99}$)-R53cit-Dbz-x, **4**= H1.2-(Thz$_{24}$-Gly$_{99}$)-R53cit-Dbz-x (H1.2-(Thz$_{24}$-Ala$_{48}$)).

### 6 H1.2 synthesis by Solution Phase Ligation

In this section, we discuss our optimization of the final solution phase ligation and the following purification for the full-length protein.

**Section 6.1** describes optimization of ligation conditions with model peptides.
**Section 6.2** describes our first full ligation and purification protocols.
**Sections 6.3—6.4** describe two additional ligation instances to generate modified H1.2-R53C and H1.2-S172ph.

##### 6.1 Ligation condition test with model peptides

The final solution phase ligation step requires activation of the N-terminal assembled fragment for ligation with the C-terminal assembled fragment. In initial ligation attempts, we observed significant amount of hydrolyzed H1.2(Ser$_1$-Gly$_{99}$)-OH which reduced our ligation yields. In order to optimize ligation conditions with less valuable materials, we synthesized a model peptide H1.2-(Lys$_{89}$-Gly$_{99}$)-Dbz-Gly-NH$_2$.

1.0mg H1.2-(Lys$_{89}$-Gly$_{99}$)-Dbz-Gly-NH$_2$ (0.8 µmol) was dissolved in 200 µL acidified guanidine buffer (6M guanidine, 0.2M phosphate, pH 3-4) and prechilled in ice/salt bath (-15 $^o$C). Add 20 µL 0.2M $NaNO_2$ (4 µmol, 5 equivalents) to the peptide and incubate the mixture in ice/salt bath for 20min. Add 200µL MPAA buffer (6M guanidine, 0.2M phosphate, 0.1M MPAA) and split the mixture into two parts, with pH adjusted to 6.0-6.5 or 7.8-8.1 (assessed by drop onto pH paper). The hydrolysis process was monitored by RP-HPLC.

We found that peptides under both conditions are converted into peptidyl thioester (**1**) 10min after addition of MPAA buffer and pH change. Not surprisingly, we found that a significant amount of hydrolyzed product (**2**) appears after 10min at pH 7.8-8.1 while there is no obvious hydrolyzed product at pH 6.0-6.5 (Figure S35). All peptidyl thioester was hydrolyzed after 6h at pH 7.8-8.1 while no significant amount of hydrolyzed product appears after 6h at pH 6.0-6.5.

To further validate the function of the thioester at pH 6.0-6.5, we added 2.1mg Cys peptide H3-(Cys$_{47}$-Leu$_{74}$)-Dbz (**3**, 0.8µmol, 2 equivalents) to the peptidyl thioester at pH 6.0-6.5 and found all peptidyl thioester is ligated to the Cys peptide (**4**) after 2h incubation (Figure S36). We therefore carried out further solution phase ligations of the activated Dbz at pH 6-6.5.

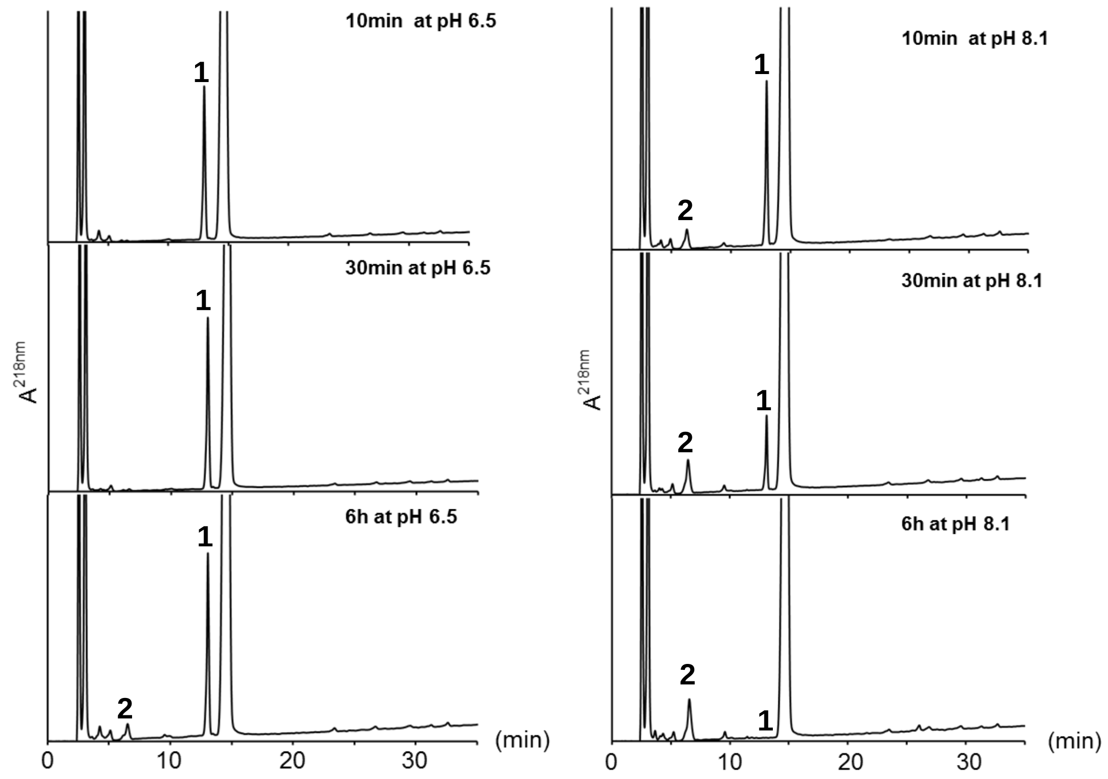

**Figure S36**: RP-HPLC monitoring of the hydrolysis process at different pH conditions. **1**= H1.2-(Lys$_{89}$-Gly$_{99}$)-MPAA, **2**= H1.2-(Lys$_{89}$-Gly$_{99}$)-OH

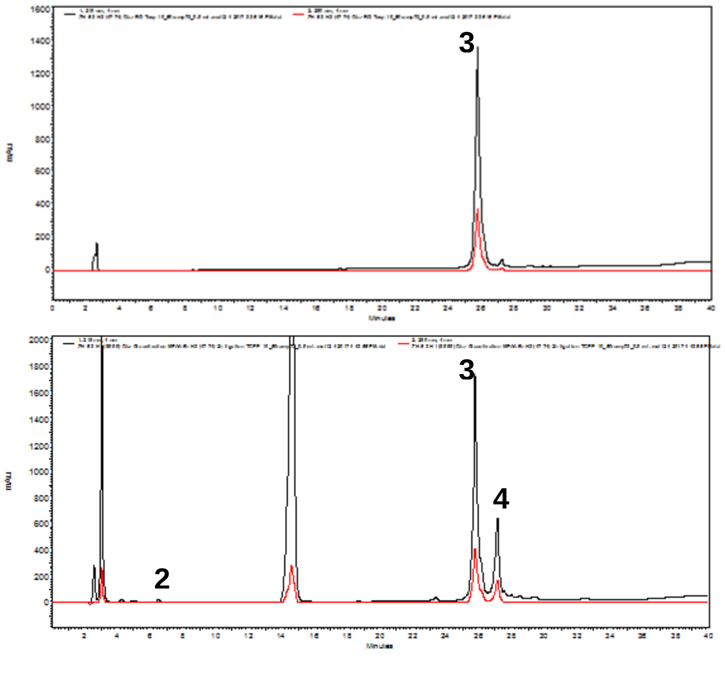

**Figure S37**: RP-HPLC monitoring of the ligation between peptide thioester H1.2-(Lys$_{89}$-Gly$_{99}$)-MPAA (**1**) and Cysteine peptide H3-(Cys$_{47}$-Leu$_{74}$)-Dbz (**3**) at pH 6.0-6.5. Top: H3-(Cys$_{47}$-Leu$_{74}$)-Dbz peptide only. Bottom: 2 hours after incubation. **4**= H1.2-(Lys$_{89}$-Gly$_{99}$)-H3-(Cys$_{47}$-Leu$_{74}$)-Dbz

**6.2 First trial synthesis of unmodified linker histone H1.2 (H1.2-umd)**

583 µg H1.2-(Ser$_1$-Gly$_{99}$)-Dbz-x' (quantified by SDS-PAGE, 50 nmol) was dissolved in 50 µL acidified guanidine buffer (6M guanidine, 0.1M phosphate, pH 3-4) and prechilled in ice/salt bath (-15 $^oC$). 5 µL 0.4M $NaNO_2$ (2 µmol, 40 equivalents) was added and incubated in ice/salt bath for 20min. 2.0mg H1.2-(Cys$_{100}$-Lys$_{212}$)-OH (200 nmol, 4 equivalents) was dissolved in 50 µL MPAA buffer (6M guanidine, 0.1M phosphate, 0.1M MPAA, pH 7) and added to the above $NaNO_2$-activated peptide solution. The reaction process was monitored by SDS-PAGE (Figure S38A). The ligation was almost complete in the 7 hours viewed by the gel. The ligation mixture was dialyzed against guanidine buffer (6M guanidine, 0.1M phosphate, pH 7) three times to remove all MPAA. Desulfurization was carried out under the conditions described in S5.5 and assessed by MALDI-TOF MS (Figure S38B).

The final protein was purified by RP-HPLC with a gradient of 30~50% buffer B over 50min on a C5 column and the collected fractions were assessed by SDS-PAGE and MALDI-TOF MS (Figure S38C-D) before lyophilization. The analysis loss in the was ~10%. About 140 µg of pure H1.2 protein was obtained in a yield of ~13% (excludes analysis loss) after ligation-dialysis-desulfurization and RP-HPLC purification, quantified by in-gel comparison against commercial H1.0 standard (NEB) (Fig. S38C).

Significantly, purification of the full-length product was complicated due to very close retention time of the full-length H1.2 and N-terminal segment H1-(Ser$_1$-Gly$_{99}$)-Dbz-x on RP-HPLC. To compensate for this and minimize overlap, we typically include excess C-terminal segment in the ligation – and all test ligations proceeded efficiently to near completion assessed on this N-terminal segment. However, the recovery yield after RP-HPLC purification remains relatively low (10-13%). This is consistent with previous results on core histones carried out by our group and other labs. We assume that this yield loss may result from non-specific interaction between histone proteins and RP-HPLC columns. This highlights the significance of combining solid phase ligation and convergent ligation to reduce or even avoid RP-HPLC purification steps for large protein preparation.

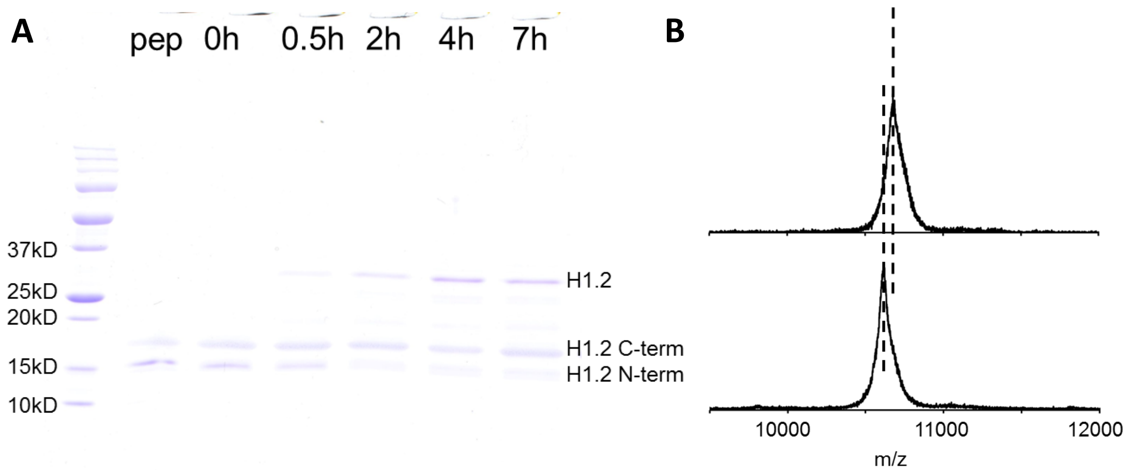

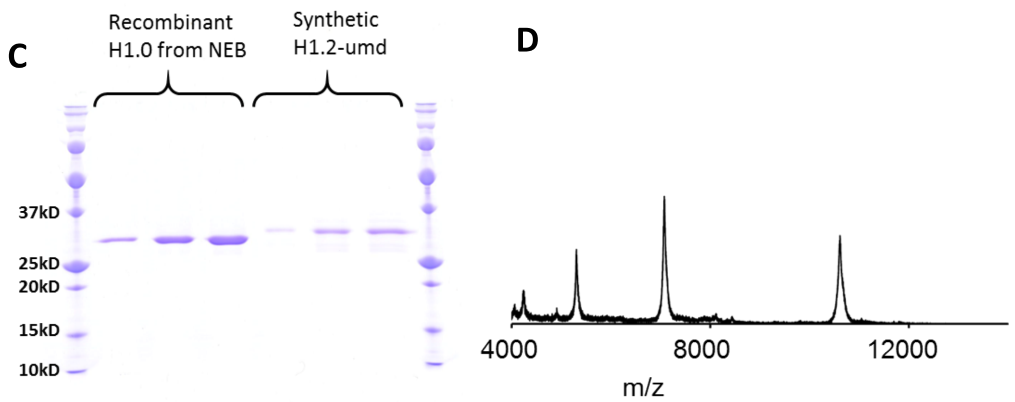

**Figure S38**: Synthesis of H1.2-R53cit accessed by SDS-PAGE and MALDI-TOF MS. A: timepoint monitoring of the ligation process. B: MS difference before and after desulfurization. Top: before desulfurization $[M+2H]^{2+}$ expected m/z:10682. observed m/z: 10679. Bottom: after desulfurization $[M+2H]^{2+}$ expected m/z: 10618. observed m/z: 10617. C: SDS-PAGE quantification of final purified product. D: MALDI-TOF MS of final purified product. $[M+2H]^{2+}$ expected m/z:10618. observed m/z: 10617. $[M+3H]^{3+}$ expected m/z:7079. observed m/z: 7076.

##### 6.3 Synthesis of citrullinated linker histone H1.2 (H1.2-R53cit)

In our first trial of preparing citrullinated linker histone H1.2, 1.1 mg H1.2-($Ser_1$-$Gly_{99}$)-R53cit-Dbz-x' (quantified by SDS-PAGE, 100 nmol) was dissolved in 50 µL acidified guanidine buffer (6M guanidine, 0.1M phosphate, pH 3-4) and prechilled in ice/salt bath (-15 °C). Add 5 µL 0.2M $NaNO_2$ (1 µmol, 10 equivalents) to the peptide and incubate the mixture in ice/salt bath for 20min. 2.0mg H1.2-($Cys_{100}$-$Lys_{212}$)-OH (by weight, 200 nmol, 2 equivalents) was dissolved in 50 µL MPAA buffer (6M guanidine, 0.1M phosphate, 0.1M MPAA, pH 7) and added to the above $NaNO_2$-activated peptide solution. The reaction process was monitored by RP-HPLC or SDS-PAGE. The ligation is almost complete after 4h as shown on SDS-PAGE and RP-HPLC (Figure 5B &D in main paper). The ligation mixture was dialyzed against guanidine buffer (6M guanidine, 0.1M phosphate, pH 7) three times until all MPAA was removed. Desulfurization was carried out as described in S5.5 and assessed for completion by MALDI-TOF MS. The final protein is purified by RP-HPLC at a gradient from 30-45% buffer B on a C5 column. The pooled product was assessed by SDS-PAGE and MS (Figure 5D, E &F in main paper) and quantified by in-gel comparison to commercial H1.0 standard (NEB). 55% of the sample was taken for analysis over the full process. After accounting for this 55% (ie: calculated from 45nmol of limiting N-terminal segment), we obtained ~110 µg pure H1.2-R53cit protein for a yield of ~11%.

**6.4 Synthesis of citrullinated linker histone H1.2 (H1.2-R53cit) using MESNa**

The use of MPAA as the ligation co-thiol requires thorough dialysis steps between ligation and desulfurization, to remove the aromatic thiol that quenches the free radical desulfurization reaction. Some yield loss in unavoidable during the dialysis process. For ligations carried out with the non-aromatic mercaptoethanesulfonate (MESNa) co-thiols, desulfurization can be carried out directly on the ligation mixture – however, ligation tends to proceed more slowly, which can lead to greater loss through hydrolysis of the thioester prior to ligation. To assess the balance between these different potential sources of yield loss, we explored the use of MESNa as the thiol additive during the last solution phase ligation to generate H1.2-R53cit.

2.5mg of peptide H1.2-($Ser_1$-$Gly_{99}$)-R53cit-Dbz-x' (by weight, 240nmol) was dissolved in 100μL acidified guanidine buffer (6M guanidine, 0.1M phosphate, pH 3-4) and prechilled in ice/salt bath (-15 $^{o}$C). 5 μL 0.2M $NaNO_2$ (1 μmol, 4 equivalents) was added to the peptide mixture and incubated in ice/salt bath for 20min. 5.6mg H1.2-($Cys_{100}$-$Lys_{212}$)-OH (by weight, 490 nmol, 2 equivalents) was dissolved in 200 μL MESNa buffer (6M guanidine, 0.1M phosphate, 0.2M MESNa, pH 7) and added to the above $NaNO_2$-activated peptide solution. The pH was adjusted to ~6.5 by addition of NaOH. However, minimal to no ligation product was observed by SDS-PAGE analysis in the first 30min (Figure. S39A) or RP-HPLC (data not shown). We allowed the reaction to continue through 2h, but still observed no ligation product (data not shown).

At this point, we determined MESNa co-thiol was substandard in this context and, to rescue the synthetic materials to proceed to product, added MPAA to a final concentration of 50mM and adjusted the pH to ~6.5. This did initiate the ligation reaction, but unfortunately only after some hydrolysis of starting material, as assessed by the incomplete ligation observed after 44h (Figure. S39A).

The ligation mixture was then dialyzed against guanidine buffer (6M guanidine, 0.1M phosphate, pH 7) three times until all MPAA was removed. Desulfurization was carried out with the same protocol as for solid phase and assessed by MALDI-TOF MS (Figure S39B). The final protein was purified by RP-HPLC at gradient 30-50% buffer B over 50min on C5 column. The pure pool protein was assessed by SDS-PAGE and MALDI-TOF MS (Figure S39C &D). ~20% of the sample was taken for analysis over the full ligation/desulfurization process. Finally, 423 μg pure protein H1.2-R53cit was obtained in a yield of 10% (excludes analysis loss).

We do not recommend the use of MESNa as a thiol additive in our H1.2 case based on our results here. Although the ligation did not go to completion as observed in other trials, the yield was comparable. We hypothesize that this is due to lower loss during purification due to the larger scale of the ligation reaction.

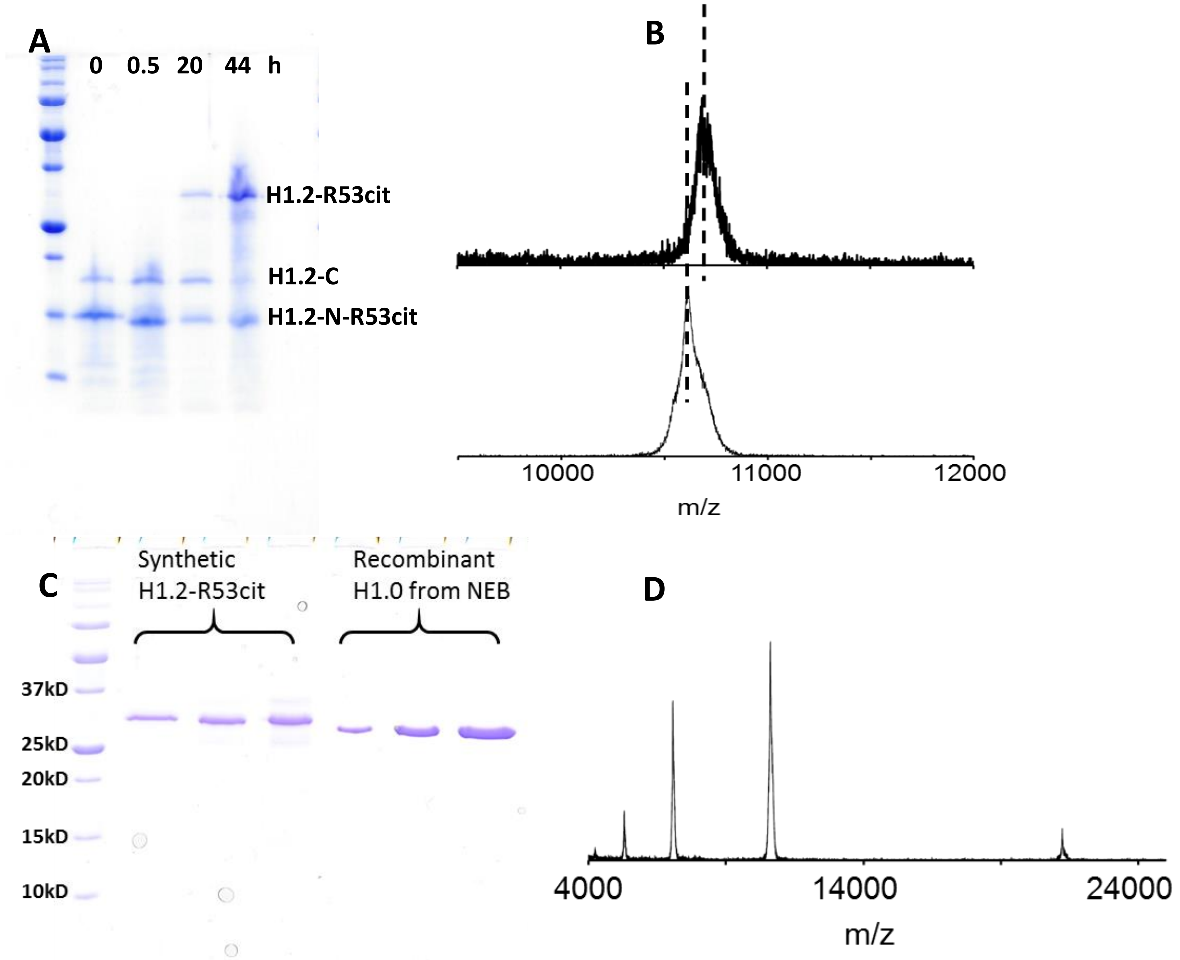

**Figure S39**: Synthesis of H1.2-R53cit accessed by SDS-PAGE and MALDI-TOF MS. A: timepoint monitoring of the ligation process at 0, .5, 20, and 44h. B: MS difference before and after desulfurization. Top: before desulfurization $[M+2H]^{2+}$ expected m/z:10682. observed m/z: 10682. Bottom: after desulfurization $[M+2H]^{2+}$ expected m/z: 10618. observed m/z: 10617. C: SDS-PAGE quantification of final purified product. D: MALDI-TOF MS of final purified product. $[M+H]^{+}$ expected m/z:21236. observed m/z: 21236. $[M+2H]^{2+}$ expected m/z:10618. observed m/z: 10617. $[M+3H]^{3+}$ expected m/z:7079. observed m/z: 7072.

##### 6.5 Synthesis of phosphorylated linker histone H1.2 (H1.2-S172ph)

0.9 mg µg H1.2-($Ser_1$-$Gly_{99}$)-Dbz-x' (by weight, 85 nmol) and 1.4 mg H1.2-($Cys_{100}$-$Lys_{212}$)-S172ph-OH (by weight, 120 nmol) were dissolved in 75 µL acidified guanidine buffer (6M guanidine, 0.1M phosphate, pH 3-4) and prechilled in ice/salt bath (-15 °C). Add 10 µL 0.2M $NaNO_2$ (2 µmol, 24 equivalents) to the peptide solution and incubate the mixture in ice/salt bath for 20min. 10 µL MPAA buffer (6M guanidine, 0.1M phosphate, 0.5M MPAA, pH 8) and added to the above $NaNO_2$-activated peptide solution. The reaction process was monitored by RP-HPLC or SDS-PAGE (Figure S40A). The ligation mixture was dialysis against guanidine buffer (6M guanidine, 0.1M phosphate, pH 7) several times until all MPAA are removed. Desulfurization was carried out with the same condition as on solid phase. Fresh free-radical initiator VA-044US will be added until desulfurization is complete (Figure S40B). The final protein is purified by RP-HPLC at gradient from 30-50% buffer B over 50min on C5 column and assessed by SDS-PAGE and MALDI-TOF MS (Figure S40C-D). The analysis loss in the whole process is about 55%. ~60 µg pure H1.2-S172ph protein was obtained in a yield of ~8% (excludes analysis loss) after ligation-dialysis-desulfurization and RP-HPLC purification.

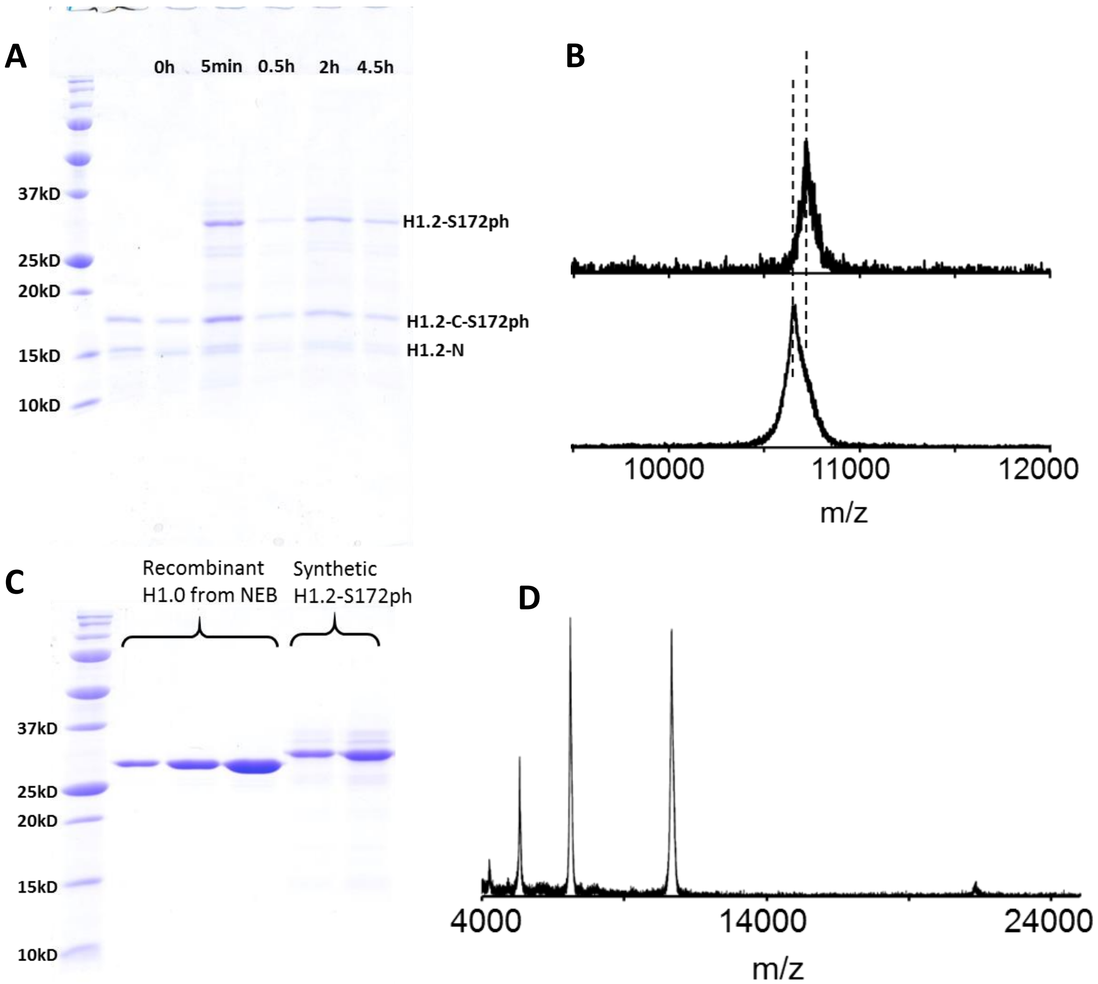

**Figure S40**: Synthesis of H1.2-S172ph accessed by SDS-PAGE and MALDI-TOF MS. A: timepoint monitoring of the ligation process. B: MS difference before and after desulfurization. Top: before desulfurization $[M+2H]^{2+}$ expected m/z:10721. observed m/z: 10722. Bottom: after desulfurization $[M+2H]^{2+}$ expected m/z: 10657. observed m/z: 10657. C: SDS-PAGE quantification of final purified product. D: MALDI-TOF MS of final purified product. $[M+2H]^{2+}$ expected m/z:10657. observed m/z: 10657. $[M+3H]^{3+}$ expected m/z:7106. observed m/z: 7103. $[M+4H]^{4+}$ expected m/z:5329. observed m/z: 5326.

### 7 Electrophoretic mobility shift assay to assess H1 binding

##### 7.1 Preparation of DNA molecules

DNA molecules containing the Widom 601 positioning sequence[10] with an additional 50 bp on the 5' end and 75 bp on the 3' end of the sequence and a Gal4-2C binding site (5'-CCGGAGGGCTGCCCTCCGG-3')[11] were produced by PCR using Cy3/biotin labeled oligonucleotides and 601 sequence containing plasmid. The final DNA construct is 272 bp long and contains Cy3 8 bp away from the 5' end of the 601 sequence, the Gal4-2C binding site at bases 8-26 of the 601 sequence, and a biotin at the 3' end of the construct. Oligonucleotides were labeled with Cy3 NHS ester (GE Healthcare) at an amino modified thymine and HPLC purified with a 218TP C18 column (Grace/Vydac). After PCR amplification, DNA constructs were HPLC purified with a MonoQ 5/50 GL anion exchange column (GE Healthcare).

##### 7.2 Preparation of core histones and histone octamer

Core histones and histone octamer were produced as previously described[12] with some modifications. Briefly, human H2A(K119C), H2B, H3(C110A), and H4 were expressed in Rosetta or BL21 (DE3) pLysS cells. Core histones were purified in denaturing conditions using size exclusion chromatography followed by cation exchange chromatography. Histone octamer was formed by resuspending lyophilized core histones in unfolding buffer (20 mM Tris-HCl pH 7.5, 7 M guanidinium, 10 mM DTT) at 5 mg/mL and mixing in a ratio of H2A:H2B:H3:H4 of 1.1:1.1:1:1. Histones were refolded and octamer produced by performing double dialysis into refolding buffer (10 mM Tris-HCl pH 7.5, 1 mM EDTA, 2 M NaCl, 5 mM BME). Histone octamer was labeled with Cy5 maleimide (GE Healthcare) as described previously[13] with addition of 10 fold molar excess dye dissolved in DMF at 22 mM reacted at room temperature on a spinning rotisserie for 1 hour then overnight at 4 °C. Labeling was quenched with 10 mM DTT and octamer was purified with a Superdex 200 size exclusion column (GE Healthcare) to remove free histones, heterodimer, and excess dye.

##### 7.3 Preparation of nucleosomes

Nucleosomes were prepared, as done previously[12,13], by mixing DNA and octamer in a molar ratio of 1.25:1 (DNA:octamer) in 0.5x TE pH 8.0, 2 M NaCl, 1 mM benzamidine-HCl in 50-100 µL and reconstituted by salt double dialysis into 0.5x TE pH 8.0, 1 mM benzamidine-HCl. Nucleosomes were added to the top of a 5%-30% w/v sucrose gradient and purified with an Optima L-90K Ultracentrifuge (Beckman Coulter) with a SW41 rotor spinning at 41,000 rpm for 22 hours at 4 °C. Sucrose gradients were fractionated into 0.4 mL fractions and those containing correctly positioned nucleosomes were concentrated and buffer exchanged into 0.5x TE pH 8.0.

##### 7.4 H1 concentration determination

The concentration of H1 used for *in vitro* assays was determined with Coomassie stained SDS-PAGE gels and a modified Lowry assay[14] (Pierce). For Coomassie stained gels, a range of H1.0 (NEB) of known concentration, determined by absorbance at 280 nm by the manufacturer, was loaded onto the gel with the unknown H1. The intensity of stained bands was determined by ImageJ and a standard curve was made with the H1.0 and used to quantify unknown H1 concentrations. Separately, concentration was determined with a modified Lowry assay following the manufacturer's instructions using H1.0 to produce a standard curve and calculate the unknown H1 concentration.

##### 7.5 Electrophoretic mobility shift assay

EMSAs were performed with 1 nM nucleosomes incubated with a range of H1 (0-300 nM) in 10 mM Tris-HCl pH 8.0, 130 mM NaCl, 10% Glycerol, 0.005% TWEEN20 in a 20 uL volume at room temperature for 20 min. Reactions were ran on a 4% polyacrylamide, 0.3x TTE, 10% glycerol gel at 4 °C for 2 hours and imaged on a Typhoon imager (GE Healthcare).
